# HEARTSVG: a fast and accurate method for spatially variable gene identification in large-scale spatial transcriptomic data

**DOI:** 10.1101/2023.08.06.552154

**Authors:** Xin Yuan, Yanran Ma, Ruitian Gao, Shuya Cui, Yifan Wang, Botao Fa, Shiyang Ma, Ting Wei, Shuangge Ma, Zhangsheng Yu

## Abstract

Identifying spatially variable genes (SVGs) is crucial for understanding the spatiotemporal characteristics of diseases and tissue structures, posing a distinctive challenge in spatial transcriptomics research. We propose HEARTSVG, a distribution-free, test-based method for fast and accurately identifying spatially variable genes in large-scale spatial transcriptomic data. Extensive simulations demonstrate that HEARTSVG outperforms state-of-the-art methods with higher *F*_1_ scores (average *F*_1_ score=0.903), improved computational efficiency, scalability, and reduced false positives (FPs). Through analysis of twelve real datasets from various spatial transcriptomic technologies, HEARTSVG identifies a greater number of biologically significant SVGs (average recall=0.985, average AUC=0.788) than other comparative methods without prespecifing spatial patterns. Furthermore, by clustering SVGs, we uncover two distinct tumor spatial domains characterized by unique spatial expression patterns, spatial-temporal locations, and biological functions in human colorectal cancer data, unraveling the complexity of tumors.

## 1. Introduction

Spatial transcriptomics enables the measurement of gene expression and positional information in tissues^1–6^. The evolution of spatial transcriptomics technologies advanced the reconstruction of tissue structure and provided profound insights into developmental biology, physiology, cancer, and other fields^2,7,8^. However, the complexity and high dimensionality of spatial transcriptomics (ST) data pose new challenges and requirements for analytical approaches^8-9^. One crucial analytical challenge in spatial transcriptomics studies is the identification of spatially variable genes (SVGs) whose expressions correlate with spatial location^7,10,11^, also known as SE genes (genes with spatial expression patterns)^12^. Identifying SVGs promotes characterizing spatial patterns within tissues and predicting spatial domains^7,10,13,14^. Several methods have been developed for detecting SVGs. Trendsceek^15^ models the data as marked point processes and tests the significant dependency between spatial distributions and expression levels of pairwise points. SpatialDE^11^ decomposes gene expression variability into a spatial component and an independent noise term based on Gaussian process regression and tests statistical significance by comparing the SpatialDE model to a null model without the spatial variance component. SPARK^16^, an extension of SpatialDE, uses the Gaussian process regression as the underlying data model and ten different spatial kernels to represent common spatial patterns in biological data, thereby improving statistical power. SPARK-X^12^ tests the dependence of gene expressions and spatial locations based on the covariance test framework. scGCO^17^ applies graph cuts in computer vision to address SVG identification. It utilizes the hidden Markov random field to identify candidate regions with spatial dependence for individual genes and tests their dependence under the complete spatial randomness framework.

Trendsceek, SpatialDE, and SPARK have limited applicability for large-scale datasets due to their high computational complexity. Trendsceek employs the permutation strategy to compute multiple statistics of different paired points, which requires extensive computational work and is only scalable to small-scale datasets. The Gaussian process framework hinders the detection of SVGs and model parameter convergence in SpatialDE and SPARK when analyzing high-dimensional and sparse ST data. SPARK-X offers significantly faster computational speed than the aforementioned methods, but its effectiveness depends heavily on how well the constructed spatial covariance matrix matches the true underlying spatial patterns. The above four methods identify SVGs by searching for predefined relationships between expressions and locations. They have limited generalizability to a wide range of spatial patterns due to the arbitrary nature of the true spatial pattern of SVGs and the resulting uncertainty in the relationship between expression and coordinates. scGCO has the capability to identify SVGs with unknown exact locations and shapes, however, it suffers from false negatives due to the limited accuracy of the graph cuts algorithm in identifying candidate regions for SVGs, especially in sparse ST datasets.

Hence, we propose HEARTSVG to overcome the limitations without prior knowledge or specification of information about SVGs. We take an opposite approach by identifying non-SVGs and using this information to infer the presence of SVGs. Although the relationship between gene expression and spatial position of an SVG is uncertain, it is unequivocal that a non-SVG has “no relationship” between gene expression and spatial position. HEARTSVG identifies non-SVGs by testing the serial autocorrelations in the marginal expressions across global space. By excluding non-SVGs, the remaining genes are considered as SVGs. As a test-based method without assuming underlying spatial patterns, HEARTSVG detects SVGs with arbitrary spatial expression shapes and is suitable for diverse types of large-scale ST data. We conduct extensive simulations and apply HEARTSVG method to twelve real ST datasets generated from different technologies (including 10X Visium, Slide-seqV2, MERFISH, and HDST) to demonstrate its accuracy, robustness, and computational efficiency. HEARTSVG outperforms existing methods in simulations with higher accuracy metrics, computational efficiency, and lower false positives (FPs). When analyzing real ST data, HEARTSVG identifies biologically meaningful SVGs with distinct spatial expression patterns across diverse datasets obtained from different spatial transcriptomic technologies. HEARTSVG has the potential to scale to datasets comprising millions of data points and offers a comprehensive range of meticulously designed analytical tools for studying SVGs, enabling the unraveling of complex biological phenomena.

### 2.1 Overview of HEARTSVG

HEARTSVG aims to identify SVGs that display spatial expression patterns in spatial transcriptomics data. Each gene in the ST data is presented as a vector containing three elements: gene *g* = (*x*, *y*, *e*)*^T^*, where *x* and *y* are defined as the row and column positions of the spot, respectively, and *e* is the gene expression count of the gene *g* at the spatial coordinates (*x*, *y*). HEARTSVG is based on the intuitive concept that the non-SVG *g* does not display a spatial expression pattern, its expression distribution is expected to be independent and random, with marginal expression distributions along the *x*-axis (row) and *y*-axis (column) also being independent and random. Conversely, suppose the *g* exhibits a spatial expression pattern, both its spatial expression and marginal expression should have serial correlation along the single direction (row or column) or both. Therefore, a non-SVG demonstrates low autocorrelations, while an SVG has high autocorrelations (Fig.**1a**, Derivations and more details are provided in the “Methods” section and Supplementary 1).

**Fig.1.**
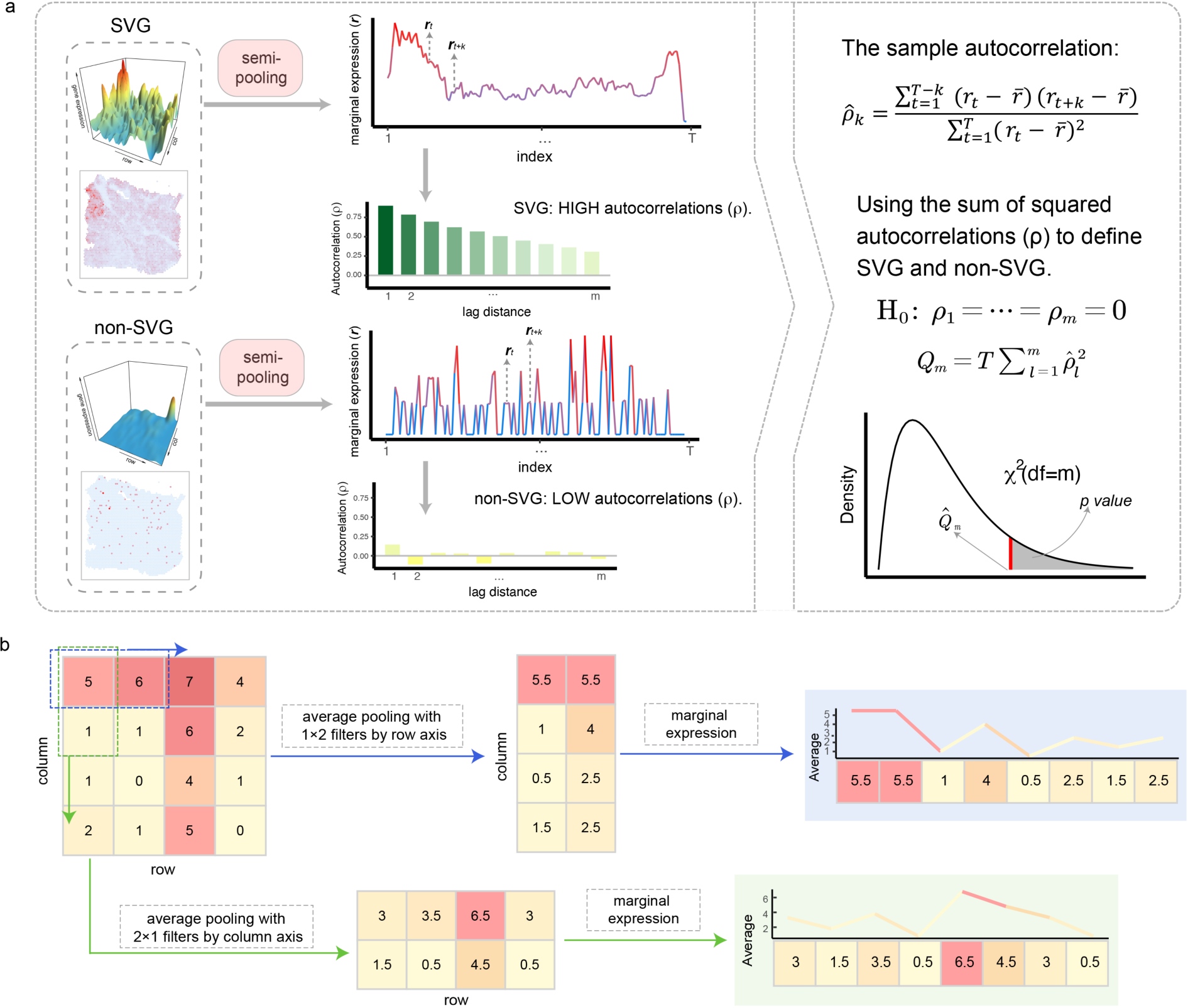
Schematic representation of the HEARTSVG. **a,** HEARTSVG utilizes the semi-pooling process to convert the spatial expression of genes into marginal expressions (*r*) and calculates autocorrelations (*ρ*) of marginal expressions. The autocorrelations (*ρ*) of marginal expressions (*r*) for the SVG and non-SVG exhibit different level scales. Representative autocorrelation estimator plots are plotted below the corresponding marginal expression plots for the SVG and non-SVG. The color depth represents the magnitude of autocorrelation. HEARTSVG distinguishes between SVGs and non-SVGs by summarizing the sum of the squared autocorrelations 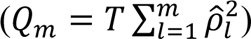 as shown in the right panel of Fig.**1a**. **b,** Illustration of the semi-pooling process along the row (feature map: 1 × 2), and column (feature map: 2 × 1) directions.

HEARTSVG uses the semi-pooling process to transform the gene’s two-dimensional spatial expression to one-dimensional marginal expression serials along the single direction (row or column) (Fig.**1b**, more details are provided in the “Methods” section and Supplementary 1). This process aims to extract information and reduce data noise and sparsity from gene spatial expression data. The Portmanteau test^18,19^ is then performed to test serial autocorrelations of the gene’s marginal expression series. The non-SVG’s marginal expressions exhibit constant variance, zero autocorrelation, and no trend or periodic fluctuations across locations (More details are provided in the “Methods” section and Supplementary 1). Conversely, marginal expressions of SVGs have high autocorrelations (Fig.**1a**). We obtained multiple p-values by conducting the Portmanteau test to evaluate four marginal expression series with different semi-pooling parameters (More details in Supplementary 1). We then combined all four p-values into a single p-value using Stouffer’s method^20,21^. The classification of a gene as an SVG was determined based on this single p-value at an adjusted p-value cutoff of 0.05. In addition, HEARTSVG provides an auto-clustering module for SVGs in the software, which is complementary to SVG detection for further biological investigations. The auto-clustering module (More details in Methods) comprises functionalities for predicting spatial domains, conducting functional studies, and visualization based on SVGs.

### 2.2 Simulation

We conducted extensive simulations to evaluate the performance of HEARTSVG and compare it with four other methods: SpatialDE, SPARK, SPARK-X and scGCO. Simulation data were generated with three representative SVG spatial expression patterns: hotspot, streak, and gradient (Fig.**2a**). Hotspot and streak spatial patterns had small pattern sizes (5% of the marked area), while the gradient spatial pattern had a large pattern size (15% of the marked area). Spot coordinates of each spot were generated using a Poisson random point process and gene expression counts were generated from the zero-inflated negative binomial distribution (ZINB). Each simulation scenario was replicated 50 times. Extensive simulation datasets were generated by varying the sample size, ZINB parameters (mu, size), and SVG percentages (More details in the “Methods” section). The *F*_1_ score was used to assess the performances of identification SVGs of HEARTSVG and three other SVG detection methods in identifying SVGs.

**Fig.2.**
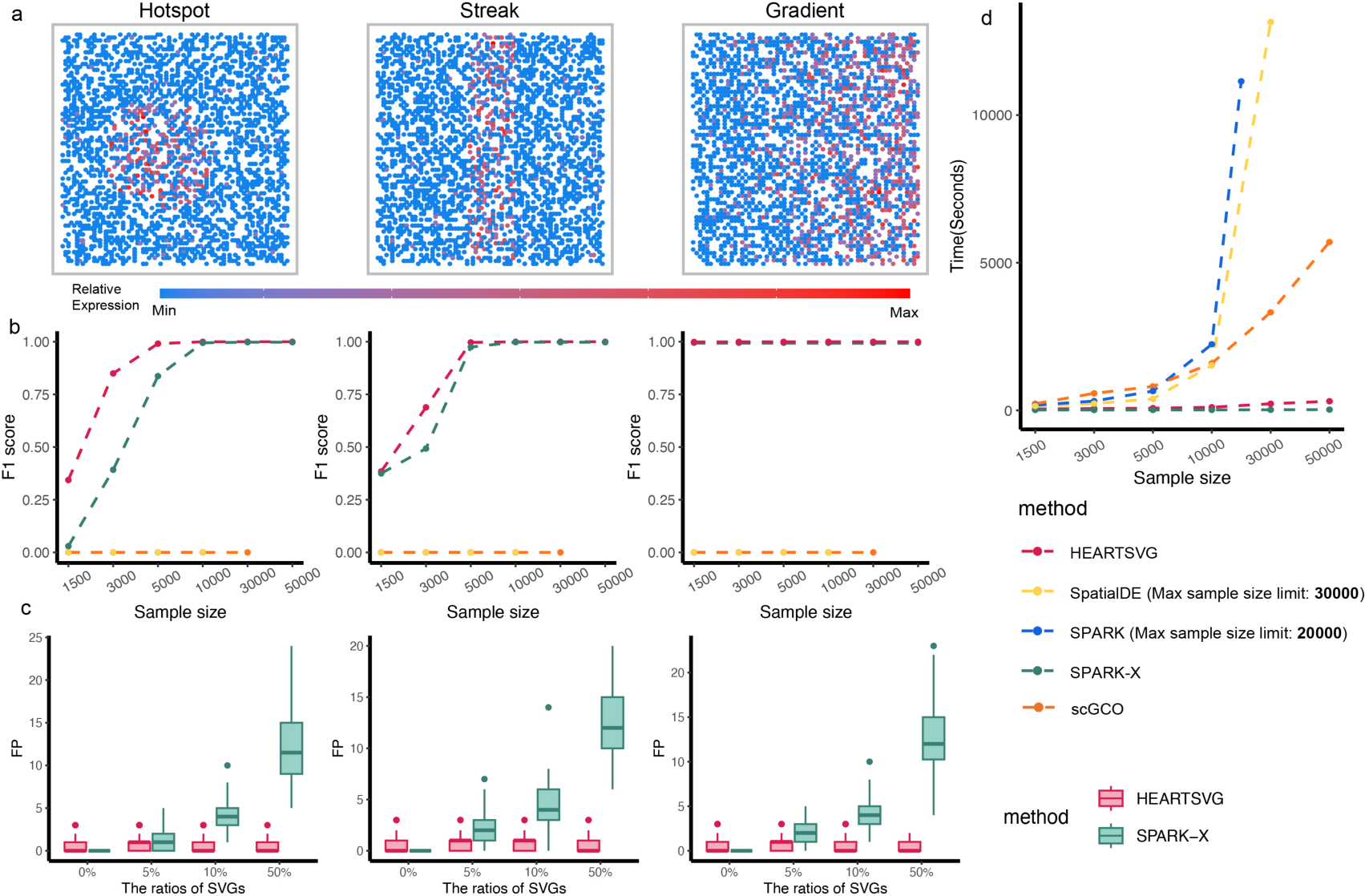
Simulation results show that HEARTSVG has high accuracy and good scalability and computational efficiency. **a,** Visualization of three representative spatial expression patterns: hotspot, streak, and gradient. **b,** *F*_1_ score plots compare the accuracy (y-axis) of HEARTSVG (red), SpatialDE (yellow), SPARK (blue), SPARK-X (green), and scGCO (orange) in simulation data with varying sample sizes (x-axis) at an adjusted p-value cutoff of 0.05. Each plot corresponds to the upper spatial patterns in Fig.**2a**. Simulations were conducted under high sparsity (*zero* − *inflaation parameter* = 0.941, *size* = 0.5, *mu* = 0.5) and a moderate proportion of SVGs (10%, 1,000 SVGs). The *F*_1_ score for each method in each simulation scenario represents the average of fifty replications. For easy computation, we did not apply SpatialDE and SPARK to datasets with sample sizes of more than 10,000 in these simulations. **c,** False positive (FP, y-axis) boxplots of HEARTSVG (red) and SPARK-X (green) in simulation data with different SVGs ratios (x-axis) at an adjusted p-value cutoff of 0.05. Each scenario has 10,000 genes, including both SVGs and non-SVGs. Each boxplot corresponds to the upper spatial patterns in Fig.**2a**. Simulations were conducted under high sparsity (*zero* − *inflaation parameter* = 0.941, *size* = 0.5, *mu* = 0.5) and moderate sample size (n=5,000). In each boxplot, the lower hinge, upper hinge, and center line represent the 25th percentile (first quartile), 75th percentile (third quartile), and 50th percentile (median value), respectively. Whiskers extend no further than ±1.5 times the inter-quartile range. Data beyond the end of the whiskers are considered outliers and are plotted individually. **d**, Plot shows time consumption in seconds (y-axis) of each method for analyzing 10,000 genes with different sample sizes (x-axis). Considering the limitation of scalability, we did not apply SpatialDE to datasets with sample sizes exceeding 30,000 and did not apply SPARK to datasets with sample sizes exceeding 20,000. Fig.**2b** and Fig.**2d** have the common figure legend.

Simulation results demonstrated that HEARTSVG and SPARK-X achieved high accuracy in identifying SVGs while maintaining superior computational efficiency (Fig.**2b, d****, S1-3**). SpatialDE, SPARK struggled to identify SVGs in sparse data due to their adoption of the Gaussian data-generative model, which is not well-suited for gene spatial expression distribution. scGCO exhibited many false negatives in due to its difficulty in accurately identifying candidate regions for SVGs in highly sparse datasets. On the low sparsity simulated data, scGCO showed improved *F*_1_scores on simulated data with lower sparsity (*F*_1_ score=0.333). However, both HEARTSVG (*F*_1_ score=0.998) and SPARK-X (*F*_1_score=0.997) exhibited superior performance (**Fig. S7**). Identification performance was influenced by the pattern sizes of SVGs and sample sizes (Fig.**2b****, S1-3**). HEARTSVG and SPARK-X excelled in identifying almost all SVGs (*F*_1_ score=0.999, 0.998) in the gradient spatial pattern with a large pattern size. In simulations with smaller pattern sizes and sample sizes, HEARTSVG outperformed SPARK-X in identifying SVGs. For hotspot and streak patterns with a sample size of 3,000 spots, HEARTSVG achieved higher *F*_1_ scores (*F*_1_ scores=0.850, 0.689) than SPARK-X, which achieved only 0.392 and 0.493, respectively. Furthermore, HEARTSVG performed stably across different spatial expression patterns of SVGs, while SPARK-X exhibited pattern preference, being sensitive to streak patterns and insensitive to hotspot patterns due to the assumptions regarding its pre-constructed spatial covariance matrix.

To account for the uncertainty regarding the number of SVGs in real data, we generated additional simulation datasets with varying percentages of SVGs (Fig.**2c**). We specifically compared the performance of HEARTSVG and SPARK-X, which showed better results in the previous simulations. With the exception of the theoretical scenario of 0% SVG proportion, which only exists in simulations, HEARTSVG had lower average false positives compared to SPARK-X. As the proportion of SVGs increased, the false positive (FP) numbers of SPARK-X grew, while HEARTSVG maintains low FPs (less than 5). The *F*_1_ scores of HEARTSVG and SPARK-X were similar, but SPARK-X exhibited large variations of *F*_1_ scores (Fig.**S4-6**). In addition, HEARTSVG demonstrated good scalability and computational performance (Fig.**2d**). HEARTSVG, SPARK-X, and scGCO can scale to datasets exceeding 50,000 spots. However, HEARTSVG and SPARK-X exhibited comparable computational speeds, significantly faster than SpatialDE, SPARK and scGCO. HEARTSVG completed the computation on a 50,000-spot dataset in approximately 5 minutes, while scGCO took 95 minutes. On the other hand, SPARK and SpatialDE were limited to sample sizes of 20,000 and 30,000 spots, respectively. SPARK necessitated over 3 hours for a 20,000-spot dataset, and SpatialDE took nearly 4 hours for a 30,000-spot dataset.

### 2.3 Applications to ST datasets from different spatial technologies

Spatial transcriptomic technologies have various sequencing methods and yield different data characteristics. Therefore, in addition to large-scale simulations, we evaluated the accuracy, robustness, and generality of HEARTSVG on several real ST datasets from different ST technologies, comprising three next-generation sequencing (NGS)-based spatial technologies (10X Visium^5,22–24^, Slide-seqV2^3^ and HDST^4^), and one imaging-based spatial technology (MERFISH^6^).

#### HEARTSVG identifies SVGs and predicts spatial functional domains in the 10X Visium colorectal cancer data

10X Visium is the most widely used commercial spatial transcriptomics technology in cancer research. We applied HEARTSVG to a human colorectal cancer (CRC) dataset^22^ generated using 10X Visium technology, involving 4,174 spots and 15,427 genes. We performed unsupervised clustering and cell type annotation on this dataset, incorporating information from the Wu et al. study^22^ and the hematoxylin and eosin-stained (H&E) tissue image (Fig.**3a**). This tissue contains five main cell types: tumor cells, smooth muscle cells, normal epithelium, lamina propria, and fibroblast, with the tumor cells located in two distinct regions (Fig.**3a**). HEARTSVG identified 8,020 SVGs, and SpatialDE, SPARK, SPARK-X and scGCO identified 11,190, 12,198, and 13,946, 1,244 SVGs, respectively, at an adjusted p-value cutoff of 0.05. scGCO missed many SVGs with clear spatial expression patterns comparing with other methods (Fig.**S10**). For instance, RPS20, RPS29, ARPC3, and GAS5 exhibited clear and similar spatial expression patterns. scGCO only identified RPS20, HEARTSVG and other three methods successfully identified all four genes. The top 10 genes ranked by HEARTSVG and SPARK-X exhibited more pronounced spatial expression patterns compared to SpatialDE SPARK, and scGCO (Fig.**S11**). SpatialDE’s top 10 selected SVGs displayed minimal spatial patterns, while SPARK and scGCO outperformed SpatialDE to some extent. Notably, SPARK-X demonstrated a preference for selecting SVGs with large stripe patterns, aligning with previous simulation findings. Furthermore, we constructed four artificial genesets as benchmarks to assess the accuracy and robustness of all methods (Fig.**3b**).

Firstly, we selected the top 500 overlaps between HEARTSVG and SPARK-X to form a set of true SVGs (Set 1), while randomly rearranging the expression of these genes resulted in a non-SVGs set (Set 4). Additionally, we introduced noise into both of these sets, generating a set of true SVGs with noise (Set 2) and a set of non-SVGs with noise (Set 3, For more detailed information on the four genesets in **Supplementary 1**). Among these four genesets, HEARTSVG exhibited the highest recalls (average recall = 0.990), as shown in Fig.**3b**. The other four methods exhibited low recalls on the genesets with noise. The SVGs identified by HEARTSVG exhibited significant biological relevance, as confirmed by pathway enrichment analyses conducted for each method. The enrichment analysis results (Fig.**3c**) showed that HEARTSVG displayed smaller p-values and larger gene intersection sizes compared to the other four methods across 19 tumor-related KEGG pathways, including Cancer: Overview and Signal transduction. HEARTSVG demonstrated the highest AUCs (AUC=0.839, 0.718) using single-cell level common gene modules linked with tumor microenvironments^25,26^ and consensus molecular markers of colorectal cancer subtypes^27,28^ as reference standards for true SVGs, HEARTSVG demonstrates the highest AUCs (AUC=0.839, 0.718), underscoring its biological interpretability.

For the identified SVGs, we utilized the auto-clustering module to predict six primary spatial domains and performed enrichment analyses of the SVGs in each spatial domain (Fig. **3e-g****, S12**). Some spatial domains were correlated with specific cell types, consistent with the unsupervised spatial clustering results. The SVGs in spatial domain 4 expressed highly in the muscle cell region and identified many Gene Ontology (GO) terms and KEGG pathways associated with smooth muscle cells (Fig.3**e-g**). The representative genes of spatial domain 4, DES^29,30^, MYL9^31^, and ACTB^32,33^, were essential for the functions of smooth muscle cells. The SVGs of spatial domains 1, 2, 3, and 5 showed high-expression patterns in the tumor cell regions. However, we also identified some spatial domains beyond explained cell types. The SVGs of spatial domains 1 and 2 showed high expression in the left and right tumor cell regions, respectively (Fig.**3e**). The spatial domain 1 was enriched in immune-associated Go terms and KEGG pathway (Fig.**3g**, **S13**). Several representative SVGs in spatial domain 1, such as IGKC, IGHG4, and CD28, which are associated with immune infiltration ^34–36^ (Fig.**3f**). The spatial domain 2 were enriched in the Go terms and KEGG pathways of cell differentiation^37–39^ (Fig.**3f**, **S14**), and included cell markers, such as EPCAM, KRT8, and CLDN3, which are connected with epithelial carcinogenesis, epithelial-mesenchymal transition (EMT) or cancer enhancement. The spatial domain 3 corresponded to the location of tumor cells. Some SVGs of the spatial domain 3, such as B2M^40–43^ and FTL^42–44^ are important encoding antigen genes in many cancers, as well as FTH1^45–47^ and FTL^44,48,49^, which are closely related to iron metabolism in cancer cells. The functional differences between the left and right tumor cells could explain why the spatial expression pattern in the right tumor cell region has clearer boundaries than the left tumor cell region.

**Fig.3.**
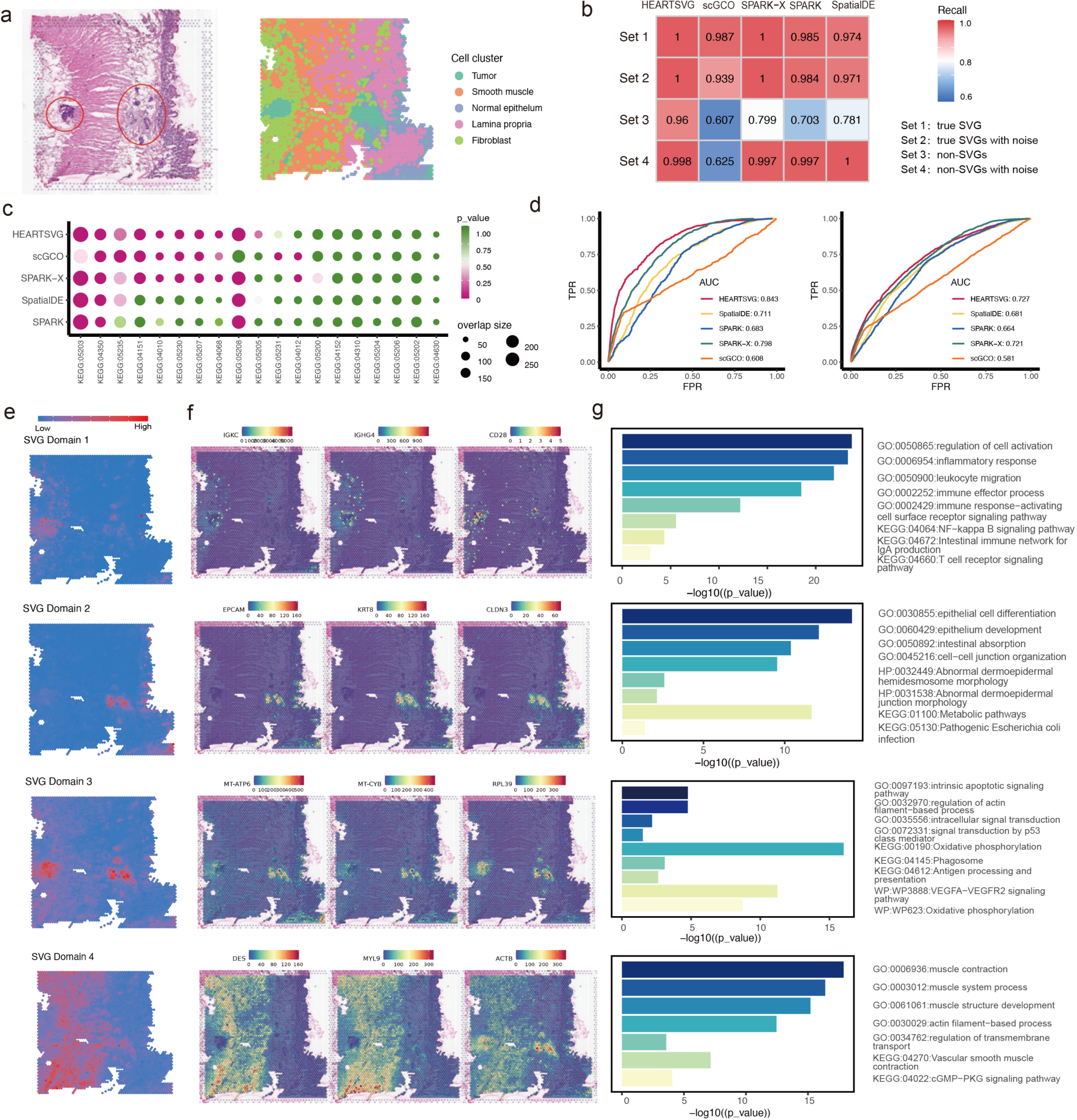
HEARTSVG identifies tumor related SVGs and predicts spatial functional domains with distinct biological functions in the 10X Visium colorectal cancer data. **a,** Original hematoxylin and eosin stained (H&E) tissue image (left) and results of unsupervised spatial clustering (right). The red-circled areas in the HE image represent the tumor regions. **b,** The heatmap depicts the comparison of recall values among four genesets. Set 1 represents the true SVGs set, derived by selecting the top 500 overlaps between HEARTSVG and SPARK-X. Set 4 corresponds to the non-SVGs set, obtained by randomly rearranging the gene expressions within Set 1. Set 2 and Set 3 are generated by introducing noise to the true SVGs set and non-SVGs set, respectively. (More details of four genesets in **Supplementary 1**). **c,** The bubble plot illustrates the results of KEGG pathway enrichment analysis for 19 tumor-related pathways (x-axis) across different methods. Each bubble represents a pathway, and its size corresponds to the overlap gene size of the pathway and SVGs detected by each method. The x-axis and y-axis of the plot represent different methods and the significance (-log10(p-value)). **d,** The ROC curves illustrate the TPR and FPR of four different methods when using common gene modules of tumor microenvironments (left) and consensus molecular markers of colorectal cancer subtypes (right) and as gold standards for true SVGs. AUC is the area under the ROC curve. **d, e,** HEARTSVG predicts four spatial domains based on SVGs and graphed the average expression of SVGs in each spatial domain. Three tumor-related spatial domains (spatial domains 1, 2, and 3) exhibit different spatial average expression patterns. **f,** Representative SVGs correspond to the four predicted spatial domains in Fig. 3e**, 3g,** Enrichment analysis corresponds to the four predicted spatial domains in Fig.**3e**. The length of bars represents the enrichment using-log10(p-value).

We applied HEARTSVG on two other colorectal cancer ST datasets and corresponding liver metastasis ST datasets from the same cohort. HEARTSVG had higher AUC (average AUC=0.788) and recalls (average recalls =0.985) than other methods (Fig. **S12**). In the six colorectal cancer and liver metastasis spatial transcriptomic (ST) datasets, we detected higher expressions of numerous ‘MT-’ genes in the tumor cells compared to the non-tumor region within the colorectal tumor samples. However, this phenomenon was not observed in the liver metastasis samples (Fig.**S19**). We supposed that tumor cells at the primary site of colorectal cancer have higher oxidative phosphorylation (OXPHOS) activity than metastatic liver cancer sites, in line with recent studies showing OXPHOS upregulates in colorectal cancer^50–53^. Overall, HEARTSVG successfully detected SVGs with visually distinct patterns. The auto-clustering module effectively predicted spatial functional domain based on the distinguished SVG patterns positioned in and beyond the cell types.

#### HEARSVG detects SVGs explained by cell types in the Slide-seqV2 cerebellum data

Slide-seqV2^3^ is a spatial transcriptomics technology that achieves transcriptome-wide measurements at near-cellular resolution. We applied HEARTSVG to mouse cerebellum data generated by Slide-seqV2, consisting of 20,141 genes measured on 11,626 spots. The cerebellum plays a crucial role in sensorimotor control^54–56^ and consists of the cortex, white matter, and cerebellar nuclei^57^. The cerebellar cortex comprises three cortical layers^55^ from the outside to the inside: the molecular layer (ML), the Purkinje layer (PCL), and the granular layer (GL). Purkinje cells are a unique kind of neuron in the cerebellar cortex and constitute a slight, convoluted monolayer.

HEARTSVG, SpatialDE, SPARK, SPARK-X and scGCO detected 710, 1,086, 421, 586, and 68 SVGs, respectively. We supported the validity of SVGs detected by HEARTSVG in two pieces of evidence. First, HEARTSVG identified marker genes of specified cell types with spatially restricted expression patterns (Fig.**4a-d**). For example, Mbp (adjusted p=0) in oligodendrocytes, Car8 (adjusted p=1.21 e-10) in Purkinje cells, and Clbn1(adjusted p=0) in granule cells. Notably, HEARTSVG detected the marker genes of Purkinje cells (Fig.4**a-c****, S20**, Tab.**S3**), Car8 (adjusted p=0), Pcp2 (adjusted p=0) and Pcp4 (adjusted p=1.21 e-10), whereas SPARK-failed to identify them. scGCO failed to detect several SVGs with distinct spatial expression patterns, including Calm1, Calm2, Itm2b, among others, which were successfully identified by the other four methods (Fig. **S20**, **S21**, Tab.**S3**). Second, we performed tissue specificity enrichment analysis for the SVGs identified by each method. HEARTSVG, SpatialDE, SPARK, SPARK-X, and scGCO enriched 40, 51, 98, 50 and 26 tissue specificity pathways (Fig.**4d,** Tab.**S2**). The enriched tissue-specific pathways identified by HEARTSVG and scGCO were all related to the brain, with the high percentage (87.5%, 35 cerebellar pathways and 92.31%, 24 cerebellar pathways) of enriched tissue-specific pathways in the cerebellum, and the remaining pathways associated with the cerebral cortex and hippocampus. Although SPARK identified the highest number of pathways (98 pathways), over 40% of these pathways were unrelated to the brain, including 37 skin-specific pathways (37.76%) and three rectum-specific pathways (3.06%). SpatialDE and SPARK-X also identified enriched pathways not associated with the brain, with SpatialDE identifying one rectum pathway (1.96%) and SPARK-X identifying three rectum pathways (6%) and three skin pathways (6%). The heatmap (Fig.**4e**) of SVGs detected by HEARTSVG corresponding to the molecular, Purkinje, and granule layers of the cerebellum confirmed the biological interpretability of the SVGs detected by the HEARTSVG. These findings demonstrated that HEARTSVG is a reliable method for detecting SVGs exhibiting arbitrary spatial patterns in structurally complex tissues, such as the brain.

**Fig.4.**
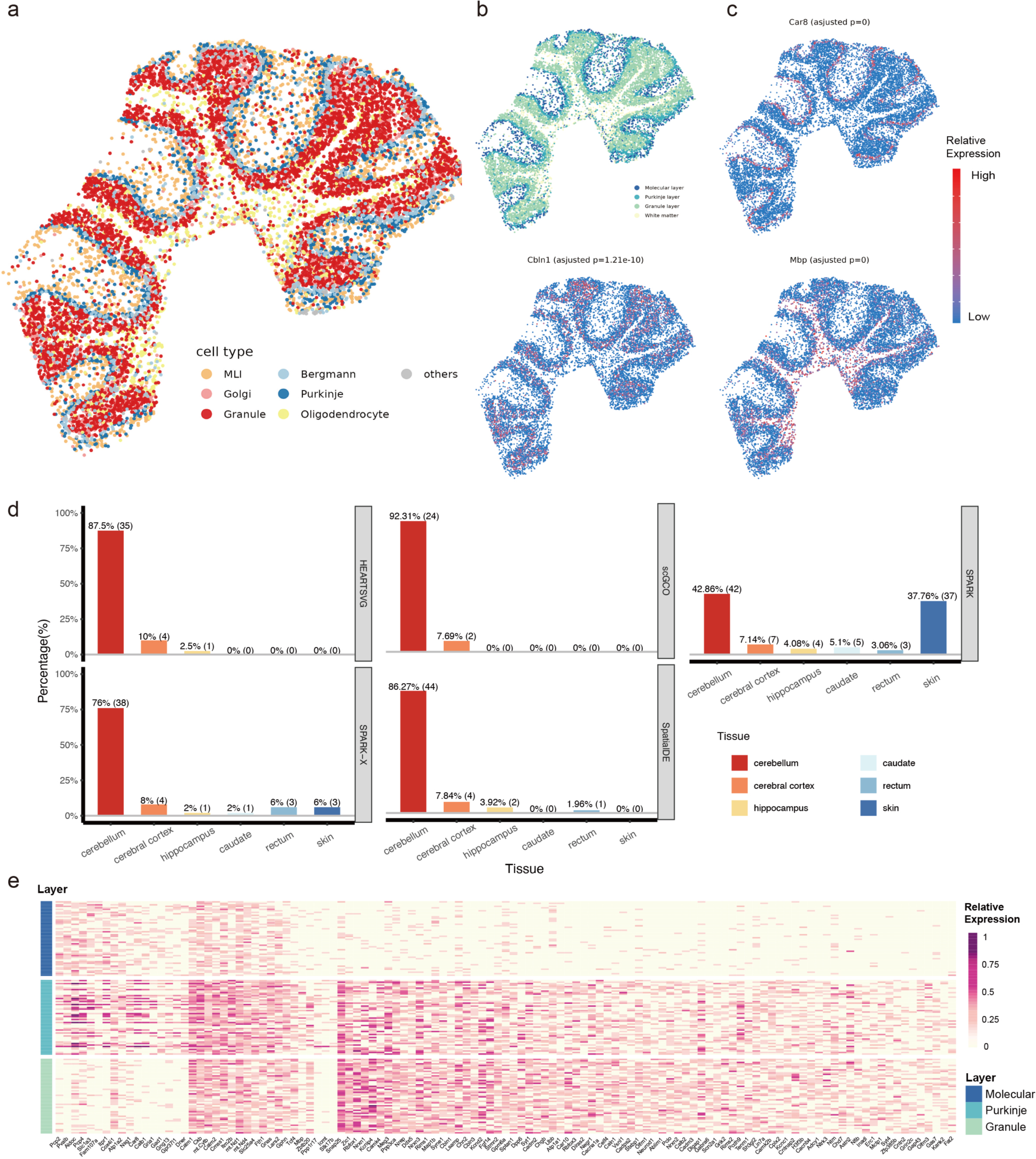
HEARTSVG detects biologically meaningful SVGs in the Slide-seqV2 cerebellum data. **a,** Visualization of unsupervised spatial clustering results. **b,** Visualization of cerebellum layer annotations. **c,** Visualizations of spatial expressions and adjusted p-values of representative SVGs of specified cell types detected by HEARTSVG. Car8 (adjusted p=1.21 e-10) in Purkinje cells, Clbn1(adjusted p=0) in granule cells and Mbp (adjusted p=0) in oligodendrocytes. **d,** The tissue-specific enrichment analysis results for each method, where the x-axis represents different tissues, and the y-axis represents the percentage of tissue-specific pathways. Each panel corresponds to each method. **e,** Heatmap of SVGs expressions in Molecular layer, Purkinje layer, and Granule layer.

#### HEARTSVG identifies marker genes with spatial patterns in the MERFISH preoptic hypothalamus data

We analyzed two datasets of mouse preoptic hypothalamus generated by multiplexed error-robust fluorescence in situ hybridization^58^ (MERFISH). MERFISH enabled spatially resolved RNA analysis of individual cells with high accuracy and high detection efficiency^5^. The data generated through MERFISH were moderately sparse, with more than 40% of the genes detected in more than half of the cells. The first dataset involved 6,112 cells and 155 genes and consisted of eight cell types (Fig.**5a**). The second dataset consisted of 10 cell types (Fig.**5b**) involving 5,665 cells and 161 features (156 genes and five blank controls. HEARTSVG, SpatialDE, SPARK, SPARK-X and scGCO identified 133, 154,149, 141 and 65 genes in the first MERFISH dataset and 128, 161,145, 132 and 46 genes in the second MERFISH dataset. The results of all methods were highly consistent (Fig. **5c**, **S22**). However, SpatialDE misclassified five blank controls as SVGs with top gene ranks. HEARTSVG, SPARK, and SPARK-X reported one blank control as false positive with low ranks and no false positives reported by scGCO. However, scGCO missed some SVGs with clear spatial expression patterns, such as Mbp in the MERFISH data 2 (Fig. **5d**), Nnat in in these two MERFISH data (Fig. **S22**).

**Fig.5.**
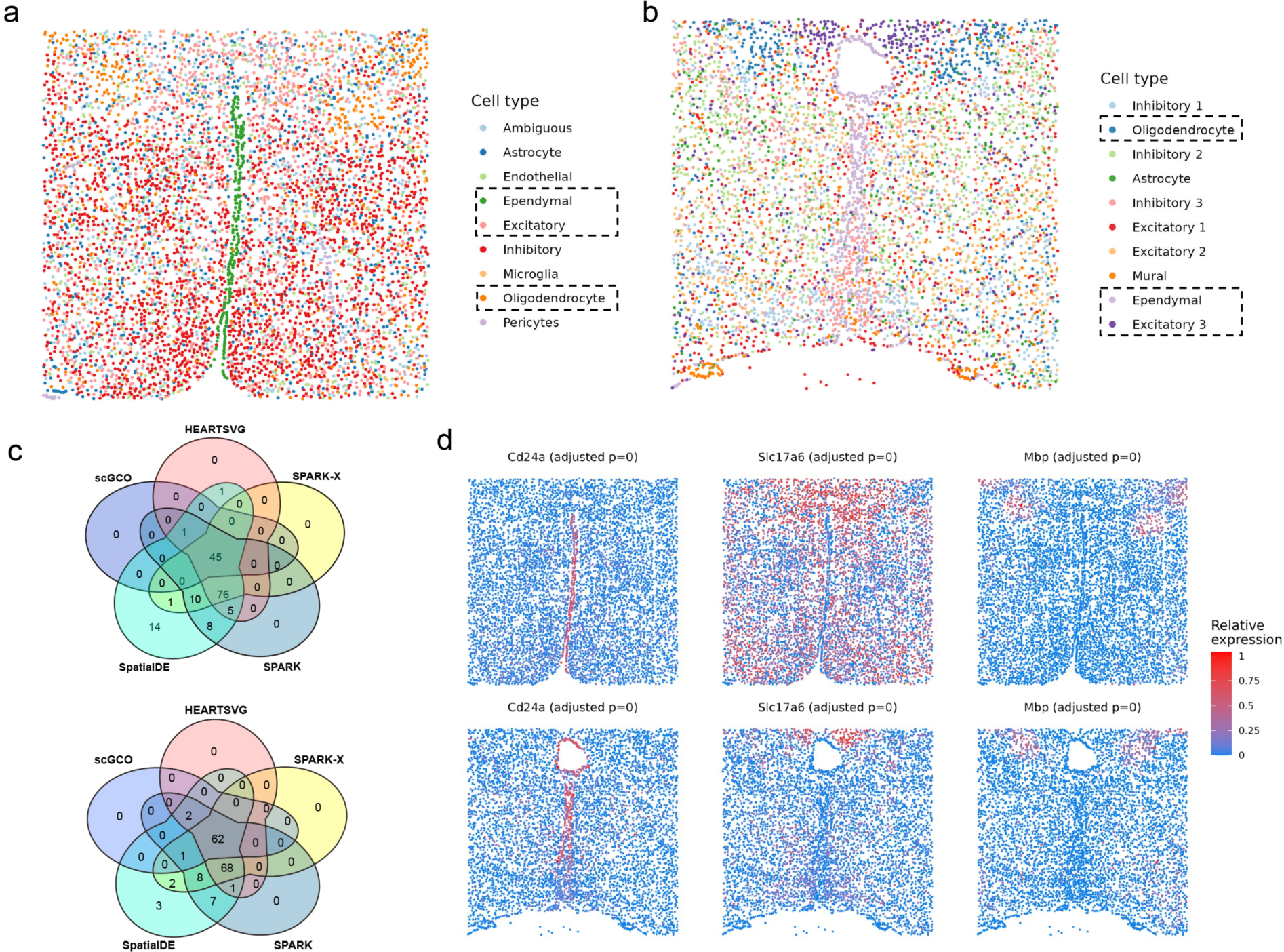
HEARTSVG identifies cell type-specific marker genes with distinct spatial patterns across different MERFISH datasets. **a,** Visualization of known cell annotations of the MERFISH data 1 (6,112 cells with 155 genes). **b,** Visualization of unsupervised spatial clustering results of the MERFISH data 2 (5,665 cells with 156 genes and 5 blank controls). **c,** Venn diagrams of SVGs identified by HEARTSVG, SpatialDE, SPARK, SPARK-X, and scGCO. **d,** Visualizations of representative marker genes for ependymal, excitatory, and oligodendrocyte cell types, along with corresponding adjusted p-values obtained by HEARTSVG in two MERFISH datasets. The panels above of Fig.5**c** and Fig.5**d** correspond to the MERFISH data 1. The panels below correspond to the MERssFISH data 2.

In both datasets, HEARTSVG efficiently identified SVGs associated with cell types spatially located in specific regions (Fig.**5c****, S20**). For example, HEARTSVG detected Cd24a (adjusted p=0), Mlc1 (adjusted p=0), and Nnat (adjusted p=0) as significantly associated SVGs in ependymal^59,60^, Slc17a6 (adjusted p=0) ^61–63^, Cbln2 (adjusted p=0), Necab1 (adjusted p=0), and Ntng1 (adjusted p=0) in excitatory neurons^64,65^. The oligodendrocyte (OD)^59,60,66^ markers, including Mbp (adjusted p=0), Ermn (adjusted p=0), Ndrg1, and Sgk1 (adjusted p=0), were also accurately identified. We utilized the auto-clustering module to obtain multiple spatial domains.

The resulting spatial domains consistently matched their corresponding cell types (Fig. **S22**). For example, in the first data, we predicted two spatial domains corresponding to Oligodendrocyte and Excitatory 3 neuron, respectively. Overall, the auto-clustering module highlighted the usefulness of the software HEARTSVG.

#### HEARTSVG has general applicability across a variety of datasets

To evaluate the generality of HEARTSVG, we applied it to a more comprehensive range of datasets, including mouse olfactory bulb data generated by high-definition spatial transcriptomics (HDST) and ST datasets of two different cancers using 10X Visium. The HDST dataset^4^ was huge and sparse, consisting of 181,367 spots and 19,950 genes, with more than 98% of spots detecting less than 50 genes. Only HEARTSVG, SPARK-X, and scGCO could flawlessly be operated on the HDST data and detected 447, 89, and 0 SVGs, respectively. scGCO failed to identify any SVGs in this sparse HDST dataset. HEARTSVG identified top-ranked SVGs (Gm42418, mt-Rnr1, mt-Rnr2, Cmss1, Gphn) that showed pronounced spatial expression patterns (Fig. **S23**), although visual-spatial expression patterns of genes were challenging to observe in such sparse data.

10X Visium is the most popular ST technology in cancer research. Therefore, we ran our method on more ST data generated by 10X Visium, a primary liver cancer (PLC) ST dataset^23^, and a renal clear cell carcinoma with brain metastasis (RCC-BM) ST dataset^24^ to demonstrate the superior performance of HEARTSVG. As in previous applications, tumor cells were complex and highly heterogeneous, with multiple tumor cell types containing various functions in the same tissue. HEARTSVG effectively identified tumor-related SVGs in cancer ST data and predicted several spatial domains with different functionalities. For example, the PLC ST data contained three distinct tumor cell types. The identification of tumor-associated SVGs by HEARTSVG, along with the prediction of their corresponding spatial domains, revealed a potential synergistic function among these cell types (Fig. **S24**). In the RCC-BM ST data, we found two spatial domains showing different high SVG expressions, corresponding to tumor small nests and tumor medium/big nests^24^ (Fig. **S25**), respectively. The regions of tumor small nests and tumor medium/big nests in this sample were adjacent. Some immune-related genes, such as CD44^67,68^ and CD14^67,69,70^ are highly expressed in the tumor small nest region. Moreover, we found that many tumor-related genes showed higher expression in the “small nests” of tumors than in the “large nests (Fig. **S25**),” which is consistent with the study of Sudmeier et al^24^. SVG detection contributed to providing further insights into intertumoral and intratumoral genetic heterogeneity and complex tumor microenvironments (TME) and cancer mechanisms, which is critical to understanding tumor progression and response to therapy.

## 3. Discussion

We proposed HEARTSVG, a distribution-free, test-based method, for rapid and precise detection of SVGs in large-scale spatial transcriptomic data. Different from existing SVG detection methods ^11,12,15–17^, HEARTSVG adopts an alternative strategy that employs the exclusion of non-SVG genes to infer the presence of SVGs, enabling it to identify SVGs of arbitrary spatial expression patterns with high accuracy, robustness, and generalizability across various types of datasets from different spatial technologies. Benefiting from the test framework and absence of underlying data-generative models, HEARTSVG has superior computational efficiency and scalability, highly suitable for large-scale spatial transcriptomics data. Moreover, the HEARTSVG software offers various functionalities for advanced analysis of SVGs, including auto-clustering, enrichment analysis, and visualization tools.

Our study evaluated the performance of HEARTSVG on both simulated and real ST data, demonstrating its accuracy, robustness, and generality across diverse scenarios, including varying sample sizes, SVG ratios, spatial patterns, and spatial transcriptomic sequencing technologies. HEATSVG had the highest *F*_1_ scores in most simulation scenarios and had good scalability and computational efficiency. Only HEARTSVG, SPARK-X, and scGCO were able to successfully run on simulated datasets of 50,000 cells or greater sample sizes. HEARTSVG and SPARK-X exhibited lower time consumption and higher *F*_1_ scores than scGCO. scGCO achieves excellent FPR control, but its performance is hampered by overlooking a substantial number of SVGs in sparse simulated datasets, due to inaccuracies in candidate region identification. In the simulation with varying SVG proportions, we found that as the proportion of SVGs increased, SPARK-X had increasing FPs while HEARTSVG maintained low FPs. Besides, HEARTSVG can detect SVGs with diverse spatial patterns, while SPARK-X has pattern preferences in recognizing SVGs and has difficulty detecting some non-striped patterns and small area patterns.

We implement HEARTSVG on twelve datasets from four different spatial transcriptome sequencing technologies (10X Visium, Slide-seqV2, HDST, and MERFISH) across three different tissues (colorectal, liver, and brain). HEARTSVG exhibited the highest recalls (average recall=0.985) and AUC (average AUC=0.788), demonstrating its accuracy and robustness across datasets with diverse data characteristics. The brain is a complex organ with intricate structures and a wide variety of cell types in constrained regions^56,71–73^. HEARTSVG can sensitively identify cell-type markers that are restricted to specific brain regions. For example, HEARTSVG identified the markers, Car8, Pcp2, and Pcp4 of the thin and curly Purkinje cell layer, which SPARK-X failed to identify. Despite favorable FPRs, scGCO’s inaccurate identification of candidate regions limits its capacity to fully recognize SVGs with similar spatial expression patterns. SpatialDE misidentified five blank control genes as SVGs with small adjusted p-values in the MERFISH preoptic hypothalamus data. We performed tissue-specific enrichment analysis of the SVGs identified by each method and illustrated the biological benefits of HEARTSVG. The enriched tissue-specific pathways identified by HEARTSVG and scGCO were all related to the brain. In contrast, SPARK identified more than 40% of enriched tissue-specific pathways that were unrelated to the brain in the Slide-seqV2 cerebellum data. This indicates that the reliability of the SVGs identified by SPARK was limited. SpatialDE (1.96%) and SPARK-X (14%) also had pathways unrelated to the brain. Only HEARTSVG, SPARK-X and scGCO can identify SVGs on the huge and sparse HDST dataset (180K∼ spots), demonstrating HEARTSVG’s excellent computing efficiency and scalability.

In this study, we conducted analyses of ST datasets for three different types of cancer (colorectal cancer, primary liver cancer, and renal cell carcinoma brain metastasis), which were generated using 10X Visium - a widely used commercial ST technology in cancer research. The ST data of tumors contained few cell types and their SVGs are primarily associated with tumor cells. HEARTSVG performed well on different cancer ST datasets, and pathway analysis results demonstrated its ability to identify many tumor-related SVGs. HEARTSVG (8 significant pathways), SPARK-X (8 significant pathways) and scGCO (7 significant pathways) identified more cancer-related KEGG pathways than SpatialDE (2 significant pathways) and SPARK (2 significant pathways) in the 10X Visium colorectal cancer data. Furthermore, the SVG auto-clustering module of the software HEARTSVG facilitated the prediction of different tumor-associated spatial domains with distinct spatial expression patterns. In the colorectal cancer ST data, tumor cells were located in two non-adjacent regions of the sample. We discovered that two tumor-associated spatial domains had high expression patterns in only one tumor cell region instead of both, as shown in Fig.**3**. Enrichment analysis revealed distinct biological processes and functions associated with the two spatial domains. We observed similar phenomena in the ST datasets of primary liver cancer and renal clear cell carcinoma with brain metastasis. In the PLC ST data, many SVGs were highly expressed in both tumor cell subtypes 1 and 3, constituting a common spatial functional domain. In the RCC-BM ST dataset, we identified two adjacent spatial domains based on different SVG clusters, corresponding to tumor small nests and tumor medium/big nests, respectively. Spatial domain prediction based on SVGs has revealed tumors’ intricate functional diversity and synergistic interactions beyond cellular classifications, shedding new light on the biological complexity of tumor tissues.

Overall, HEARTSVG is a powerful method for detecting spatially variable genes with the ability to identify spatial expression patterns of arbitrary shapes. Moreover, the inclusion of an auto-clustering module in the HEARTSVG software enhances the understanding of the biological process, demonstrating the versatility and potential of HEARTSVG in spatial transcriptomics data analysis. However, HEARTSVG has such limitations as relying solely on spatial coordinates. In future studies combing gene expression with corresponding H&E tissue images, incorporating information from H&E tissue images will provide a more comprehensive understanding of the cellular mechanism in disease progression.

## 4. Methods

### Identification of spatially variable genes

In spatial transcriptomics (ST) data, each gene can be represented by a vector containing three elements: the gene *g* = (*x*, *y*, *e*)*^T^*, where *x*, *y*, and *e* correspond to the row coordinates, column coordinates, and the expression counts of the gene on the spot at the (*x*, *y*) (Fig.**S7**). To simplify notation, we assume in the following proof that there is only one gene *g*. HEARTSVG tested for each gene, so the “*only one gene*” assumption does not affect the derivation and conclusion. We determine whether g is an SVG by testing whether the expression of gene *g* is randomly distributed in the ST data. In practice, we assume that the expression counts of the non-SVG gene at a given location (*x_i_*, *y_j_*) are independent of expressions at nearby locations. Therefore, we applied the Portmanteau test to test several autocorrelations of *r_t_* that are simultaneously at zero to determine whether the gene is an SVG. *r_t_* is the gene marginal expression series after the semi-pooling step. The null and alternative hypotheses are:

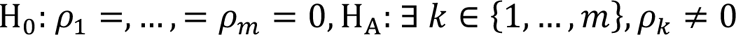

To simplify the symbolic representation, we rewrite the subscript of the marginal expression series as **r** = (*r*_1_, . ., *r_t_*,…, *r_T_*)*^T^*, define the autocovariance of order *k* as: *γ_k_* = *Cov*(*r_t_*, *r_t_*_-*k*_) = *Cov*(*r_t_*, *r_t_*_+*k*_), for all *k* ≥ 0, and the *k* − th order autocorrelation (ACF) as 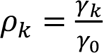.

If gene *g* is non-SVG without a spatial pattern in ST data, our purpose is to test the null hypothesis: H_0_: *ρ*_1_= ⋯ = *ρ_m_* = 0. The test statistic is defined as 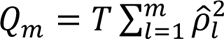 followed by chi-distribution with *m* degree of freedom, where 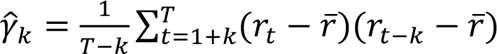, *k* = 0,…, *T* − 1, *r̅* is the mean of **r**, and introduce 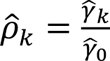. The *P* value for testing the null hypothesis can be calculated by *p* = P(χ^2^(*df* = *m*) > *Q_m_* | H_0_ is true). We combined all individual p-values into a single p-value by Stouffer’s method. The Stouffer’s statistic is defined ^as^ 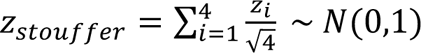, where *z_i_* = Φ^-1^(1 – *p_i_*), Φ^-1^(·) is the inverse of the cumulative distribution function of a standard normal distribution. Hence, the combined p-value of four p-values is calculated by ***p_c_* = 1 − Φ(*z*)**. We use the continuously adjusted combined p-value to determine whether a gene is an SVG. If the adjusted p-value of a gene is less than 0.05, it is recognized as an SVG.

### Auto-clustering module

The auto-clustering module utilizes the hierarchical clustering algorithm and includes the following steps.

Step 1: Calculate the similarity between each pair of genes based on spatial expression and generation of the distance matrix.

Step 2: Construct a clustering tree based on the distance matrix using the complete linkage criterion. The resulting hierarchy of clusters can be visualized as a dendrogram.

Step 3: Determination of the final clustering results by cutting the dendrogram at a certain height or distance threshold. The cutting height is chosen using the maximum breakpoint of all breakpoints selected by the Yamamoto test.

We predicted spatial domains based on each SVG cluster’s regions and expression levels.

### Simulation Design

We generated extensive simulation scenarios to evaluate the performances of HEARTSVG and three other existing SVG methods. Each scenario had 50 replications. For spatial expression pattern settings, we set three different spatial expression patterns (hotspot, streak, gradient) and random patterns (Fig.**1b**), following Trendsceek and SPARK-X. We generated the spatial locations of spots by the random-point-pattern Poisson process (intensity parameter lambda=0.5). The expression counts are generated from the zero-inflated negative binomial distribution (ZINB). The ZINB model has two possible data generation processes. Process 1 of ZINB is chosen with probability *φ*, and Process 2 with probability 1 − *φ*. Process 1 generated only zero counts; the Process 2 generated expression counts from the negative binomial model with parameters *NB*(*size*, *mu*). In all simulations, we set 10,000 genes. For Simulation 1, the SVGs ratios of all simulation datasets are 10% (1000 SVGs vs. 9000 non-SVGs). We varied the number of spots from 1500, 3000, 5000, 10,000, 30,000, and 50,000. For both SVGs and non-SVGs, we varied the dispersion parameter of the Process 2 varied from 0.15, 0.5, and 1.5. The SVGs in non-marked areas and non-SVGs had the same ZINB parameters. The probability *φ* is 0.8, and the mean parameters varied from 0.5, 5, and 15. For SVGs in marked areas, the expression counts are generated from the ZINB model with 1/3 *φ* and two times the mean parameter of the non-marked areas. This is set up to ensure that the SVG expression rate and expression values are different in the pattern and non-pattern areas, which is related to the biological significance of the SVGs.

Simulation1 changed the sample size, mu, and size and used the *F*_1_ score to evaluate the identification performance of all methods. The *F*_1_ score is a measurement of accuracy that balances precision and recall. The calculations are as follows. For Simulation 2, we compared the number of false positives and *F*_1_ scores of HEARTSVG and SPARK-X with the variation in the proportion of SVG. We varied the ratios of SVGs from 0%, 5%, 10%, and 50%, and the number of spots from 3000, 5000, and 10,000. Other simulation settings were similar to simulation1. Like other SVG detection methods, we use the continuously adjusted p-value to determine whether a gene is an SVG. If the adjusted p-value of a gene is less than 0.05, it is identified as an SVG. Based on this criterion, we converted continuously adjusted p-values to binary results and calculated performance indices. Performance indices calculations were as follows.

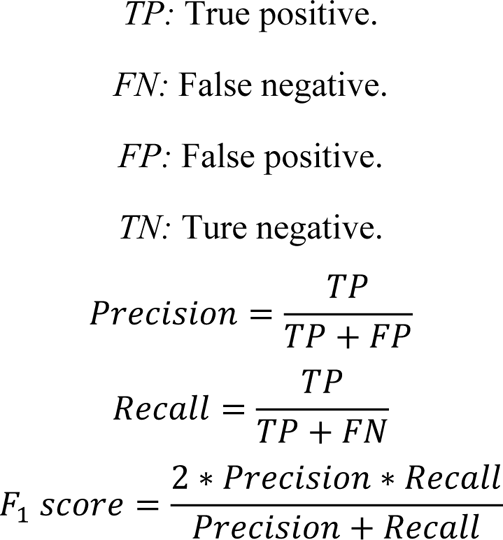

## Acknowledgements

The computations in this paper were run on the Siyuan-1 cluster supported by the Center for High Performance Computing at Shanghai Jiao Tong University. We express our gratitude to Ph.D. students Yudi Chen and Panpan Zhang for their valuable feedback on the manuscript. We sincerely appreciate the helpful advice on codes provided by Ph.D. student Jie Zhou and Mr. Zhaochang Yang. Additionally, we are grateful to Ms. Kaiqi Zhang, Ms. Xiwen Sun, Ms. Congwen Xiao, and Ms. Xiaoya Sun for their helpful discussions.

## Ethical approval

Ethical approval was not required for this study as it did not involve human or animal subjects.

## Data Availability

We used 12 publicly available actual ST datasets in applications. These data can be obtained by the following:

1. Six colorectal cancer and corresponding liver metastasis ST datasets are available from http://www.cancerdiversity.asia/scCRLM/.
2. The Slideseq-V2 data is available from https://singlecell.broadinstitute.org/single_cell/study/SCP815/highly-sensitive-spatial-transcriptomics-at-near-cellular-resolution-with-slide-seqv2#study-summary
3. The MERFISH is available from https://datadryad.org/stash/dataset/doi:10.5061/dryad.8t8s248.
4. The HDST data can be obtained by the GEO database repository under accession code <u>GSE130682</u>.
5. The primary liver cancer data is available from https://ngdc.cncb.ac.cn/gsa-human/browse/HRA000437.
6. The renal clear cell cancer brain metastasis data can be obtained by the GEO database repository under accession code <u>GSE179572.</u>

## Code Availability

HEARTSVG is implemented in R, and the source codes have been deposited at https://github.com/cz0316/HEARTSVG.git.

## Author Contributions

X.Y., Z.Y., and S.M. designed the HEARTSVG algorithm and the simulation framework. X.Y. implemented the HEARTSVG software, performed data analyses, and conducted comparisons. Z.Y. and S.M. secured funding for the study. Y.M., R.G. and T.W. were responsible for dataset preprocessing. Y.W., S.C., and B.F. contributed to figure design and conducted analyses using real data. X.Y. and Z.Y. wrote this paper. All authors have reviewed and approved the final manuscript.

## References

1. Crosetto, N., Bienko, M. & van Oudenaarden, A. Spatially resolved transcriptomics and beyond. Nat. Rev. Genet. 16, 57–66 (2015).

2. Moses, L. & Pachter, L. Museum of spatial transcriptomics. Nat. Methods 19, 534–546 (2022).

3. Stickels, R. R. et al. Highly sensitive spatial transcriptomics at near-cellular resolution with Slide-seqV2. Nat. Biotechnol. 39, 313–319 (2021).

4. Vickovic, S. et al. High-definition spatial transcriptomics for in situ tissue profiling. Nat. Methods 16, 987–990 (2019).

5. Ståhl, P. L. et al. Visualization and analysis of gene expression in tissue sections by spatial transcriptomics. Science 353, 78–82 (2016).

6. Xia, C., Fan, J., Emanuel, G., Hao, J. & Zhuang, X. Spatial transcriptome profiling by MERFISH reveals subcellular RNA compartmentalization and cell cycle-dependent gene expression. Proc. Natl. Acad. Sci. 116, 19490–19499 (2019).

7. Rao, A., Barkley, D., França, G. S. & Yanai, I. Exploring tissue architecture using spatial transcriptomics. Nature 596, 211–220 (2021).

8. Williams, C. G., Lee, H. J., Asatsuma, T., Vento-Tormo, R. & Haque, A. An introduction to spatial transcriptomics for biomedical research. Genome Med. 14, 68 (2022).

9. Larsson, L., Frisén, J. & Lundeberg, J. Spatially resolved transcriptomics adds a new dimension to genomics. Nat. Methods 18, 15–18 (2021).

10. Zeng, Z., Li, Y., Li, Y. & Luo, Y. Statistical and machine learning methods for spatially resolved transcriptomics data analysis. Genome Biol. 23, 83 (2022).

11. Svensson, V., Teichmann, S. A. & Stegle, O. SpatialDE: identification of spatially variable genes. Nat. Methods 15, 343–346 (2018).

12. Zhu, J., Sun, S. & Zhou, X. SPARK-X: non-parametric modeling enables scalable and robust detection of spatial expression patterns for large spatial transcriptomic studies. Genome Biol. 22, 184 (2021).

13. Dries, R. et al. Advances in spatial transcriptomic data analysis. Genome Res. 31, 1706–1718 (2021).

14. Atta, L. & Fan, J. Computational challenges and opportunities in spatially resolved transcriptomic data analysis. Nat. Commun. 12, 5283 (2021).

15. Edsgärd, D., Johnsson, P. & Sandberg, R. Identification of spatial expression trends in single-cell gene expression data. Nat. Methods 15, 339–342 (2018).

16. Sun, S., Zhu, J. & Zhou, X. Statistical analysis of spatial expression patterns for spatially resolved transcriptomic studies. Nat. Methods 17, 193–200 (2020).

17. Zhang, K., Feng, W. & Wang, P. Identification of spatially variable genes with graph cuts. Nat. Commun. 13, 5488 (2022).

18. Escanciano, J. C. & Lobato, I. N. An automatic Portmanteau test for serial correlation. J. Econom. 151, 140–149 (2009).

19. Box, G. E. P. & Pierce, D. A. Distribution of Residual Autocorrelations in Autoregressive-Integrated Moving Average Time Series Models. J. Am. Stat. Assoc. 65, 1509–1526 (1970).

20. Stouffer, S. A., Suchman, E. A., Devinney, L. C., Star, S. A. & Williams Jr., R. M. The American soldier: Adjustment during army life. (Studies in social psychology in World War II), *Vol.* 1. xii, 599 (Princeton Univ. Press, 1949).

21. Lipták, Tamás. On the combination of independent tests. Magy. Tud Akad Mat Kut. Int Kozl 3, 171–197 (1958).

22. Wu, Y. et al. Spatiotemporal Immune Landscape of Colorectal Cancer Liver Metastasis at Single-Cell Level. Cancer Discov. 12, 134–153 (2022).

23. Wu, R. et al. Comprehensive analysis of spatial architecture in primary liver cancer. Sci. Adv. 7, eabg3750 (2021).

24. Sudmeier, L. J. et al. Distinct phenotypic states and spatial distribution of CD8+ T cell clonotypes in human brain metastases. Cell Rep. Med. 3, 100620 (2022).

25. Xue, R. et al. Liver tumour immune microenvironment subtypes and neutrophil heterogeneity. Nature 612, 141–147 (2022).

26. Wu, S. Z. et al. A single-cell and spatially resolved atlas of human breast cancers. Nat. Genet. 53, 1334–1347 (2021).

27. Dienstmann, R. et al. Colorectal Cancer Subtyping Consortium (CRCSC) Identifies Consensus of Molecular Subtypes. Ann. Oncol. 25, ii115 (2014).

28. Guinney, J. et al. The consensus molecular subtypes of colorectal cancer. Nat. Med. 21, 1350–1356 (2015).

29. Takase, S., Leo, M. A., Nouchi, T. & Lieber, C. S. Desmin distinguishes cultured fat-storing cells from myofibroblasts, smooth muscle cells and fibroblasts in the rat. J. Hepatol. 6, 267–276 (1988).

30. Council, L. & Hameed, O. Differential expression of immunohistochemical markers in bladder smooth muscle and myofibroblasts, and the potential utility of desmin, smoothelin, and vimentin in staging of bladder carcinoma. Mod. Pathol. 22, 639–650 (2009).

31. Moreno, C. A. et al. Homozygous deletion in MYL9 expands the molecular basis of megacystis–microcolon–intestinal hypoperistalsis syndrome. Eur. J. Hum. Genet. 26, 669–675 (2018).

32. Lehtonen, H. J. et al. Segregation of a Missense Variant in Enteric Smooth Muscle Actin γ-2 With Autosomal Dominant Familial Visceral Myopathy. Gastroenterology 143, 1482–1491.e3 (2012).

33. Weymouth, N., Shi, Z. & Rockey, D. C. Smooth muscle α actin is specifically required for the maintenance of lactation. Dev. Biol. 363, 1–14 (2012).

34. Berntsson, J., Nodin, B., Eberhard, J., Micke, P. & Jirström, K. Prognostic impact of tumour-infiltrating B cells and plasma cells in colorectal cancer: 2.1.5 Tumor Immunology and Microenvironment. Int. J. Cancer 139, 1129–1139 (2016).

35. Wouters, M. C. A. & Nelson, B. H. Prognostic Significance of Tumor-Infiltrating B Cells and Plasma Cells in Human Cancer. Clin. Cancer Res. 24, 6125–6135 (2018).

36. Berntsson, J. et al. The clinical impact of tumour-infiltrating lymphocytes in colorectal cancer differs by anatomical subsite: A cohort study: The clinical impact of tumour-infiltrating lymphocytes. Int. J. Cancer 141, 1654–1666 (2017).

37. Sfakianos, J. P. et al. Epithelial plasticity can generate multi-lineage phenotypes in human and murine bladder cancers. Nat. Commun. 11, 2540 (2020).

38. Khaliq, A. M. et al. Refining colorectal cancer classification and clinical stratification through a single-cell atlas. Genome Biol. 23, 113 (2022).

39. Hassan, S., Blick, T., Thompson, E. W. & Williams, E. D. Diversity of Epithelial-Mesenchymal Phenotypes in Circulating Tumour Cells from Prostate Cancer Patient-Derived Xenograft Models. Cancers 13, 2750 (2021).

40. Wang, H., Liu, B. & Wei, J. Beta2-microglobulin(B2M) in cancer immunotherapies: Biological function, resistance and remedy. Cancer Lett. 517, 96–104 (2021).

41. Janikovits, J. et al. High numbers of PDCD1 (PD-1)-positive T cells and *B2M* mutations in microsatellite-unstable colorectal cancer. OncoImmunology 7, e1390640 (2018).

42. Sade-Feldman, M. et al. Resistance to checkpoint blockade therapy through inactivation of antigen presentation. Nat. Commun. 8, 1136 (2017).

43. Jhunjhunwala, S., Hammer, C. & Delamarre, L. Antigen presentation in cancer: insights into tumour immunogenicity and immune evasion. Nat. Rev. Cancer 21, 298–312 (2021).

44. Wang, K. et al. Identification of tumor-associated antigens by using SEREX in hepatocellular carcinoma. Cancer Lett. 281, 144–150 (2009).

45. Liu, Z., Arcos, M., Martin, D. R. & Xue, X. Myeloid FTH1 Deficiency Protects Mice From Colitis and Colitis-associated Colorectal Cancer via Reducing DMT1-Imported Iron and STAT3 Activation. Inflamm. Bowel Dis. izad009 (2023) doi:10.1093/ibd/izad009.

46. Chan, J. J. et al. A FTH1 gene:pseudogene:microRNA network regulates tumorigenesis in prostate cancer. Nucleic Acids Res. 46, 1998–2011 (2018).

47. Liu, N. Q. et al. Ferritin Heavy Chain in Triple Negative Breast Cancer: A Favorable Prognostic Marker that Relates to a Cluster of Differentiation 8 Positive (CD8+) Effector T-cell Response. Mol. Cell. Proteomics 13, 1814–1827 (2014).

48. Jézéquel, P. et al. Validation of tumor-associated macrophage ferritin light chain as a prognostic biomarker in node-negative breast cancer tumors: A multicentric 2004 national PHRC study. Int. J. Cancer 131, 426–437 (2012).

49. Tang, X. Tumor-associated macrophages as potential diagnostic and prognostic biomarkers in breast cancer. Cancer Lett. 332, 3–10 (2013).

50. Ashton, T. M., McKenna, W. G., Kunz-Schughart, L. A. & Higgins, G. S. Oxidative Phosphorylation as an Emerging Target in Cancer Therapy. Clin. Cancer Res. 24, 2482–2490 (2018).

51. Elgendy, M. et al. Combination of Hypoglycemia and Metformin Impairs Tumor Metabolic Plasticity and Growth by Modulating the PP2A-GSK3β-MCL-1 Axis. Cancer Cell 35, 798–815.e5 (2019).

52. Birsoy, K. et al. Metabolic determinants of cancer cell sensitivity to glucose limitation and biguanides. Nature 508, 108–112 (2014).

53. Chekulayev, V. et al. Metabolic remodeling in human colorectal cancer and surrounding tissues: alterations in regulation of mitochondrial respiration and metabolic fluxes. Biochem. Biophys. Rep. 4, 111–125 (2015).

54. D’Angelo, E. & De Zeeuw, C. I. Timing and plasticity in the cerebellum: focus on the granular layer. Trends Neurosci. 32, 30–40 (2009).

55. MacKenzie-Graham, A. et al. Purkinje cell loss in experimental autoimmune encephalomyelitis. NeuroImage 48, 637–651 (2009).

56. Zatorre, R. J., Fields, R. D. & Johansen-Berg, H. Plasticity in gray and white: neuroimaging changes in brain structure during learning. Nat. Neurosci. 15, 528–536 (2012).

57. Sillitoe, R. V. & Joyner, A. L. Morphology, Molecular Codes, and Circuitry Produce the Three-Dimensional Complexity of the Cerebellum. Annu. Rev. Cell Dev. Biol. 23, 549–577 (2007).

58. Moffitt, J. R. et al. Molecular, spatial, and functional single-cell profiling of the hypothalamic preoptic region. Science 362, eaau5324 (2018).

59. Sjöstedt, E. et al. An atlas of the protein-coding genes in the human, pig, and mouse brain. Science 367, eaay5947 (2020).

60. Shah, P. T. et al. Single-Cell Transcriptomics and Fate Mapping of Ependymal Cells Reveals an Absence of Neural Stem Cell Function. Cell 173, 1045–1057.e9 (2018).

61. Erwin, S. R. et al. Spatially patterned excitatory neuron subtypes and projections of the claustrum. eLife 10, e68967 (2021).

62. Mickelsen, L. E. et al. Single-cell transcriptomic analysis of the lateral hypothalamic area reveals molecularly distinct populations of inhibitory and excitatory neurons. Nat. Neurosci. 22, 642–656 (2019).

63. Mizuguchi, R. et al. Ascl1 and Gsh1/2 control inhibitory and excitatory cell fate in spinal sensory interneurons. Nat. Neurosci. 9, 770–778 (2006).

64. Seigneur, E. & Südhof, T. C. Cerebellins are differentially expressed in selective subsets of neurons throughout the brain: SEIGNEUR and SÜDHOF. J. Comp. Neurol. 525, 3286–3311 (2017).

65. Xie, Z. et al. Transcriptomic encoding of sensorimotor transformation in the midbrain. eLife 10, e69825 (2021).

66. Marechal, D. et al. N-myc downstream regulated family member 1 (NDRG1) is enriched in myelinating oligodendrocytes and impacts myelin degradation in response to demyelination. Glia 70, 321–336 (2022).

67. Nalio Ramos, R., et al. Tissue-resident FOLR2+ macrophages associate with CD8+ T cell infiltration in human breast cancer. Cell 185, 1189–1207.e25 (2022).

68. Nie, P. et al. A YAP/TAZ-CD54 axis is required for CXCR2 −CD44 − tumor-specific neutrophils to suppress gastric cancer. Protein Cell pwac045 (2022) doi:10.1093/procel/pwac045.

69. Wu, K., Kryczek, I., Chen, L., Zou, W. & Welling, T. H. Kupffer Cell Suppression of CD8+ T Cells in Human Hepatocellular Carcinoma Is Mediated by B7-H1/Programmed Death-1 Interactions. Cancer Res. 69, 8067–8075 (2009).

70. Sautès-Fridman, C. et al. Tumor microenvironment is multifaceted. Cancer Metastasis Rev. 30, 13–25 (2011).

71. Klemm, F. et al. Interrogation of the Microenvironmental Landscape in Brain Tumors Reveals Disease-Specific Alterations of Immune Cells. Cell 181, 1643–1660.e17 (2020).

72. Silbereis, J. C., Pochareddy, S., Zhu, Y., Li, M. & Sestan, N. The Cellular and Molecular Landscapes of the Developing Human Central Nervous System. Neuron 89, 248–268 (2016).

73. Ecker, J. R. et al. The BRAIN Initiative Cell Census Consortium: Lessons Learned toward Generating a Comprehensive Brain Cell Atlas. Neuron 96, 542–557 (2017).

