## Supplementary 1 for "HEARTSVG: a fast and accurate method for spatially variable gene identification in large-scale spatial transcriptomic data"

#### 1. Simulation Results

##### 1.1 Simulation 1 additional figures

We used $F_1$ score, recall, precision, specificity, and false positive (FP) to comprehensively evaluate the performance of HEARTSVG and SpatialDE, SPARK, and SPARK-X. All simulation settings were illustrated in the two sections, Simulation and Methods. Each simulation scenario has 50 replications. Simulation datasets were generated by varying the sample size (from 1500 to 50000), ZINB parameters ($size = 0.15,\ 0.5,\ 1.5, mu = 0.5, 5, 15$), and SVG percentages (hotspot, streak =5%, gradient=15%). The paper presented the simulation scenario with ZINB parameters ($size = 0.5, mu = 0.5$). The complete simulation results were shown in this supplementary.

**All simulation results of the hotspot pattern**

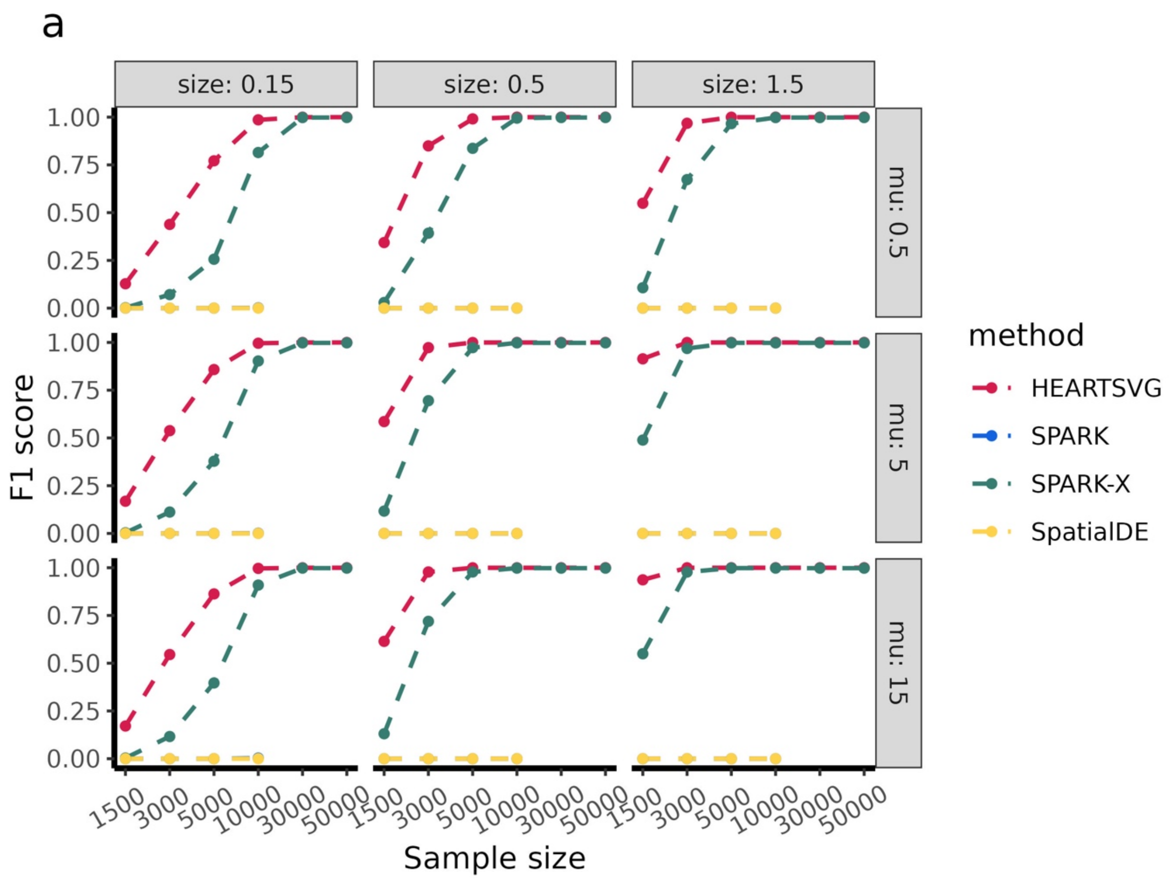

b

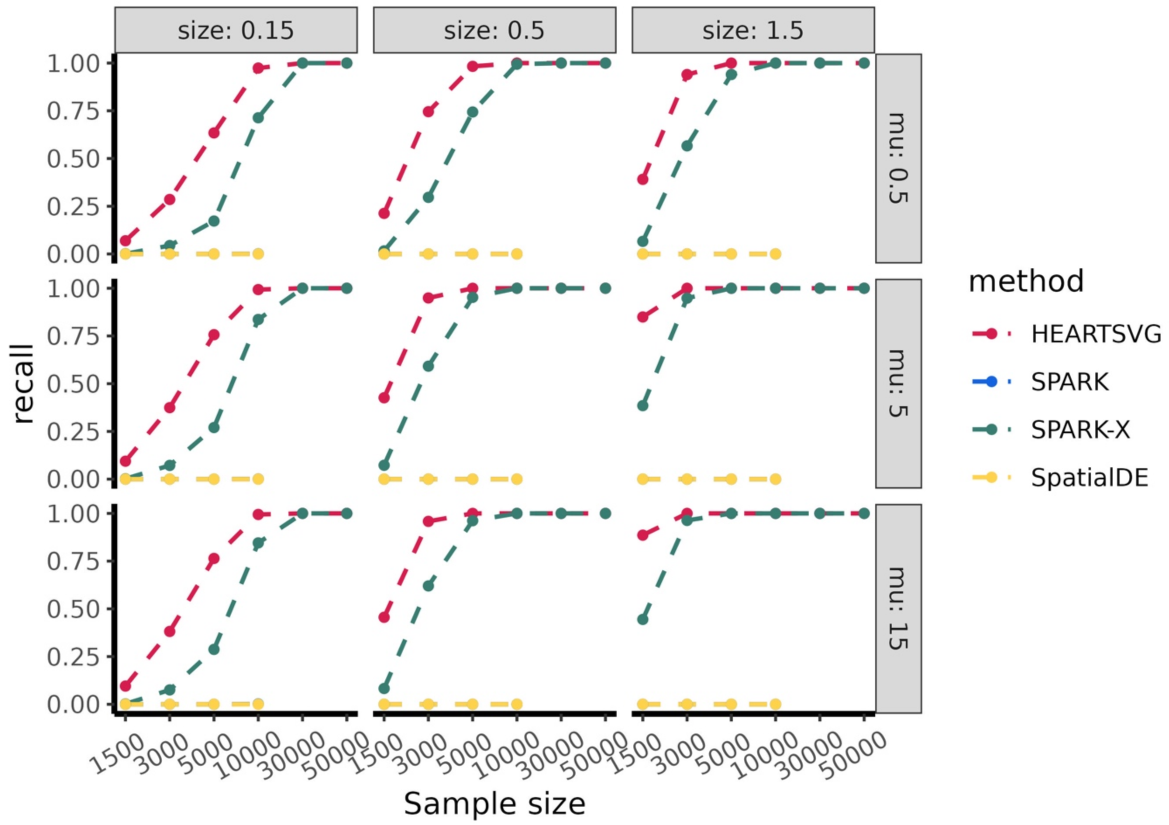
size: 0.15
size: 0.5
size: 1.5
mu: 0.5
mu: 5
mu: 15
recall
1.00
0.75
0.50
0.25
0.00
1500
3000
5000
10000
30000
50000
Sample size
method
HEARTSVG
SPARK
SPARK-X
SpatialDE

c

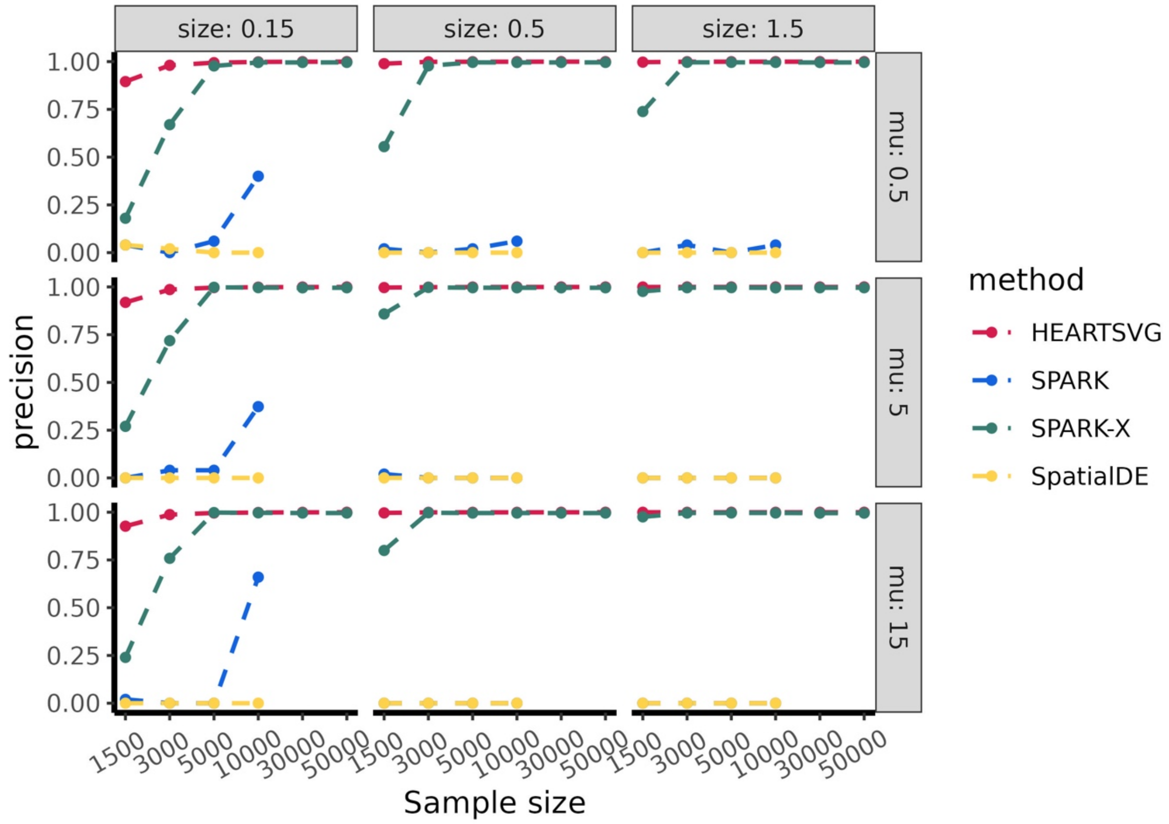
size: 0.15
size: 0.5
size: 1.5
mu: 0.5
mu: 5
mu: 15
precision
1.00
0.75
0.50
0.25
0.00
1500
3000
5000
10000
30000
50000
Sample size
method
HEARTSVG
SPARK
SPARK-X
SpatialDE

d

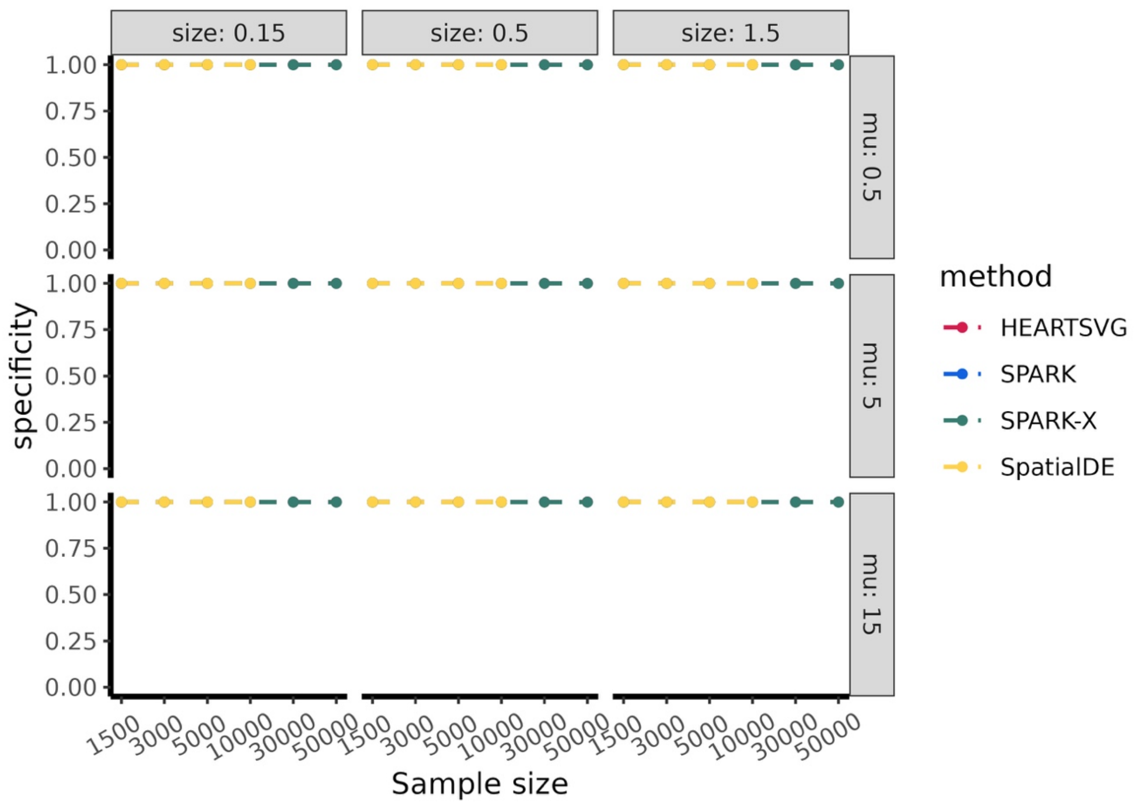


e

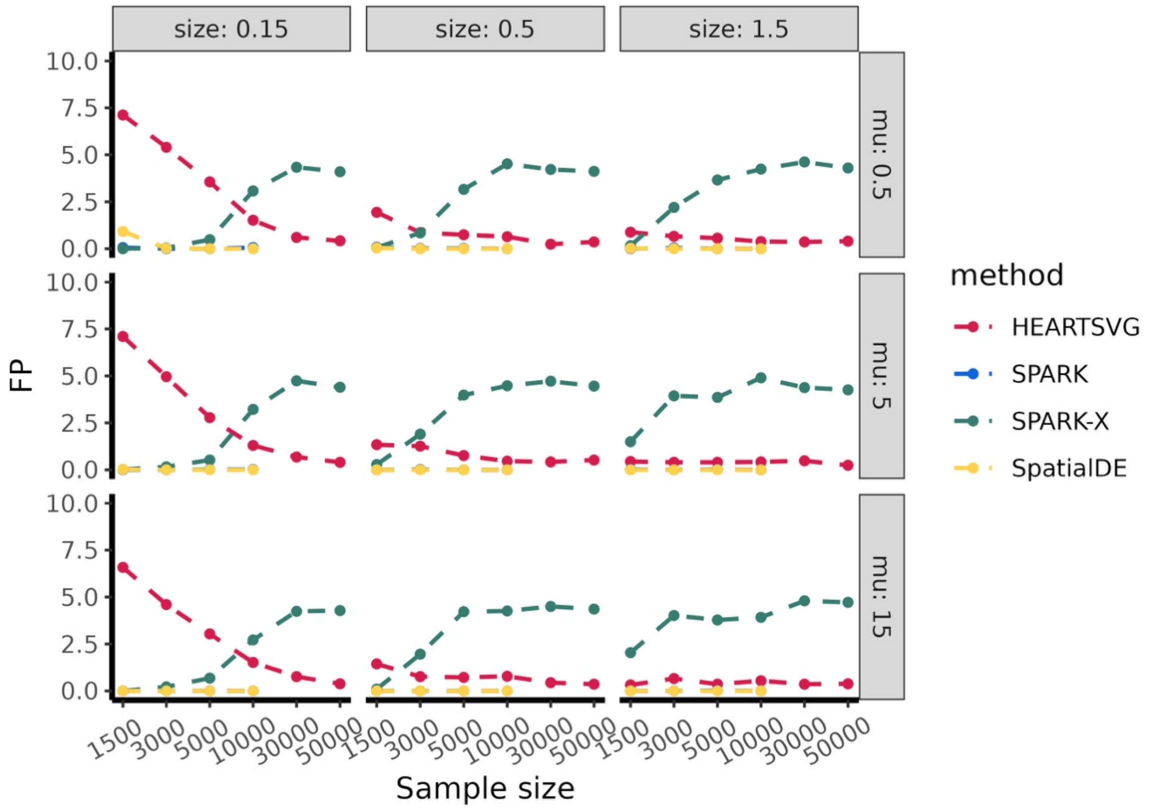


**Figure S1** Simulation results of hotspot pattern. **(a)** F1 score, **(b)** recall, **(c)** precision, **(d)** specificity, **(e)** FP.

**All simulation results of streak pattern**

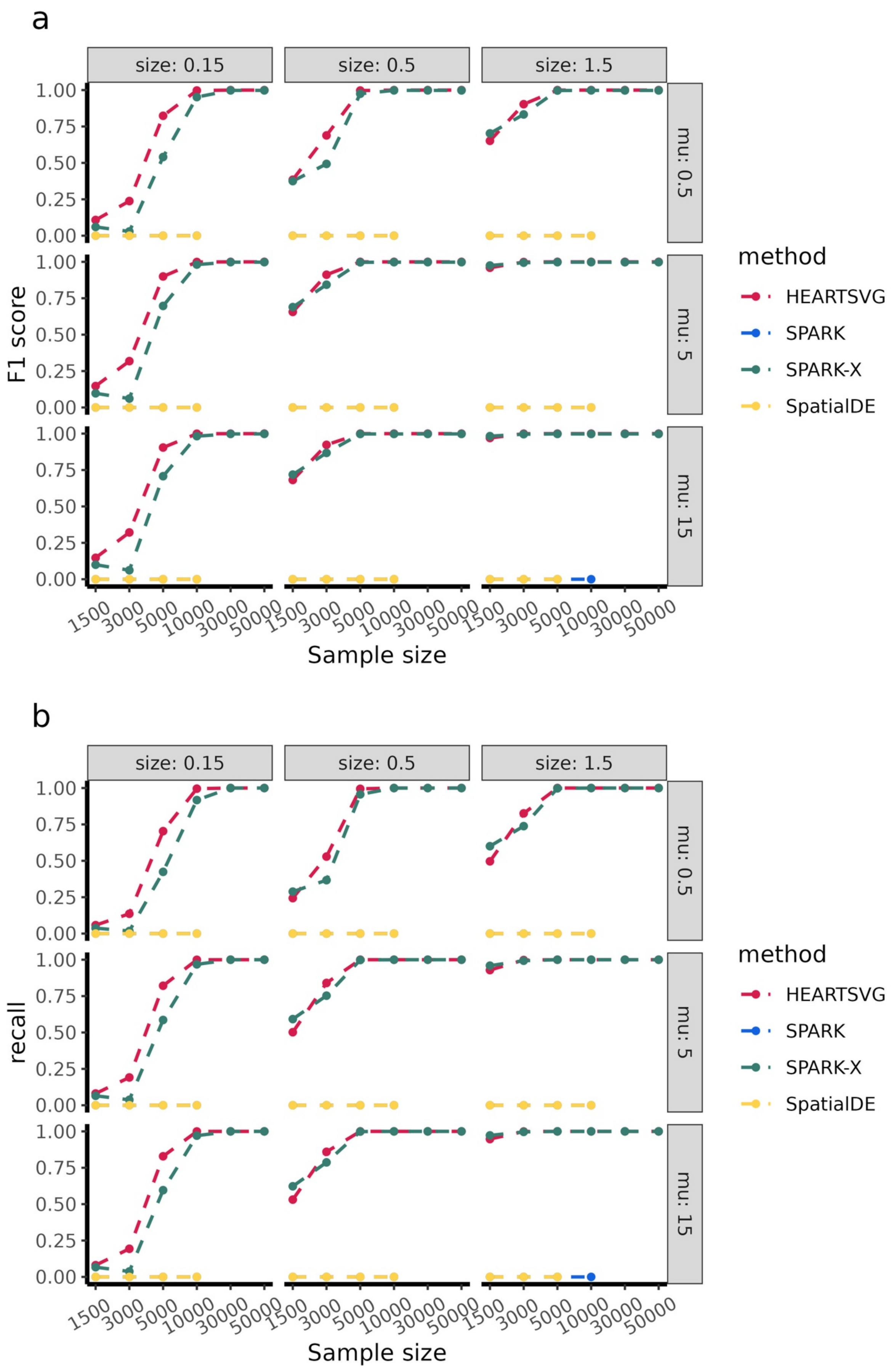

c

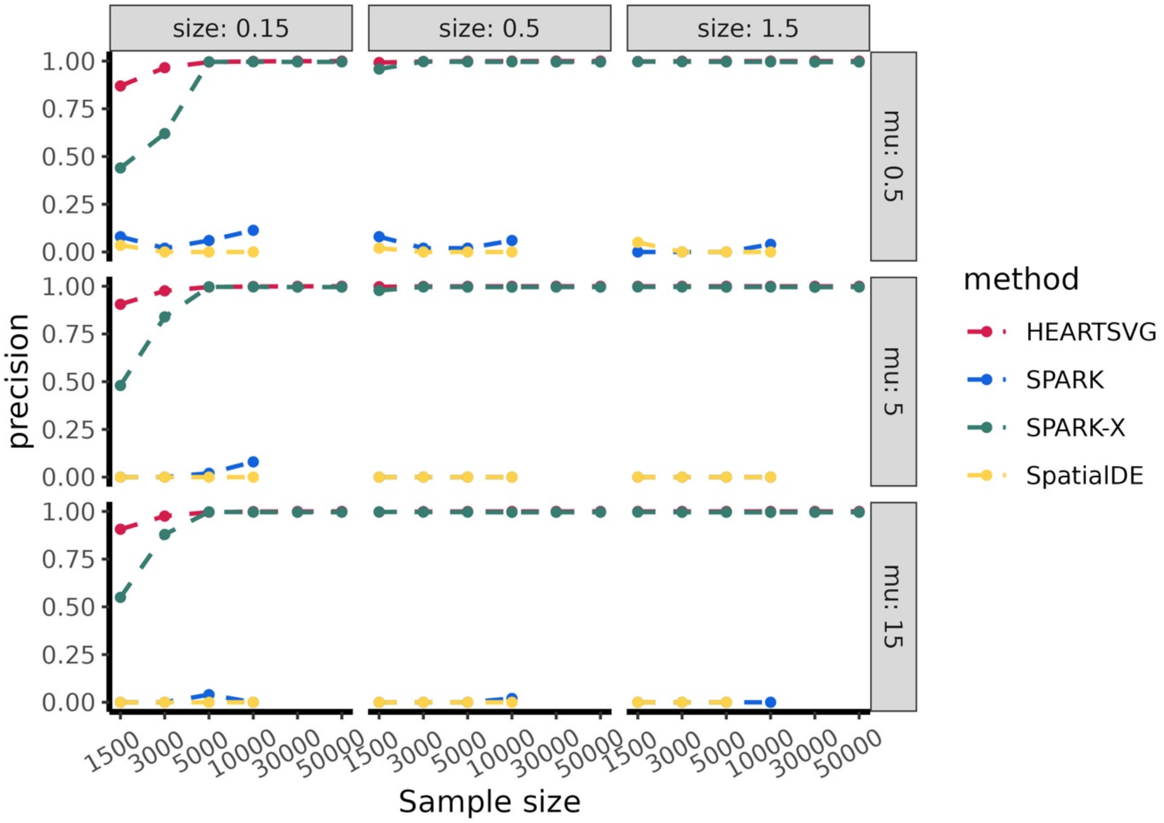
size: 0.15
size: 0.5
size: 1.5
mu: 0.5
mu: 5
mu: 15
precision
1.00
0.75
0.50
0.25
0.00
1500
3000
5000
10000
30000
50000
Sample size
method
HEARTSVG
SPARK
SPARK-X
SpatialDE

d

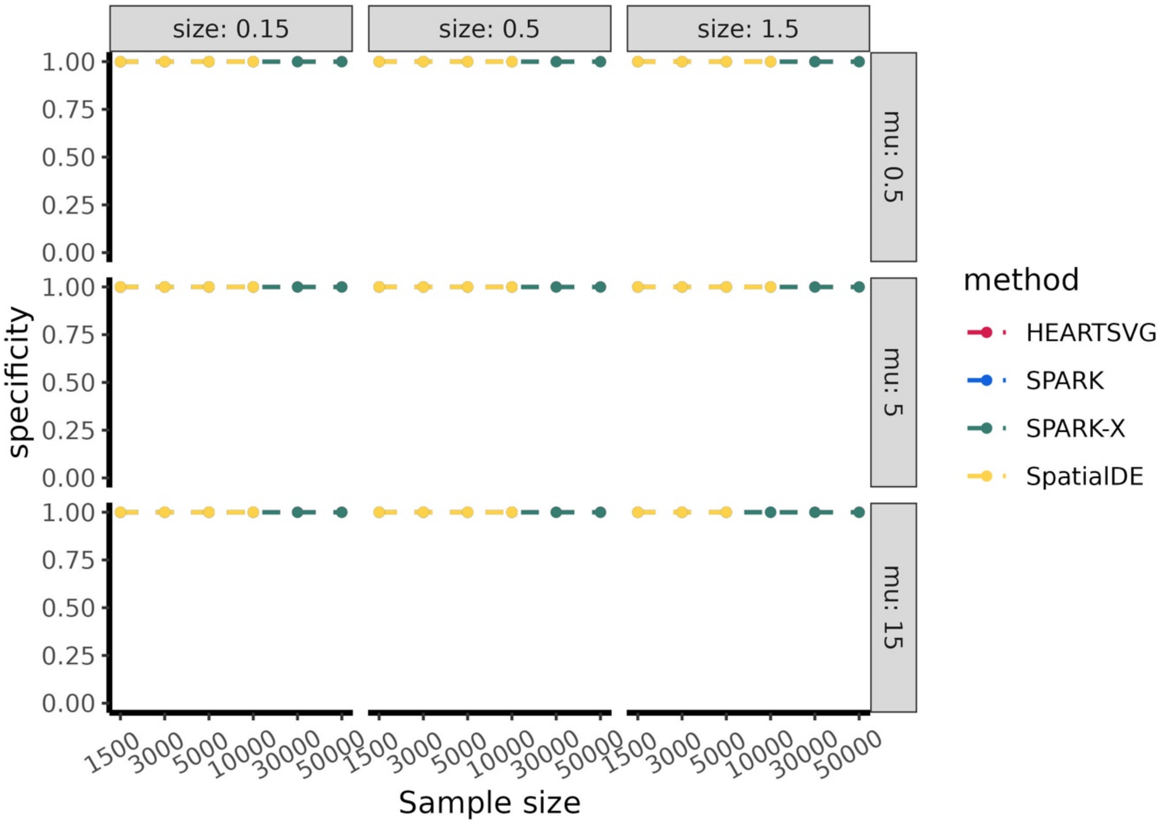
size: 0.15
size: 0.5
size: 1.5
mu: 0.5
mu: 5
mu: 15
specificity
1.00
0.75
0.50
0.25
0.00
1500
3000
5000
10000
30000
50000
Sample size
method
HEARTSVG
SPARK
SPARK-X
SpatialDE

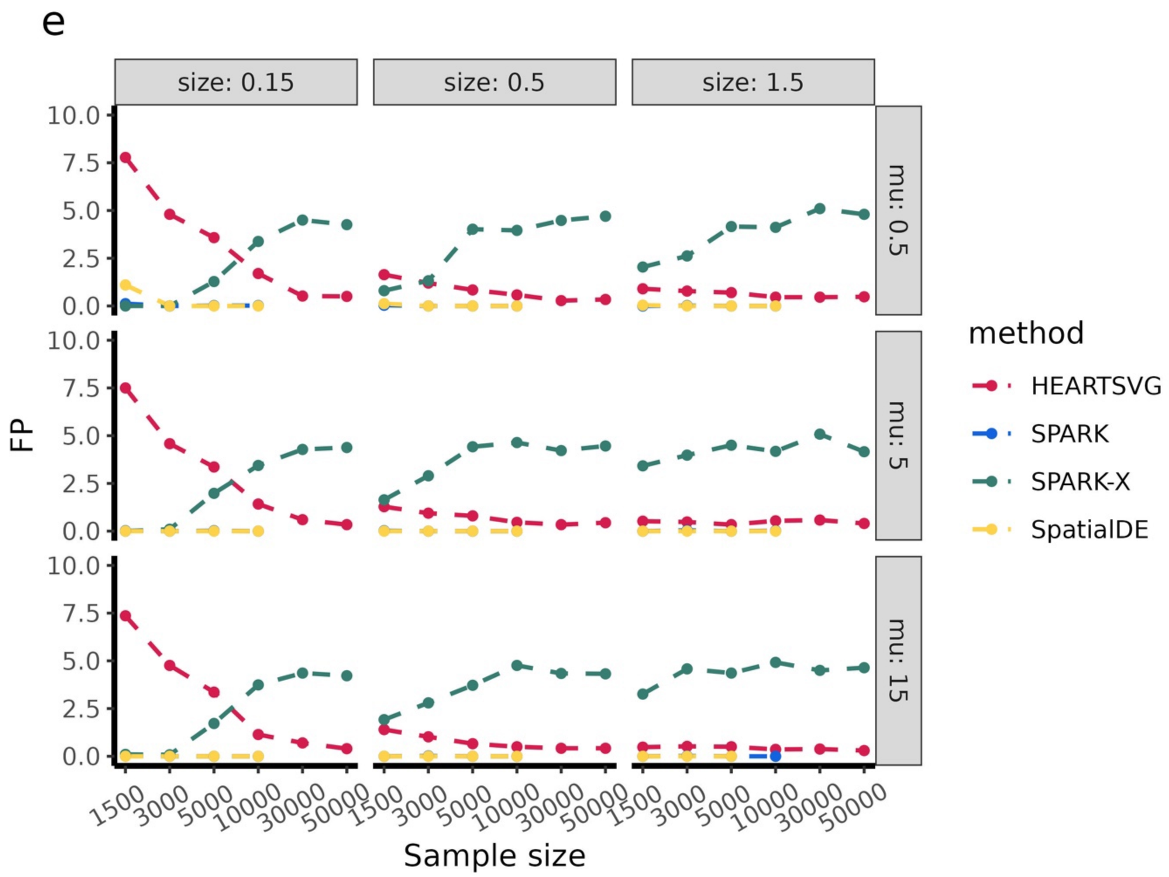


**Figure S2** Simulation results of streak pattern. **(a)** F1 score, (b) recall, **(c)** precision, **(d)** specificity, **(e)** FP.

**All simulation results of gradient pattern**

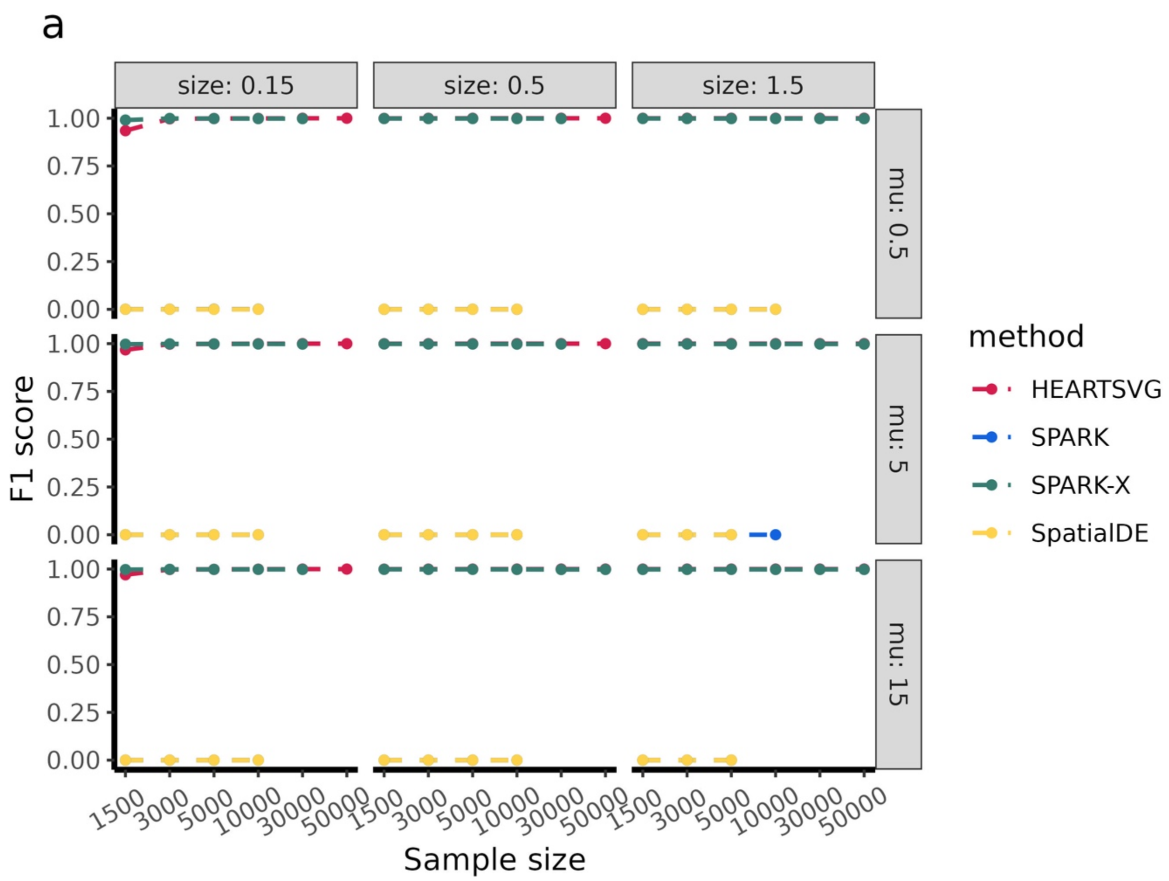


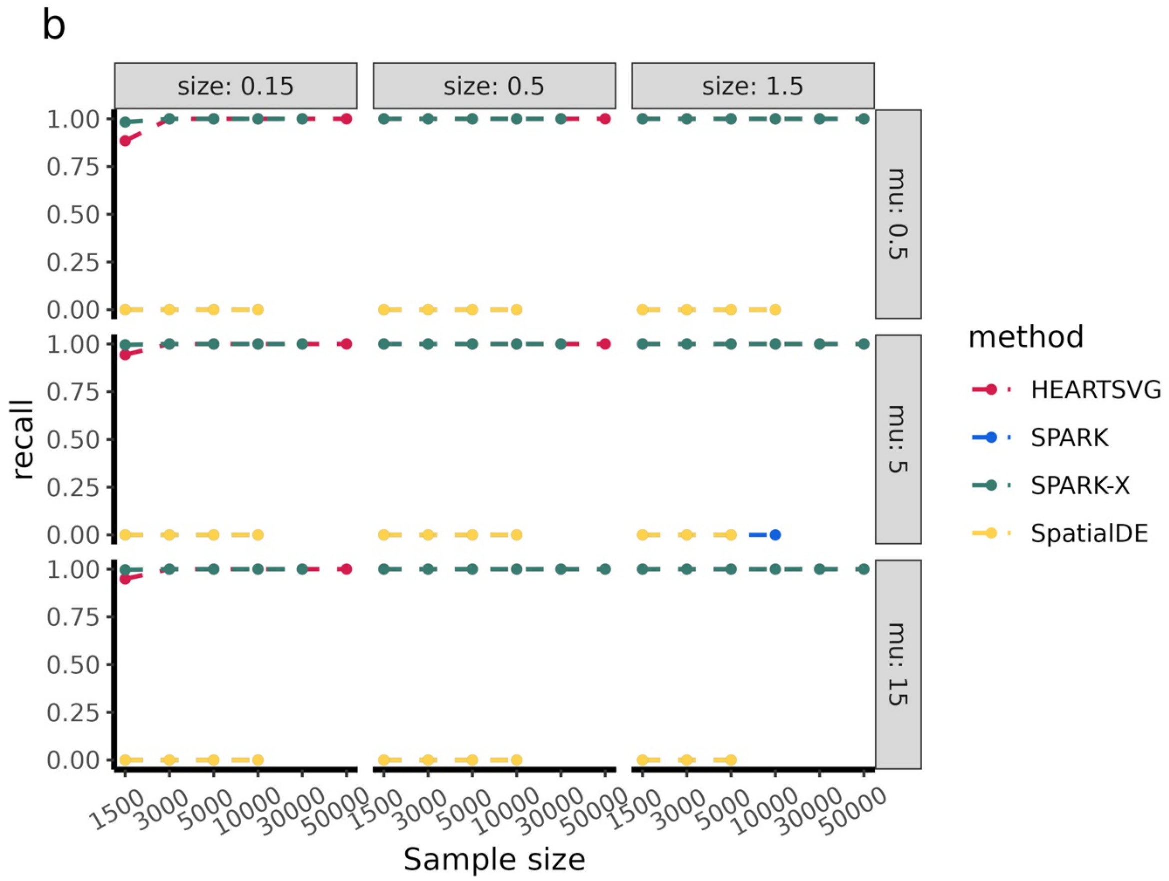

c

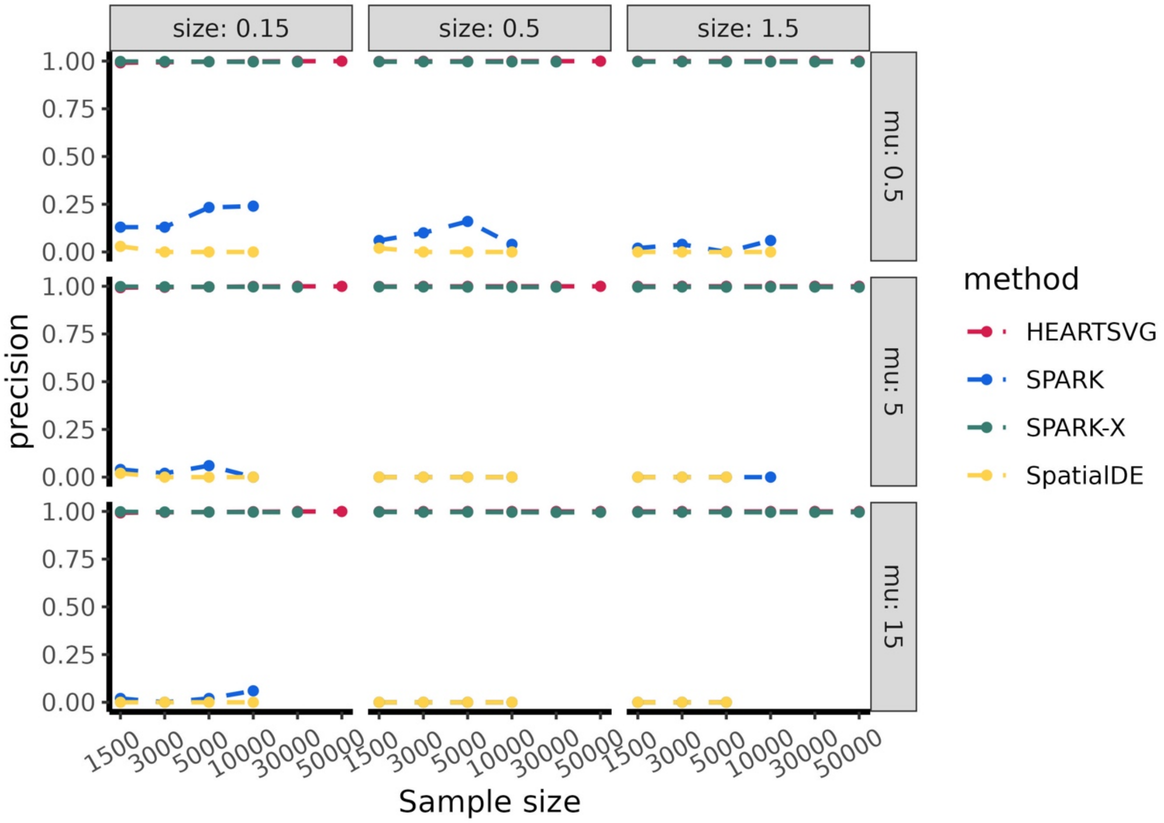
size: 0.15
size: 0.5
size: 1.5
mu: 0.5
mu: 5
mu: 15
precision
1.00
0.75
0.50
0.25
0.00
1500
3000
5000
10000
30000
50000
Sample size
method
HEARTSVG
SPARK
SPARK-X
SpatialDE

d

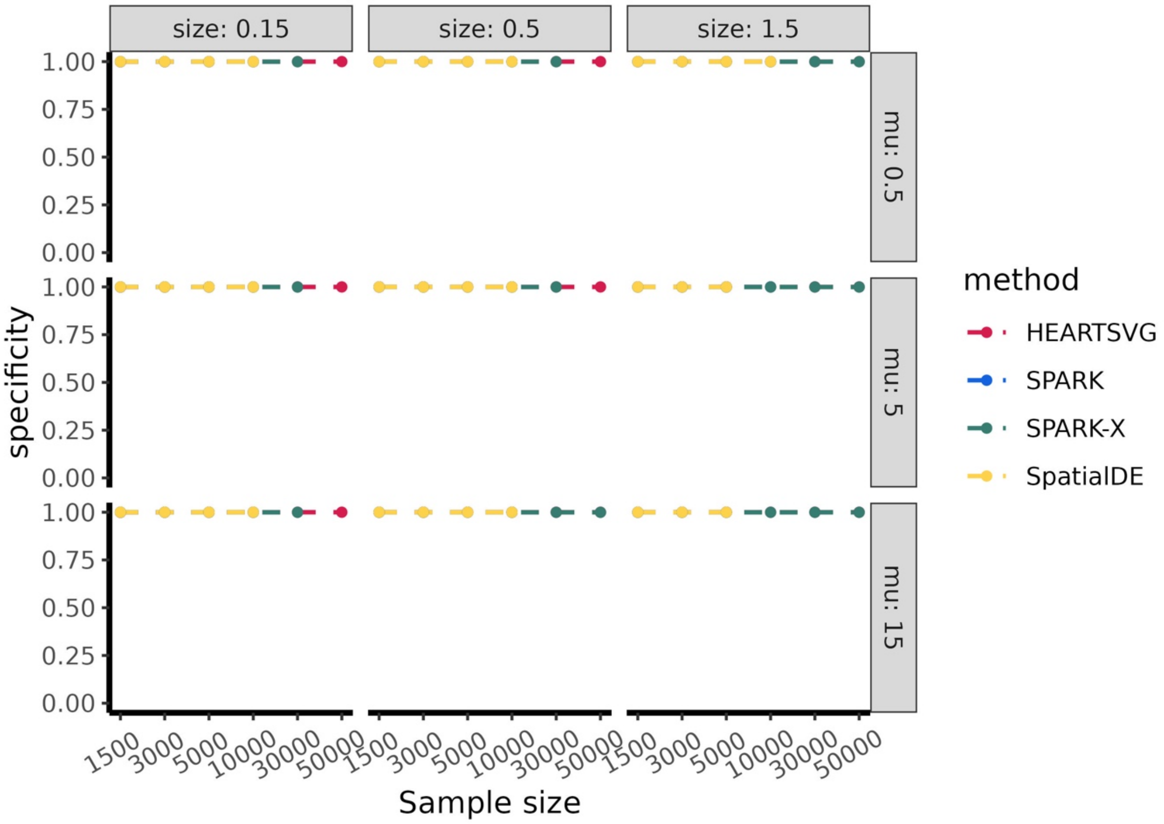
size: 0.15
size: 0.5
size: 1.5
mu: 0.5
mu: 5
mu: 15
specificity
1.00
0.75
0.50
0.25
0.00
1500
3000
5000
10000
30000
50000
Sample size
method
HEARTSVG
SPARK
SPARK-X
SpatialDE

e

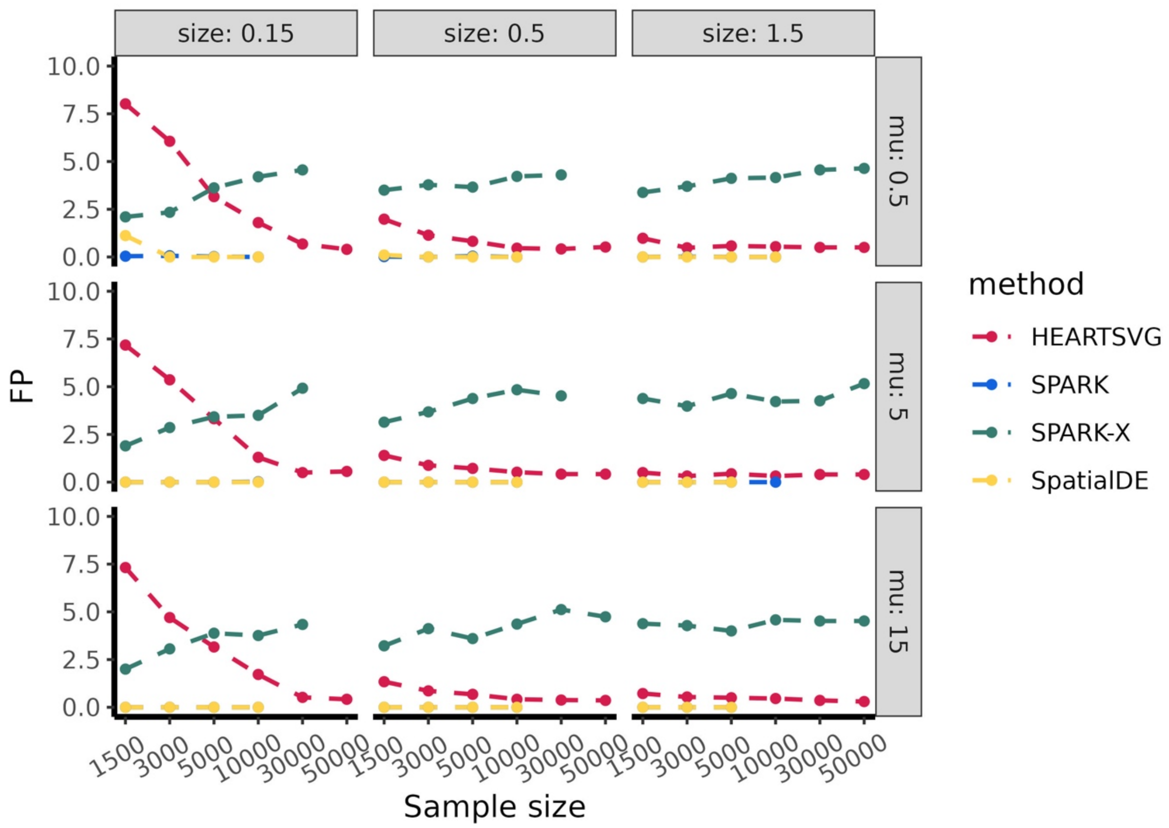


**Figure S3** Simulation results of gradient pattern. **(a)** F1 score, **(b)** recall, **(c)** precision, **(d)** specificity, **(e)** FP.

##### 1.2 Simulation 2 additional figures

Because the uncertainty surrounding the number of SVGs in real data is unknown, we generated simulation datasets with varying percentages (from 0% to 50%) of SVGs and sample sizes (from 3000 to 10,000) in three representative spatial patterns. We also compared the performance of HEARTSVG and SPARK-X using $F_1$ score, and false positive (FP). The paper presented the simulation scenario with ZINB parameters ($size = 0.5, mu = 0.5$) and moderate sample size (n=5000). The complete simulation results were shown in this supplementary.

**Hotspot pattern with varying percentages of SVGs**

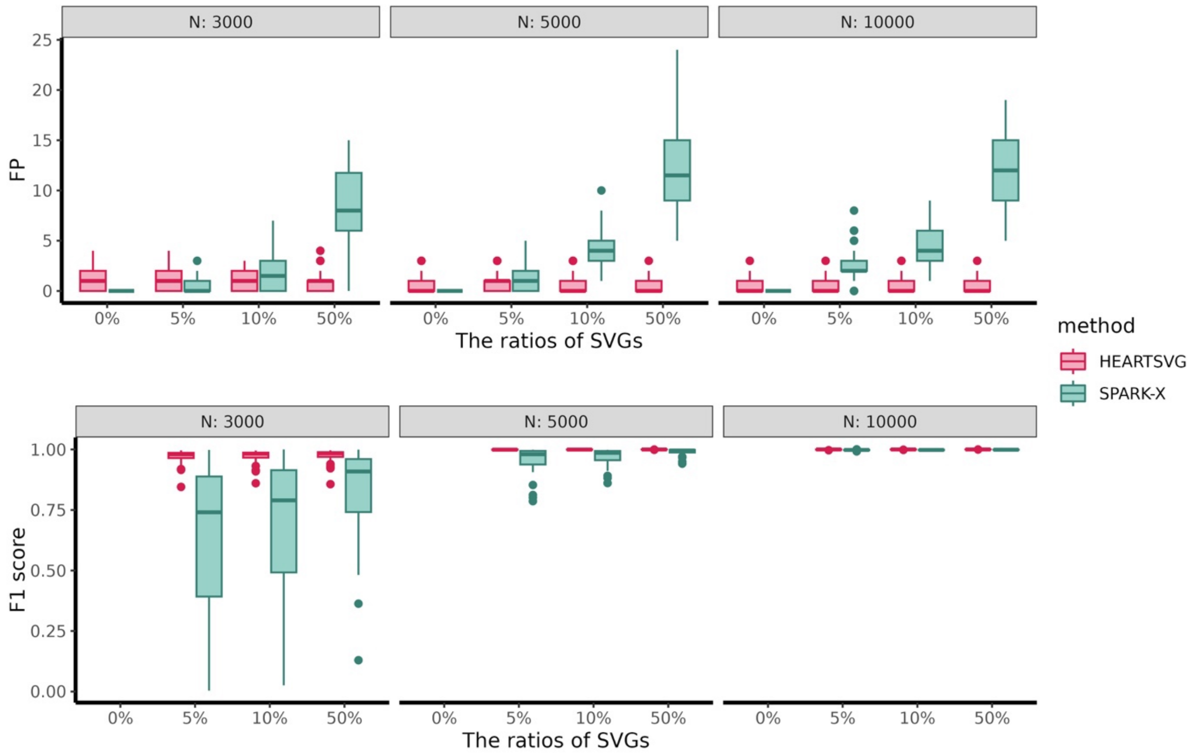


**Figure S4** Simulation2 results of hotspot pattern, F1 score and FP. In each boxplot, the lower hinge, upper hinge, and center line represent the 25th percentile (first quartile), 75th percentile (third quartile), and 50th percentile (median value), respectively. Whiskers extend no further than ±1.5 times the inter-quartile range. Data beyond the end of the whiskers are considered outliers and are plotted individually.

**Streak pattern with varying percentages of SVGs**

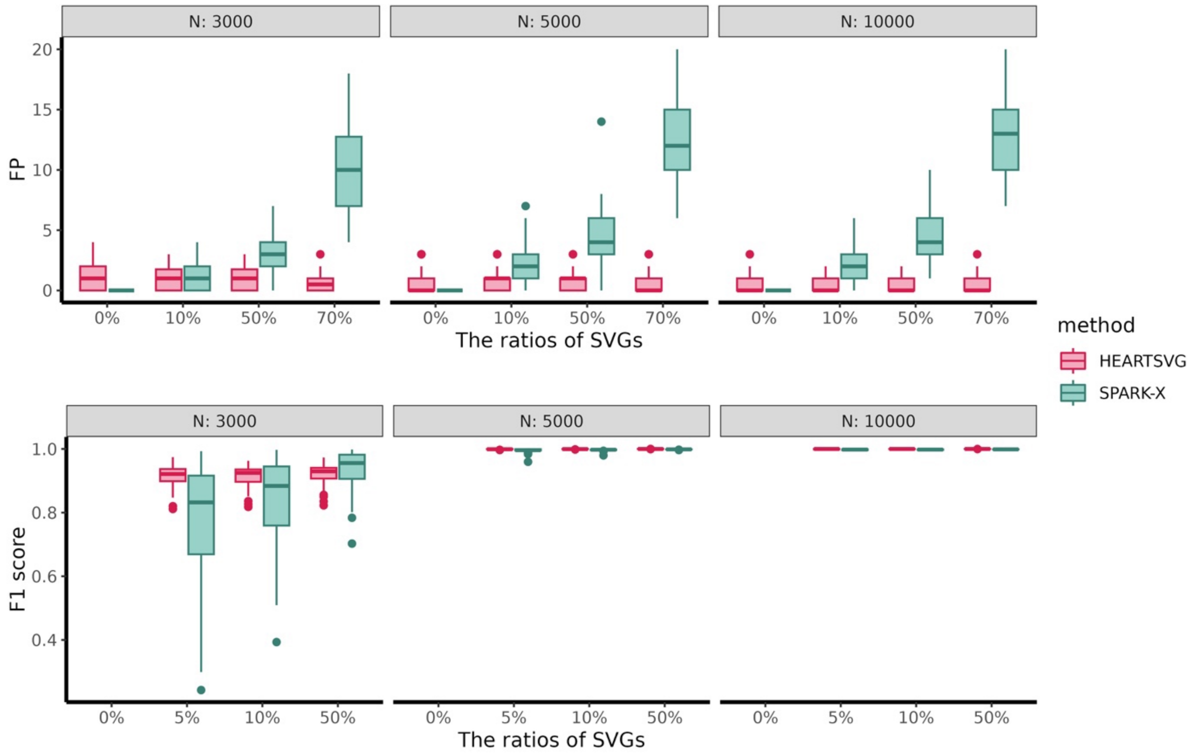


**Figure S5** Simulation2 results of streak pattern, F1 score and FP. In each boxplot, the lower hinge, upper hinge, and center line represent the 25th percentile (first quartile), 75th percentile (third quartile), and 50th percentile (median value), respectively. Whiskers extend no further than ±1.5 times the inter-quartile range. Data beyond the end of the whiskers are considered outliers and are plotted individually.

**Gradient pattern with varying percentages of SVGs**

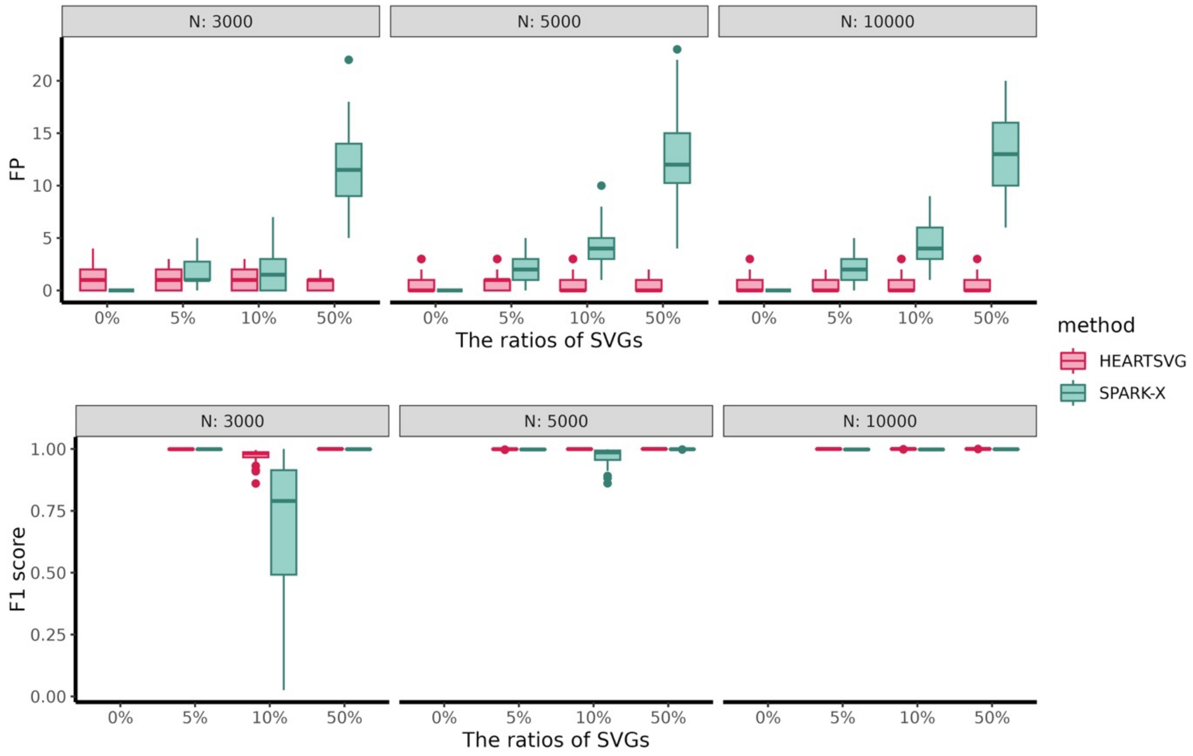


**Figure S6** Simulation2 results of gradient pattern, F1 score and FP. In each boxplot, the lower hinge, upper hinge, and center line represent the 25th percentile (first quartile), 75th percentile (third quartile), and 50th percentile (median value), respectively. Whiskers extend no further than ±1.5 times the inter-quartile range. Data beyond the end of the whiskers are considered outliers and are plotted individually.

##### 1.3 More comparison with scGCO in lowly sparse simulation data

To ensure a fair comparison between HEARTSVG and scGCO, we also included simulated data with lower sparsity. As in previous simulations, we simulated spatial locations (300, and 500 cells/spots) through a random-point-pattern Poisson process. We generated 2,000 genes in the simulated data, consisting of 1,000 SVGs and 1,000 non-SVGs. We simulated gene expressions for four distributions: negative binomial (NB), zero-inflated negative binomial (ZINB), Poisson, and zero-inflated Poisson (ZIP). The simulation setings are set as follows.

Table **S1** Simulation settings of simulated data with lower sparsity.

| | Non-SVG and SVG of non-marked cells/spots | | | SVG of marked cells/spots | | |
|---|---|---|---|---|---|---|
| | zero proportion | mu (NB, ZINB) / lambda (Poisson, ZIP) | size | zero proportion | mu (NB, ZINB) / lambda (Poisson, ZIP) | size |
| NB | 0.5 | 0.5 | 1.5 | 0.364 | 1.5 | 1.5 |
| ZINB | 0.644 | 1.5 | 5 | 0.228 | 4.5 | 5 |
| Poisson | 0.607 | 0.5 | - | 0.223 | 1.5 | - |
| ZIP | 0.654 | 2 | - | 0.202 | 6 | - |

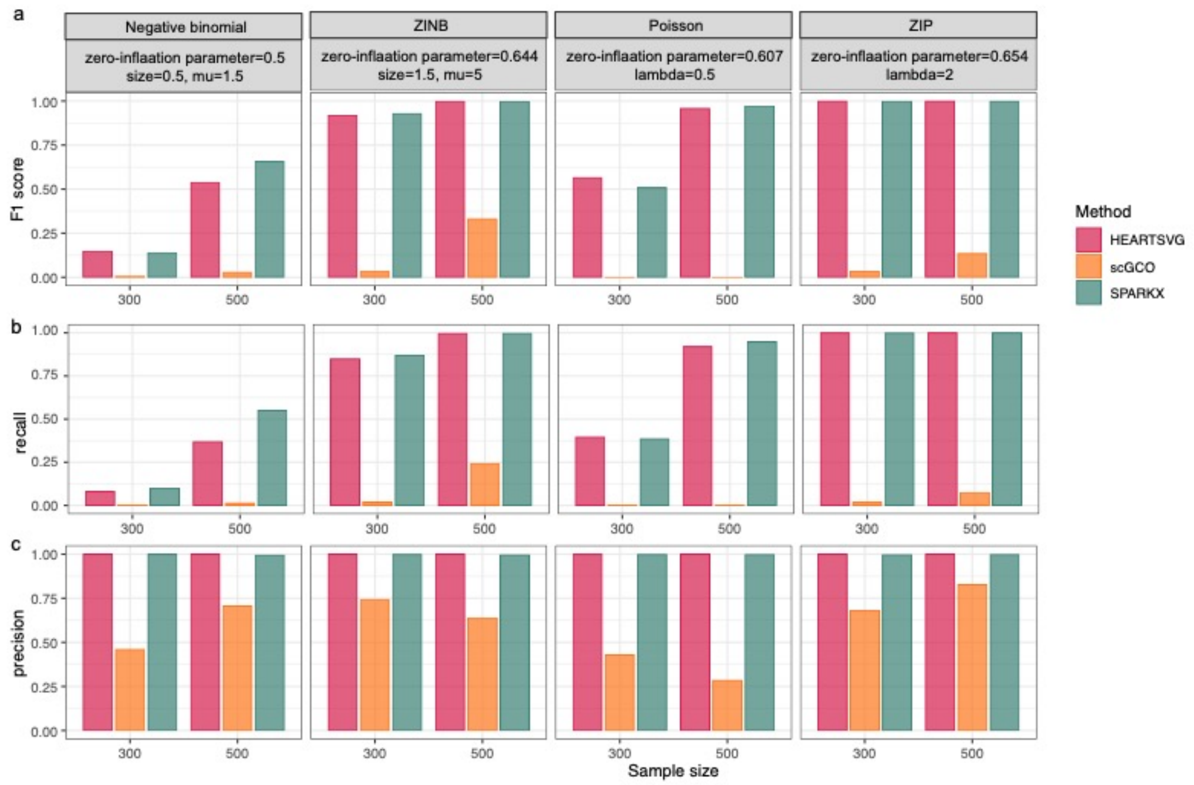


**Figure S7** Simulation results of hotspot pattern with low sparsity and large pattern size, (**a**) F1 score, (**b**) recall, (**c**) precision.

### 2. Methods

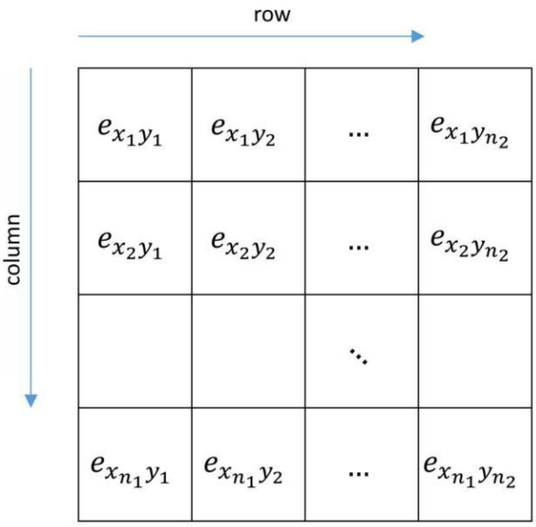


**Figure S8** Example of ST data.

##### 2.1 The following demonstrates the rationality of HEARTSVG.

For a gene $g$ without the spatial pattern in the ST data, its expression count $e$ is independent of its coordinates $(x, y)$. Define $f(x, y, e) = g(e|\boldsymbol{\theta})$, the gene $g$'s expression $e$ of each spot is only dependent on the parameters $\boldsymbol{\theta}$ (e.g., $\boldsymbol{\theta}$ could be the $\lambda$ of the Poisson distribution, $(\alpha, \beta)$ of the Gamma distribution, etc.).

That is, the gene $g$'s expression counts $e$ of the SRT data are independent and identically distributed random variables (Abbreviated as iid random variables).

We assume that every row has $n_2$ elements, every column has $n_1$ elements and expression matrix are a $n_1 * n_2$ matrix (Fig.**S8)**. Next, take the marginal expression series obtained by the semi-pooling process with the row direction and feature map $(1 * n_2)$ as an example. The gene g's expressions of the first row are denoted by $r_{x_1} = \left(e_{x_1y_1}, \ldots, e_{x_1y_j}, \ldots, e_{x_1y_{n_2}}\right)^{\boldsymbol{T}}$. The joint distribution of the first row is $f(r_{x_1}|\boldsymbol{\theta}) = f\left(e_{x_1y_1}, \ldots, e_{x_1y_j}, \ldots, e_{x_1y_{n_2}}\middle|\boldsymbol{\theta}\right) = \prod_{j=1}^{n_2} g\left(e_{x_1y_j}\middle|\boldsymbol{\theta}\right)$. Similarly, the expression vector and joint distribution of the second row are $r_{x_2} = \left(e_{x_2y_1}, \ldots, e_{x_2y_j}, \ldots, e_{x_2y_{n_2}}\right)^{\boldsymbol{T}}$ and $f\left(e_{x_2y_1}, \ldots, e_{x_2y_j}, \ldots, e_{x_2y_{n_2}}\middle|\boldsymbol{\theta}\right) = \prod_{j=1}^{n_2} g(e_{x_2y_j}|\boldsymbol{\theta})$, respectively. The expression vector and joint distribution of the $i$-th row are $r_{x_i} = \left(e_{x_i y_1}, \dots, e_{x_i y_j}, \dots, e_{x_i y_{n_2}}\right)^T$ and $f\left(e_{x_i y_1}, \dots, e_{x_i y_j}, \dots, e_{x_i y_{n_2}} \middle| \boldsymbol{\theta}\right) = \prod_{j=1}^{n_2} g(e_{x_i y_j}|\boldsymbol{\theta})$, respectively. Obviously, the joint distributions of each row's expressions are identically distributed. $r_{x_1}$ and $r_{x_2}$ have no overlapping elements. Hence, it is easy to prove that $f(r_{x_1}, r_{x_2}|\boldsymbol{\theta}) = f(r_{x_1}|\boldsymbol{\theta}) * f(r_{x_2}|\boldsymbol{\theta})$, which means $r_{x_1}$ and $r_{x_2}$ are independent and identically distributed random variables.

Similarly, we can prove that, $\forall\, i_1, i_2 = 1, \dots, n_1,\ and\ i_1 \neq i_2$, $r_{x_{i_1}}$ and $r_{x_{i_2}}$ are iid variables. By the similar derivation process as above, we can prove that the elements of the marginal expression series obtained by semi-pooling parameters with different parameters are iid variables. This is a very strong condition and is hard to verify empirically.

In practice, we assume that the expression counts of the non-SVG gene at a given location $(x_i, y_j)$ are independent of expressions at nearby locations. Therefore, we applied the Portmanteau test to test several autocorrelations of $r_t$ that are simultaneously at zero to determine whether the gene is a SVG. The null and alternative hypotheses are:

$$\mathrm{H_0}: \rho_1 =, \dots, = \rho_m = 0, \mathrm{H_A}: \exists\, k \in \{1, \dots, m\}, \rho_k \neq 0$$

To simplify the symbolic representation, we rewrite the subscript of the marginal expression series as $\mathbf{r} = (r_1, .., r_t, \dots, r_T)^T$, define the autocovariance of order $m$ as:

$\gamma_k = Cov(r_t, r_{t-k}) = Cov(r_t, r_{t+k})$, for all $k \geq 0$, and the $j$th order autocorrelation (ACF) as $\rho_k = \frac{\gamma_k}{\gamma_0}$.

If gene $g$ is non-SVG without a spatial pattern in ST data, our purpose is to test the null hypothesis: $\mathrm{H_0}: \rho_1 = \cdots = \rho_m = 0$.

The test statistic is defined as $Q_m = T \sum_{k=1}^{m} \hat{\rho}_k^2$ followed by chi-distribution with $m$ degree of freedom, where $\hat{\gamma}_k = \frac{1}{T-k} \sum_{t=1+k}^{T} (r_t - \bar{r})(r_{t-k} - \bar{r})$, $k = 0, \dots, T-1$, $\bar{r}$ is the mean of $\mathbf{r}$, and introduce $\hat{\rho}_k = \frac{\hat{\gamma}_k}{\hat{\gamma}_0}$. The $P$ value for testing the null hypothesis can be calculated by $p = \mathrm{P}(\chi^2(df = m) > Q_m \mid \mathrm{H_0} \text{ is true})$

**Stouffer's method**

We combined all four P values into a single P value by Stouffer's method. The Stouffer's statistic is defined as $z_{stouffer} = \sum_{i=1}^{4} \frac{z_i}{\sqrt{4}} \sim N(0,1)$, where $z_i = \Phi^{-1}(1 - p_i)$, $\Phi^{-1}(\cdot)$ is the inverse of the cumulative distribution function of a standard normal distribution. Hence, the combined p of four p-values is calculated by $\boldsymbol{p_c = 1 - \Phi(z)}$.

#### 2.2 Semi-pooling process

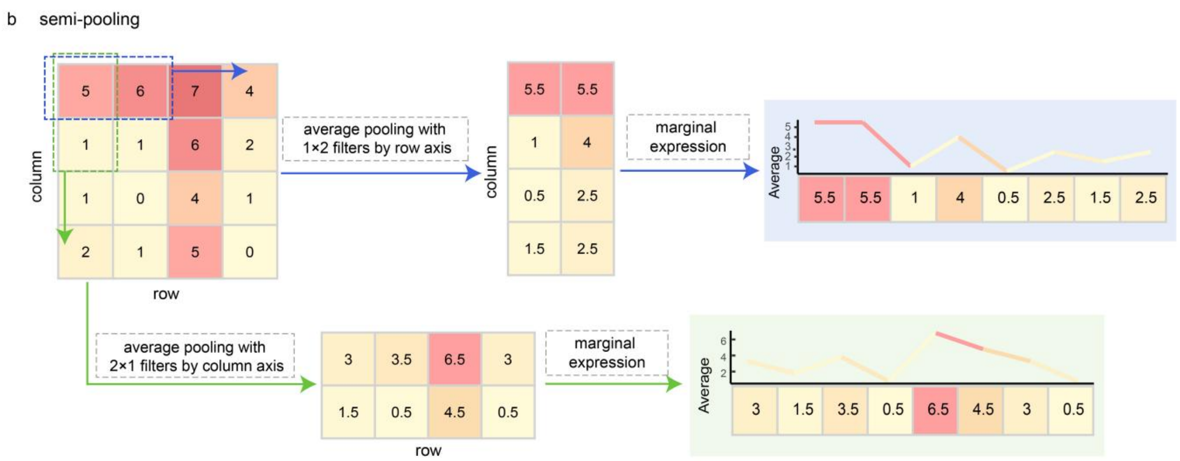


**Figure S9** Semi-pooling process

### 3. Application to colorectal cancer data by 10X Visium

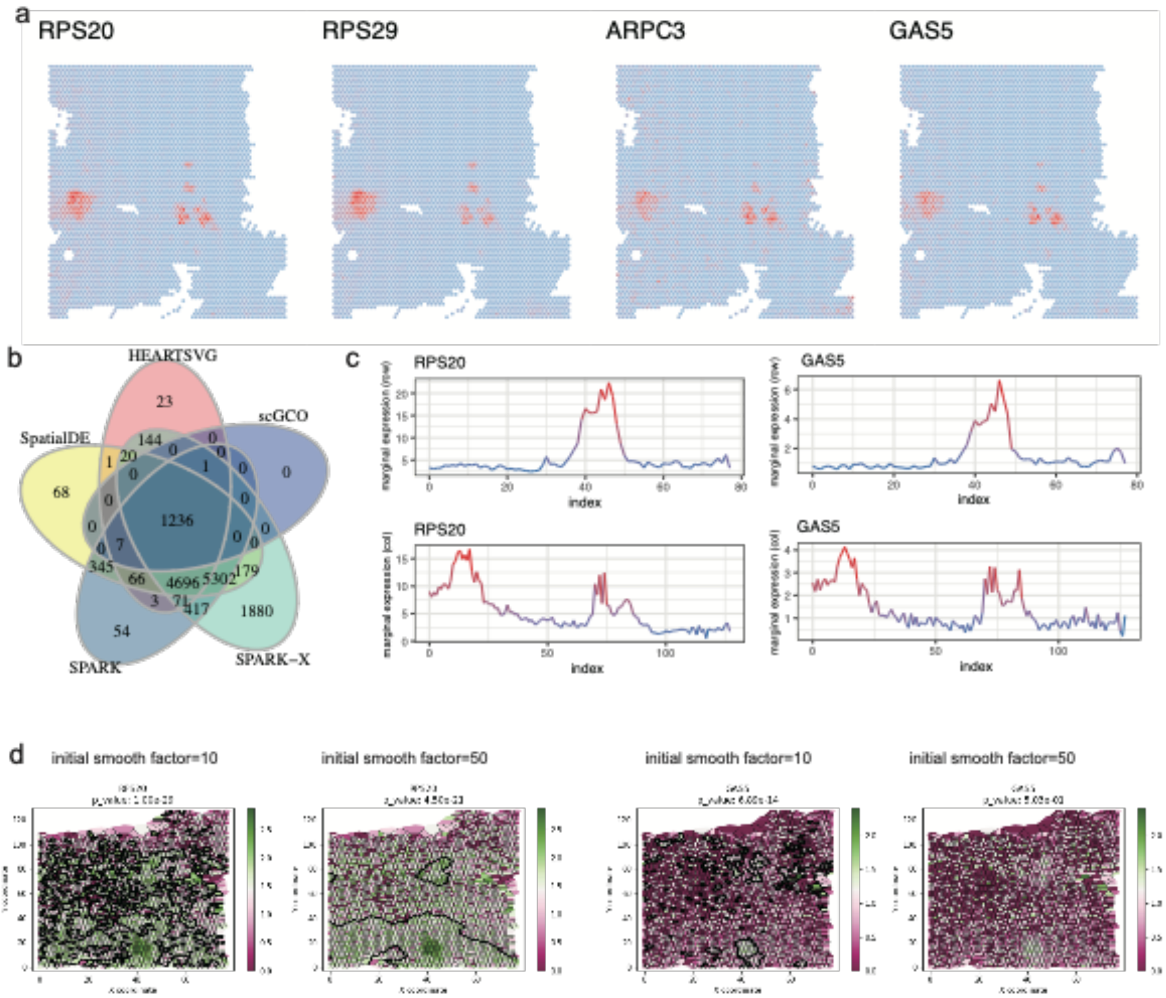


**Figure S10 scGCO missed SVGs (RPS29, ARPC3, GAS5) with clear spatial expression patterns comparing with other methods.** (**a**) Visualizations of spatial expressions of gene RPS20, RPS29, ARPC3, and GAS5. (**b**) Venn diagrams of SVGs in the colorectal cancer data identified by HEARTSVG, SpatialDE, SPARK, SPARK-X, and scGCO. (**c**) Marginal expression plots of gene RPS20, and GAS5 by HEARTSVG. (**d**) Visualizations of graph cuts by scGCO with different initial smooth factor of gene RPS20, and GAS5 by HEARTSVG

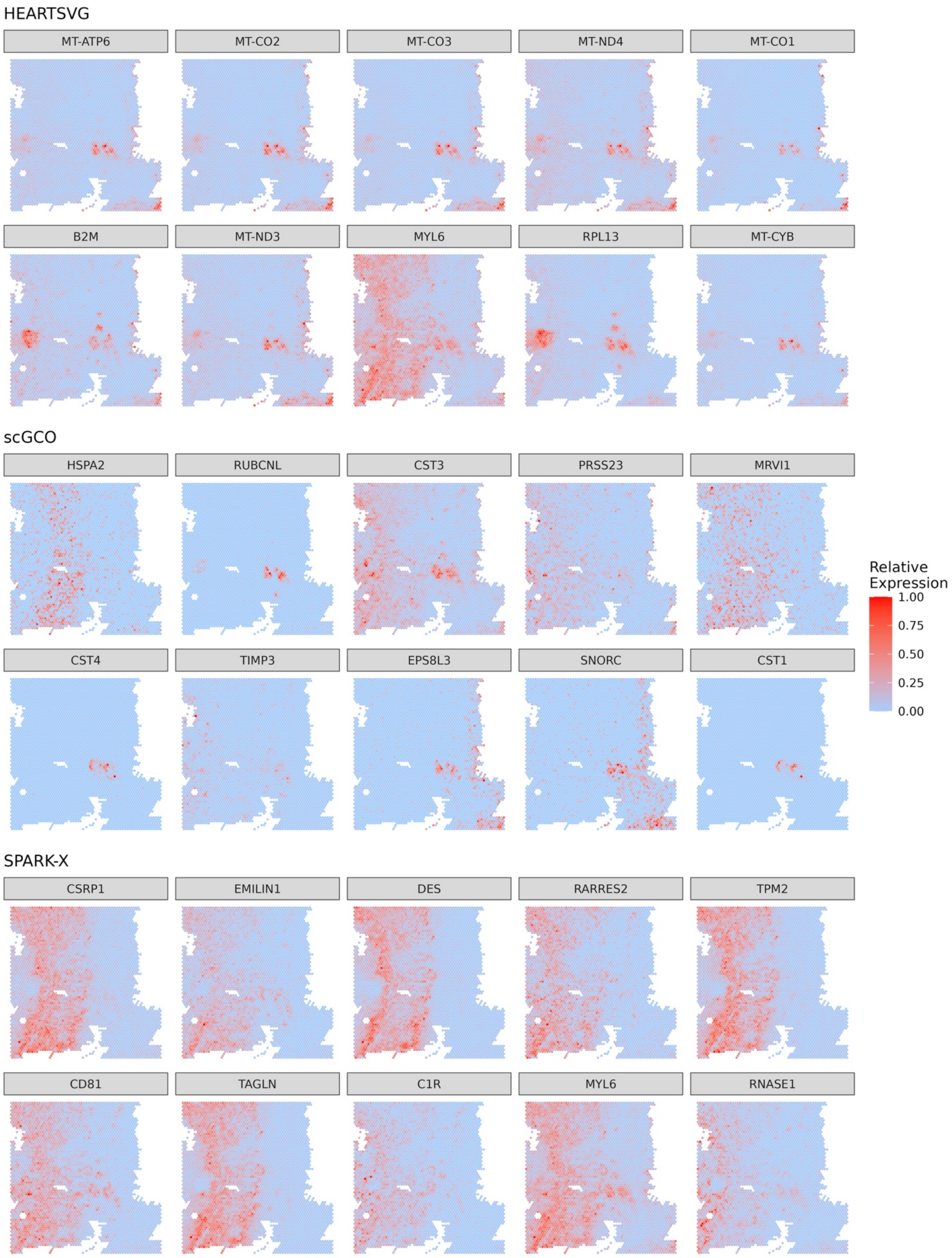
HEARTSVG
MT-ATP6
MT-CO2
MT-CO3
MT-ND4
MT-CO1
B2M
MT-ND3
MYL6
RPL13
MT-CYB
scGCO
HSPA2
RUBCNL
CST3
PRSS23
MRVI1
CST4
TIMP3
EPS8L3
SNORC
CST1
SPARK-X
CSRP1
EMILIN1
DES
RARRES2
TPM2
CD81
TAGLN
C1R
MYL6
RNASE1
Relative Expression
1.00
0.75
0.50
0.25
0.00

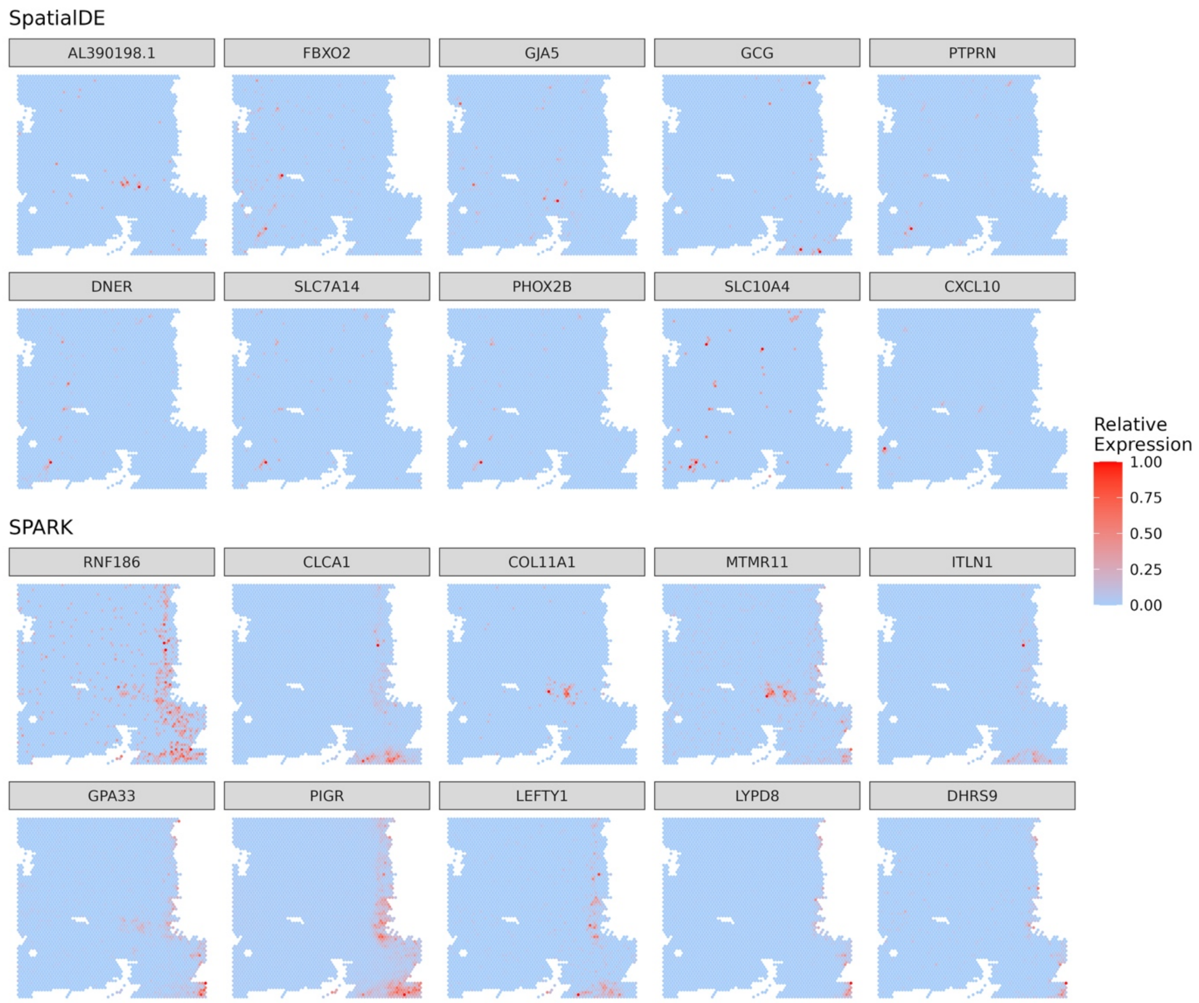


**Figure S11** Top 10 SVGs identified by each method. The top 10 genes identified by HEARTSVG, scGCO and SPARK-X showed stronger spatial expression patterns compared to SpatialDE and SPARK (Fig.S10-17). SpatialDE's top 10 selected SVGs exhibited minimal spatial patterns, whereas SPARK performed better than SpatialDE to a certain extent.

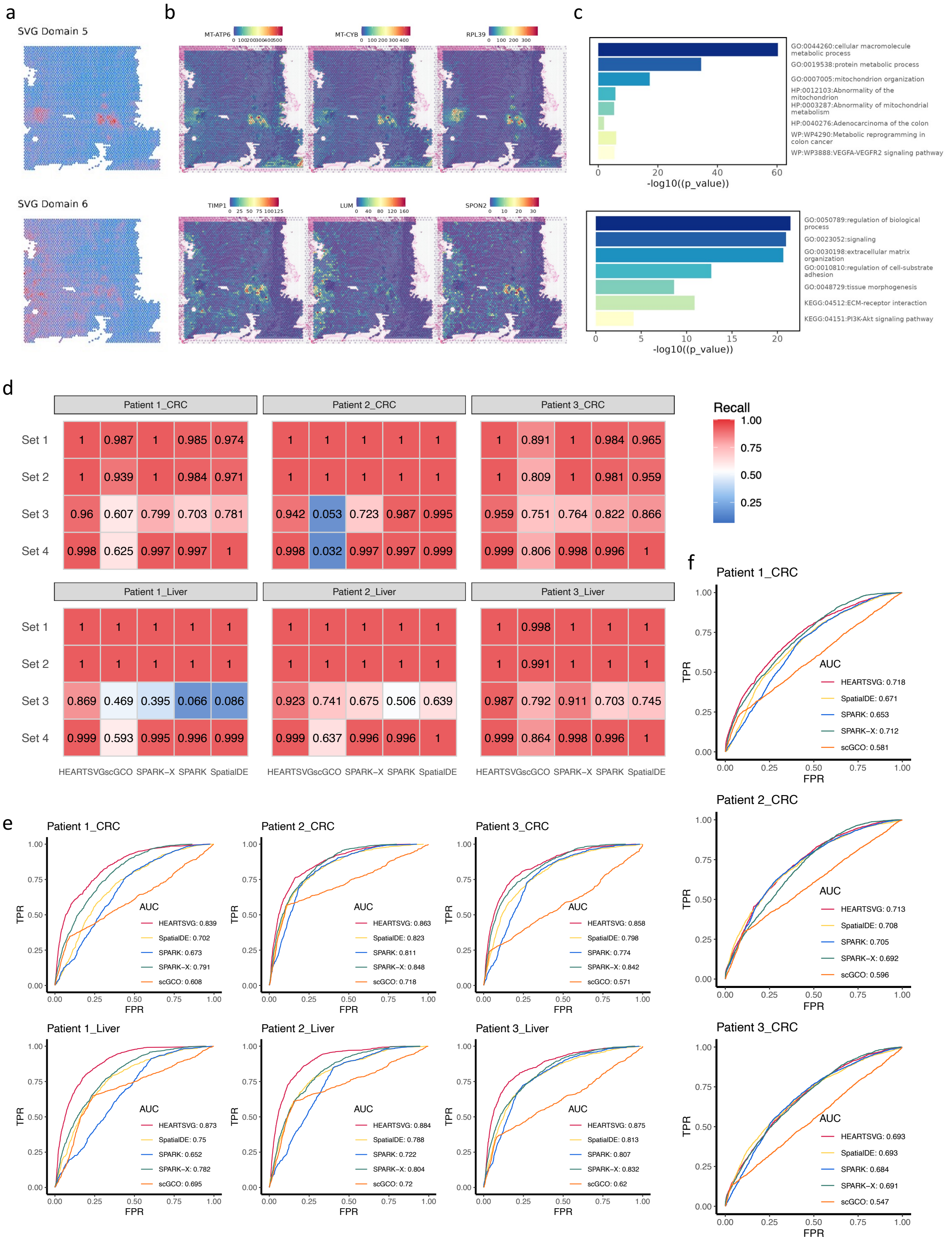


**Figure S12** **(a)** Predicted spatial domain 5,6 based on SVGs. **(b)** Representative genes of spatial domain 5,6. **(c)** Enrichment analysis of spatial domain 5,6. **(d)** The heatmap shows the comparison of recall values among four genesets on three colorectal cancer ST datasets and three corresponding liver metastasis ST datasets. Set 1 represents the true SVGs set, derived by selecting the top 500 overlaps between HEARTSVG and SPARK-X. Set 4 corresponds to the non-SVGs set, obtained by randomly rearranging the gene expressions within Set 1. Set 2 and Set 3 are generating by introducing noise to the true SVGs set and non-SVGs set, respectively. **(e)** ROC curves were used to assess the true positive rate (TPR) and false positive rate (FPR) of four different methods, using common gene modules of tumor microenvironments as gold standards for true spatially variable genes (SVGs). The figures consisted of six sub-figures representing three colorectal cancer ST datasets and three corresponding liver metastasis ST datasets. Each method was represented by a different colored line, and the area under the ROC curve (AUC) was calculated. **(f)** ROC curves were used to evaluate the TPR and FPR of four different methods, using consensus molecular markers of colorectal cancer subtypes as gold standards for true SVGs. The figures comprised three sub-figures corresponding to three colorectal cancer ST datasets.

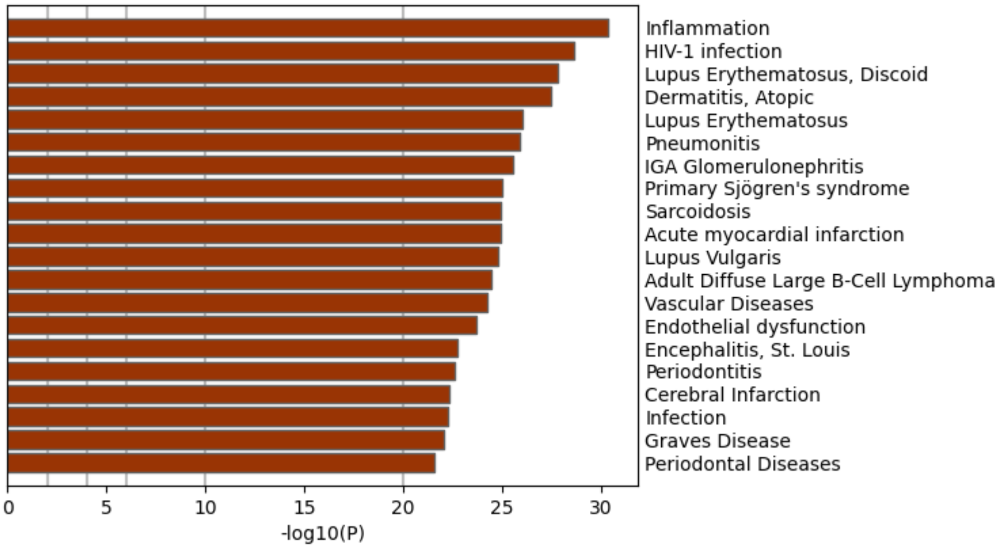


**Figure S13** Enrichment results of Spatial domain 1.

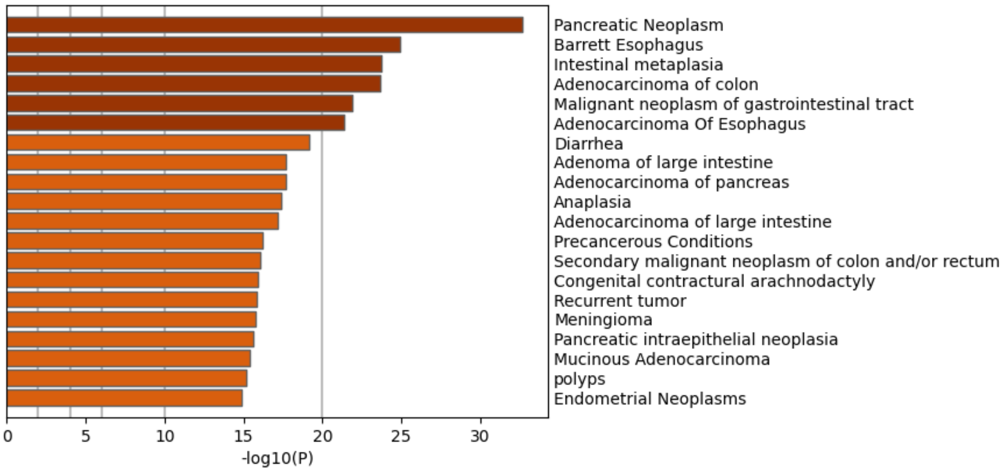


**Figure S14** Enrichment results of Spatial domain 2.

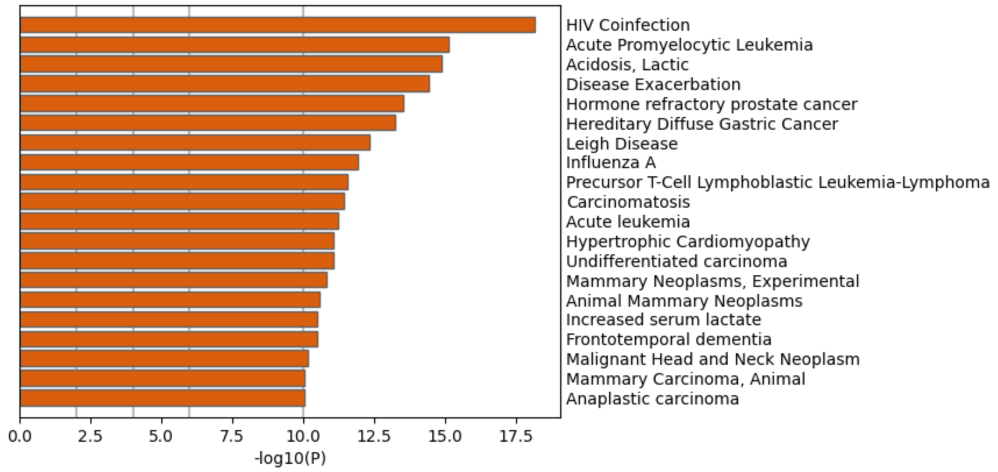


**Figure S15** Enrichment results of Spatial domain 3.

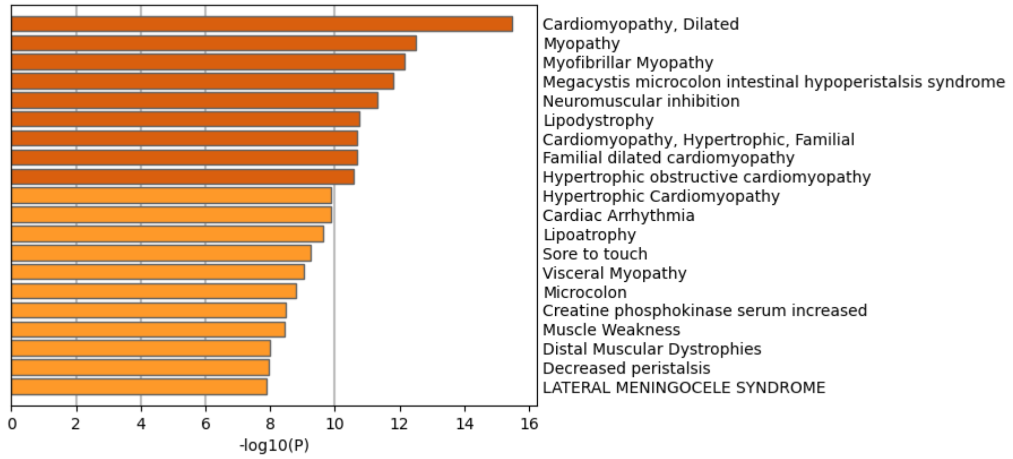


**Figure S16** Enrichment results of Spatial domain 4.

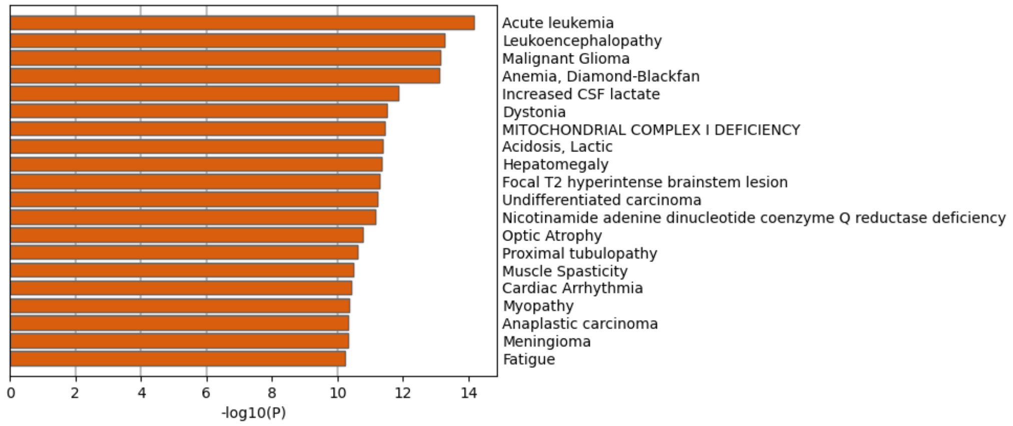


**Figure S17** Enrichment results of Spatial domain 5.

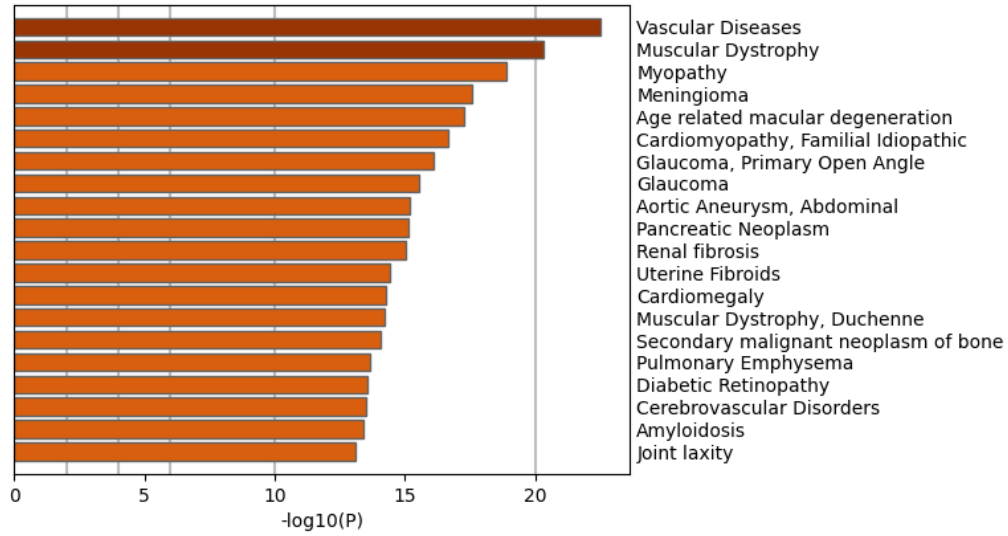


**Figure S18** Enrichment results of Spatial domain 6.

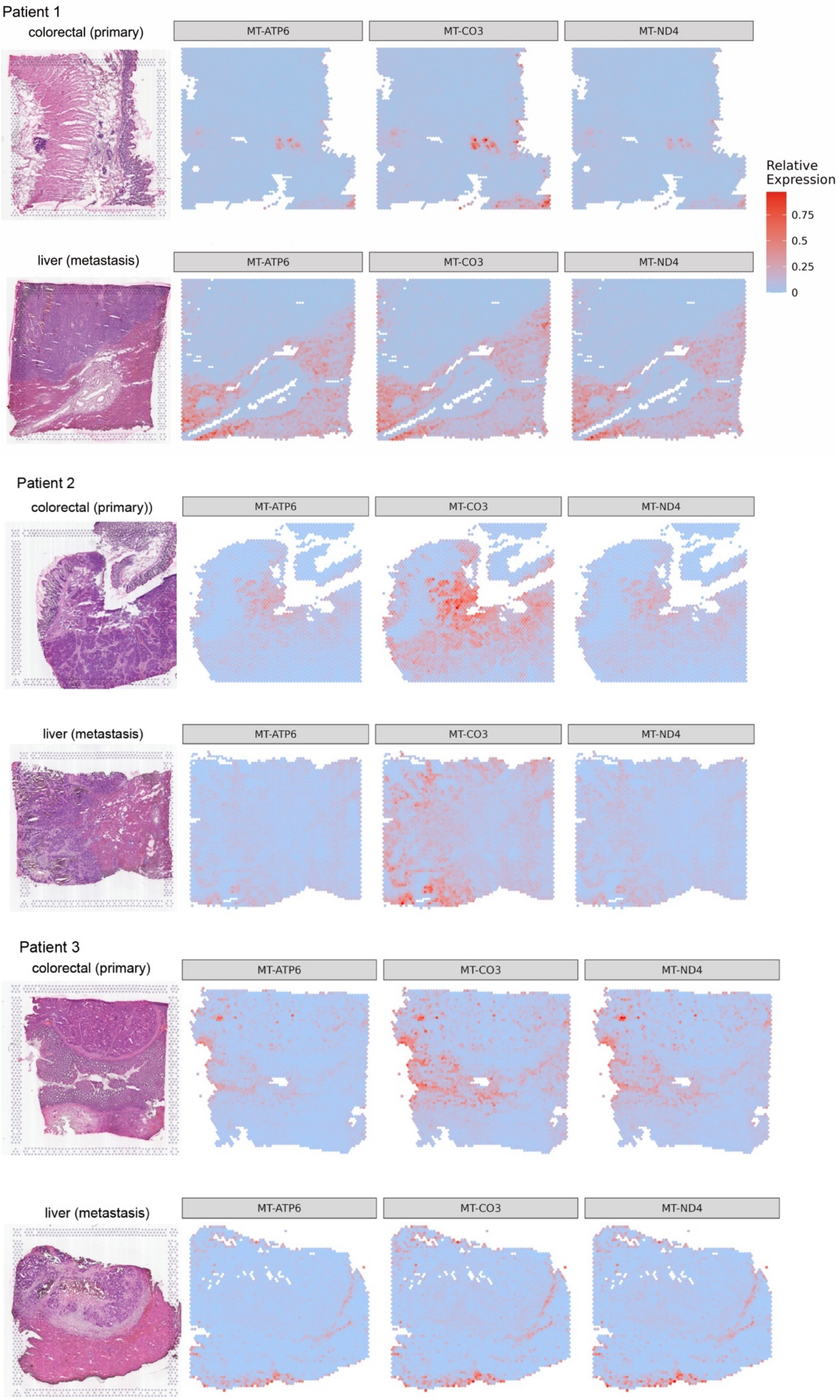


**Figure S19** . 'MT' genes in primary colorectal cancer tissue and liver metastasis cancer tissue.

### 4. Application to mouse cerebellum data by Slide-seqV2

Table **S2** Tissue-specificity enrichment analysis results of each method.

| | caudate | cerebellum | cerebral cortex | hippocampus | rectum | skin | SUM |
|---|---|---|---|---|---|---|---|
| HEARTSVG | 0 (0%) | 35 (87.5%) | 4 (10%) | 1 (2.5%) | 0 (0%) | 0 (0%) | 40 (100%) |
| scGCO | 0 (0%) | 24 (92.31%) | 2 (7.69%) | 0 (0%) | 0 (0%) | 0 (0%) | 26 (100%) |
| SPARK | 5 (5.1%) | 42 (42.86%) | 7 (7.14%) | 4 (4.08%) | 3 (3.06%) | 37.76(18.37%) | 98 (100%) |
| SPARK-X | 1 (2%) | 38 (76%) | 4 (8%) | 1 (2%) | 3 (6%) | 3 (6%) | 50 (100%) |
| SpatialDE | 0 (0%) | 44 (86.27%) | 4 (7.84%) | 2 (3.92%) | 1 (1.96%) | 0 (0%) | 51 (100%) |

Table **S3** Tissue-specificity enrichment analysis results of each method.

| gene | method | rank | adjusted p | |
|---|---|---|---|---|
| Car8 | HEARTSVG | 66 | 0.000 | *** |
| | scGCO | 2 | 4.46e-28 | *** |
| | SpatialDE | 37 | 0.000 | *** |
| | SPARK | 3 | 0.000 | *** |
| | SPARK-X | 585 | 0.050 | |
| Pcp2 | HEARTSVG | 38 | 0.000 | *** |
| | scGCO | 5 | 4.46e-28 | *** |
| | SpatialDE | 145 | 0.000 | *** |
| | SPARK | 110 | 0.000 | *** |
| | SPARK-X | 804 | 0.139 | |
| Pcp4 | HEARTSVG | 27 | 0.000 | *** |
| | scGCO | 4 | 4.46e-28 | *** |
| | SpatialDE | 48 | 0.000 | *** |
| | SPARK | 5 | 0.000 | *** |
| | SPARK-X | 2485 | 1.000 | |
| Calm1 | HEARTSVG | 6 | 0.000 | *** |
| | scGCO | 1044 | 0.501 | |
| | SpatialDE | 204 | 0.000 | *** |
| | SPARK | 25 | 4.64e-15 | *** |
| | SPARK-X | 49 | 4.33e-10 | *** |
| Calm2 | HEARTSVG | 8 | 0.000 | *** |
| | scGCO | 1031 | 0.501 | |
| | SpatialDE | 292 | 8.35e-14 | *** |
| | SPARK | 26 | 4.64e-15 | *** |
| | SPARK-X | 63 | 1.68e-08 | *** |
| Itm2b | HEARTSVG | 11 | 0.000 | *** |
| | scGCO | 1269 | 0.501 | |
| | SpatialDE | 116 | 0.000 | *** |
| | SPARK | 75 | 4.64e-15 | *** |
| | SPARK-X | 322 | 5.70e-03 | *** |

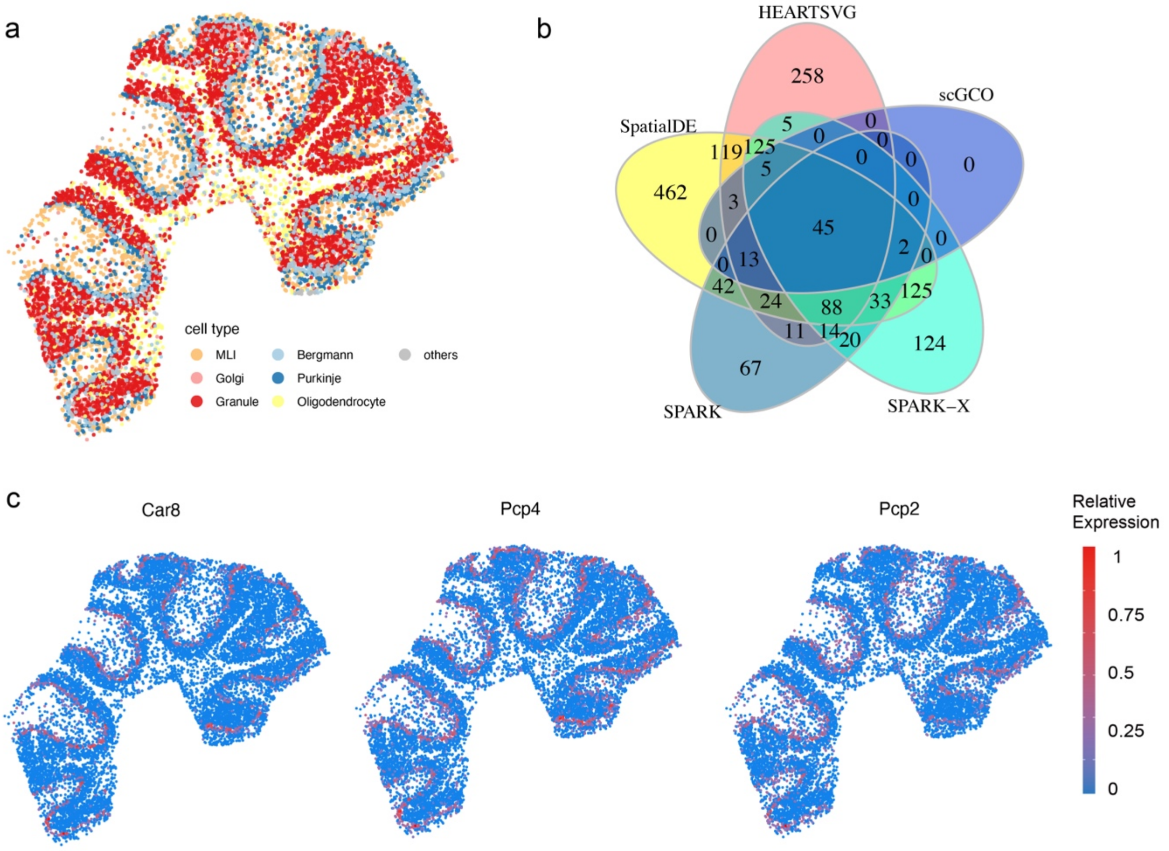


**Figure S20** **(a)** Visualization of unsupervised spatial clustering results. (**b**) Venn diagrams of SVGs in the mouse cerebellum data identified by HEARTSVG, SpatialDE, SPARK, SPARK-X, and scGCO. **(c)** Visualizations of marker genes of Purkinje cells in the mouse cerebellum data by Slide-seqV2.

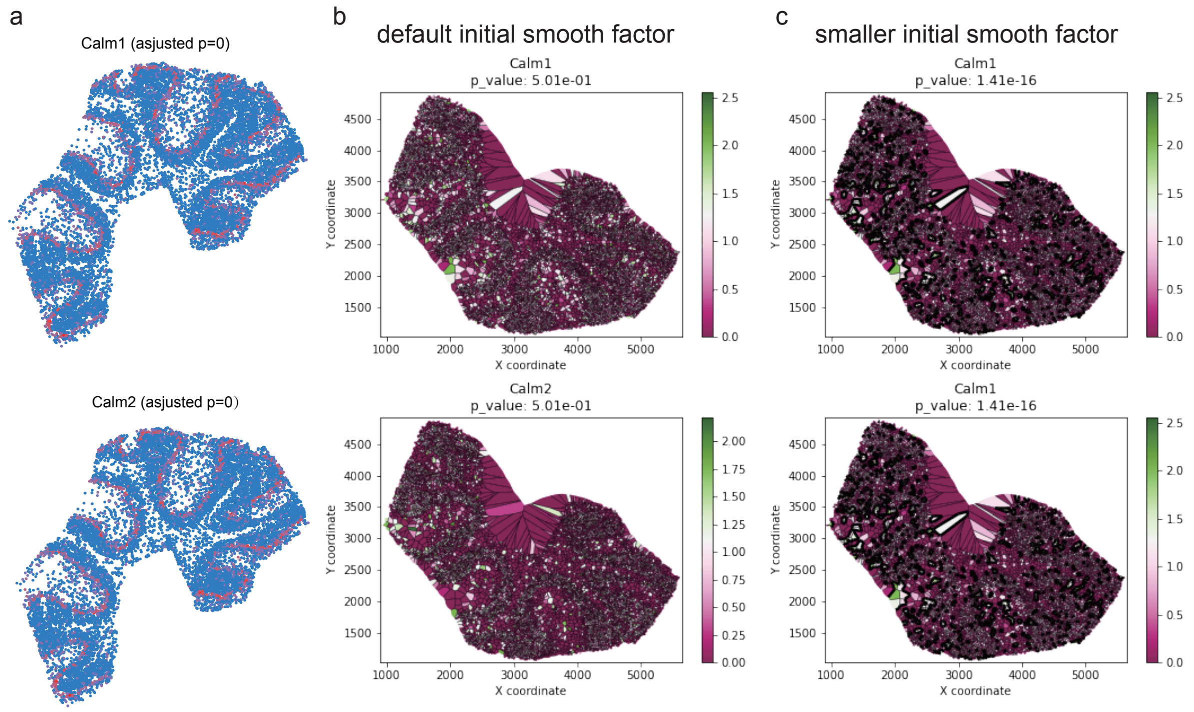


**Figure S21 scGCO missed SVGs (Calm1, Calm2) with clear spatial expression patterns comparing with other methods.** (**a**) Visualizations of spatial expressions of Calm1 and Calm2 in the in the Slide-seqV2 cerebellum data. (**b**) Visualizations of graph cuts by scGCO with default initial smooth factor of Calm1 and Calm2. (**c**) Visualizations of graph cuts by scGCO with smaller initial smooth factor of Calm1 and Calm2.

### 5. Application to mouse preoptic hypothalamus data by MERFISH

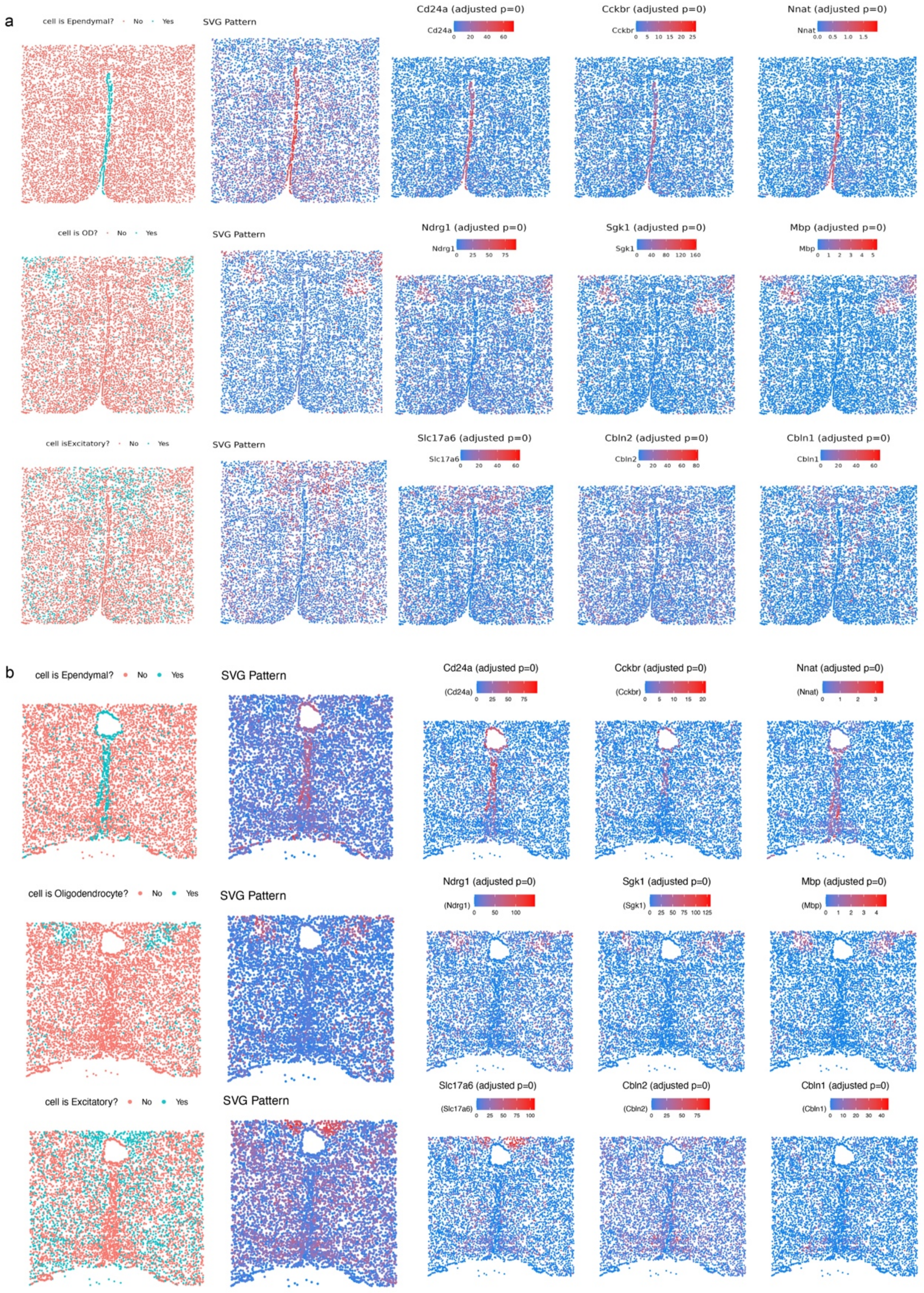


**Figure S22** (**a**) Visualizations of marker genes and cell type of the MERFISH data 1. (**b**) Visualizations of marker genes and cell type of the MERFISH data 2.

### 6. Application to mouse olfactory bulb data by HDST

a

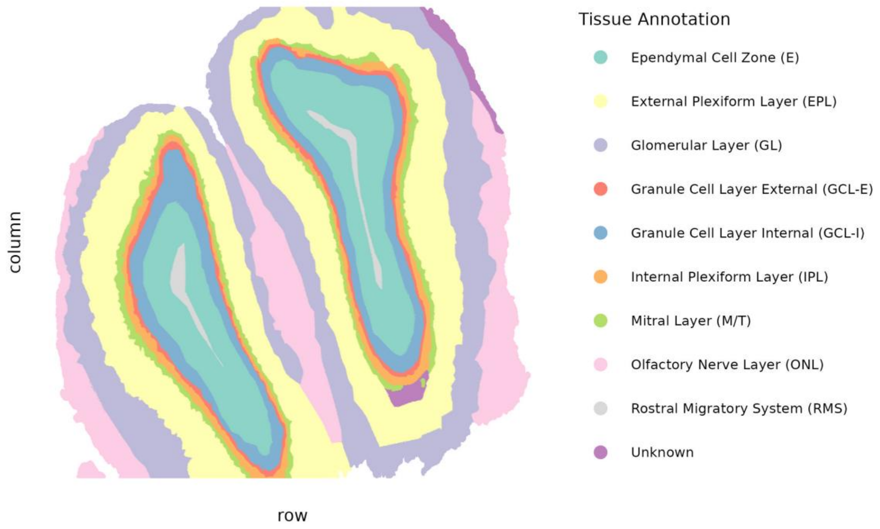


b

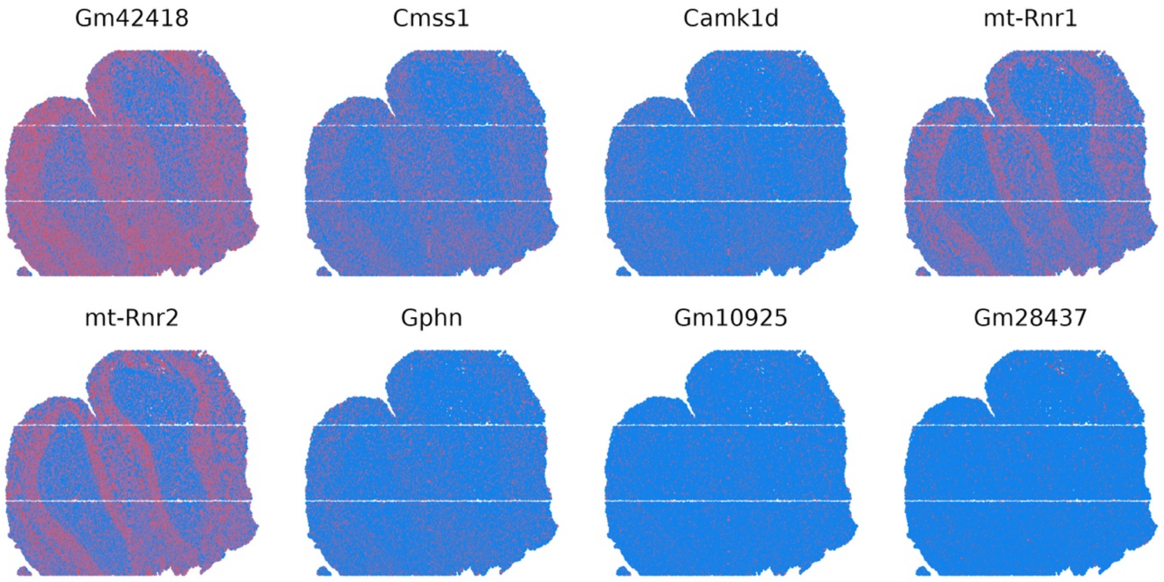


**Figure S23** **(a)** Cell annotations of HDST data. **(b)** Representative svgs identified by HEARTSVG.

### 7. Application to primary liver cancer data by 10X Visium

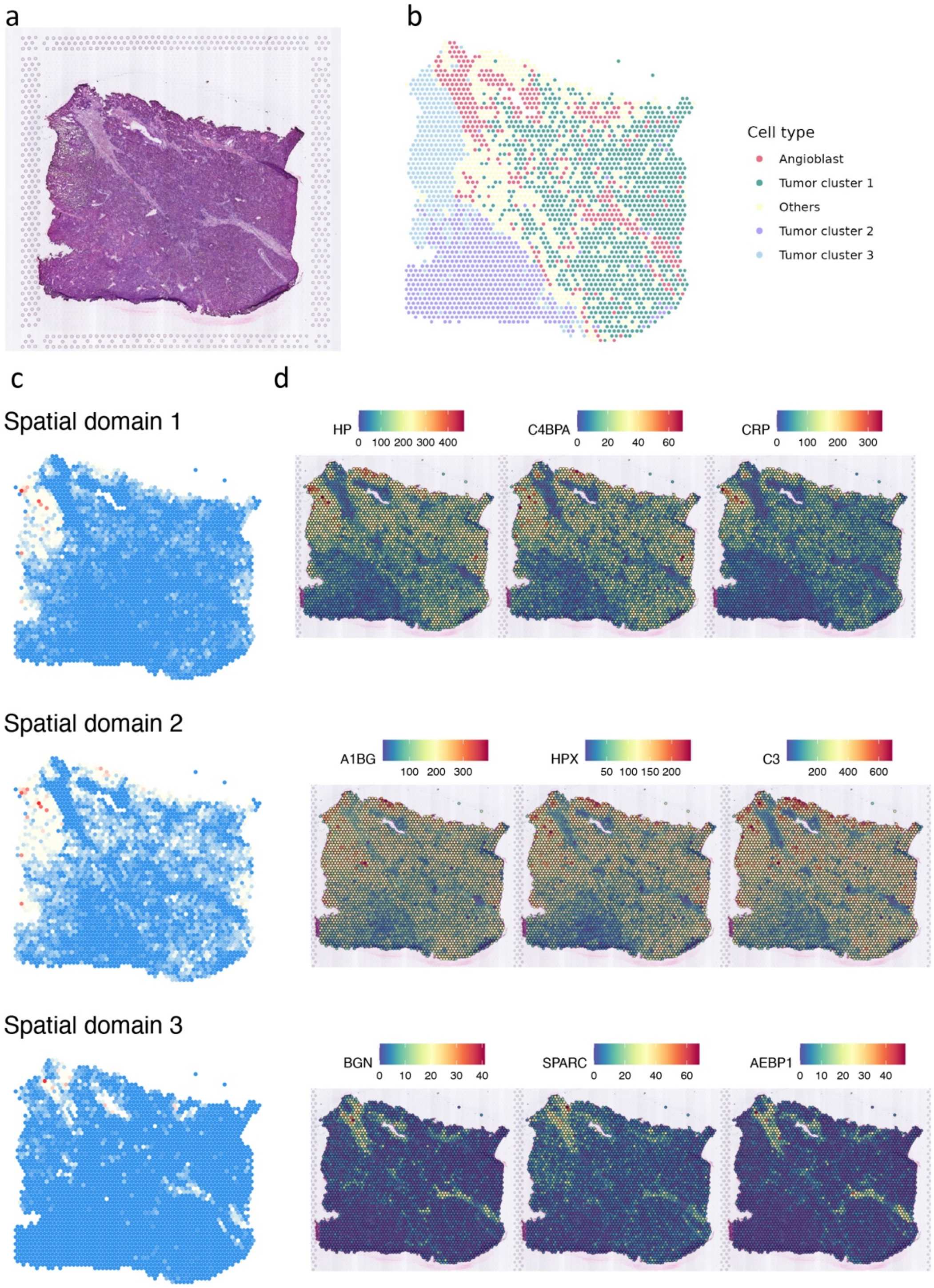


**Figure S24** **(a)** Original hematoxylin and eosin stained (H&E) tissue image. **(b)** Unsupervised spatial clustering results. **(c)** SVGs cluster patterns. **(d)** Representative genes of six SVG clusters.

#### 8. Application to prenal clear cell cancer brain metastasis data by 10X Visium

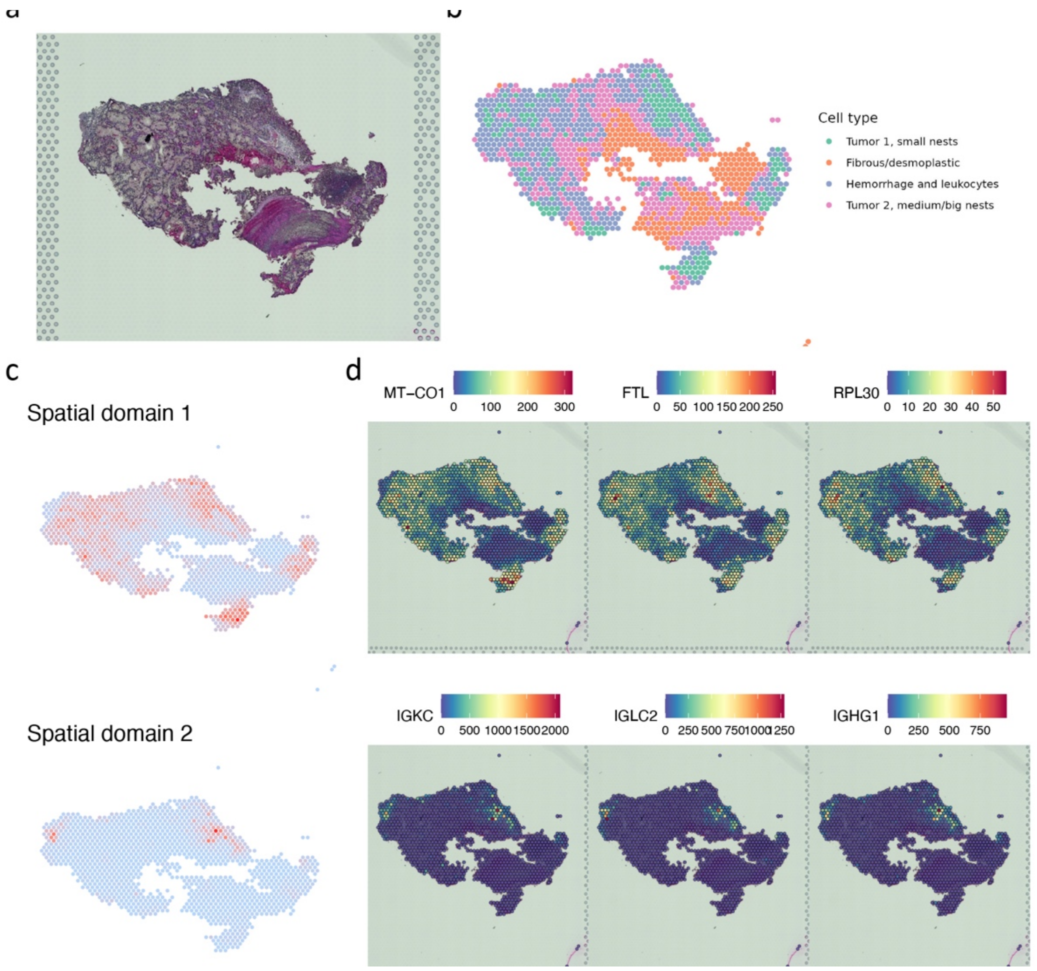


**Figure S25** **(a)** Original hematoxylin and eosin stained (H&E) tissue image. **(b)** Unsupervised spatial clustering results. **(c)** SVGs cluster patterns. **(d)** Representative genes of six SVG clusters.

### 9. Additional analysis

We applied HEARTSVG to analyze three datasets used in the scGCO study, Mouse olfactory bulb data (MOB data) amd Breast cancer data (BC data) generated by Spatial Transcriptomics technology, and Mouse neuron tissue data generated by LCM technology (LCM data).

#### 9.1. Mouse olfactory bulb data (MOB data)

HEARTSVG identified 1610 SVGs in the MOB data. We reproduced scGCO's identification and detected 830 SVGs (in the original paper, it was reported as 796 SVGs).

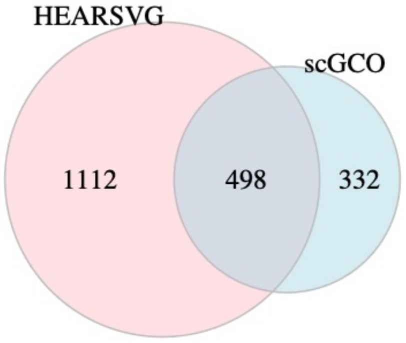


**Figure S26** Venn diagrams of SVGs in the MOB data identified by HEARTSVG, and scGCO.

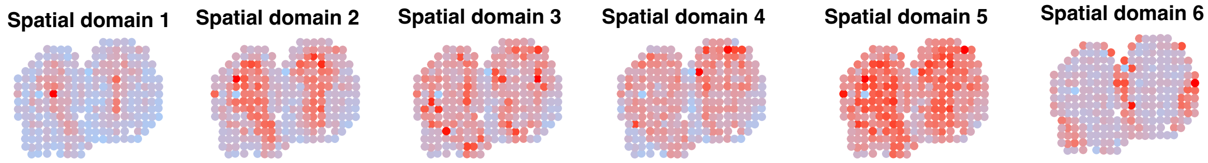


**Figure S27** HEARTSVG predicts spatial domains based on SVGs of the MOB data and graphed the average expression of SVGs in each spatial domain.

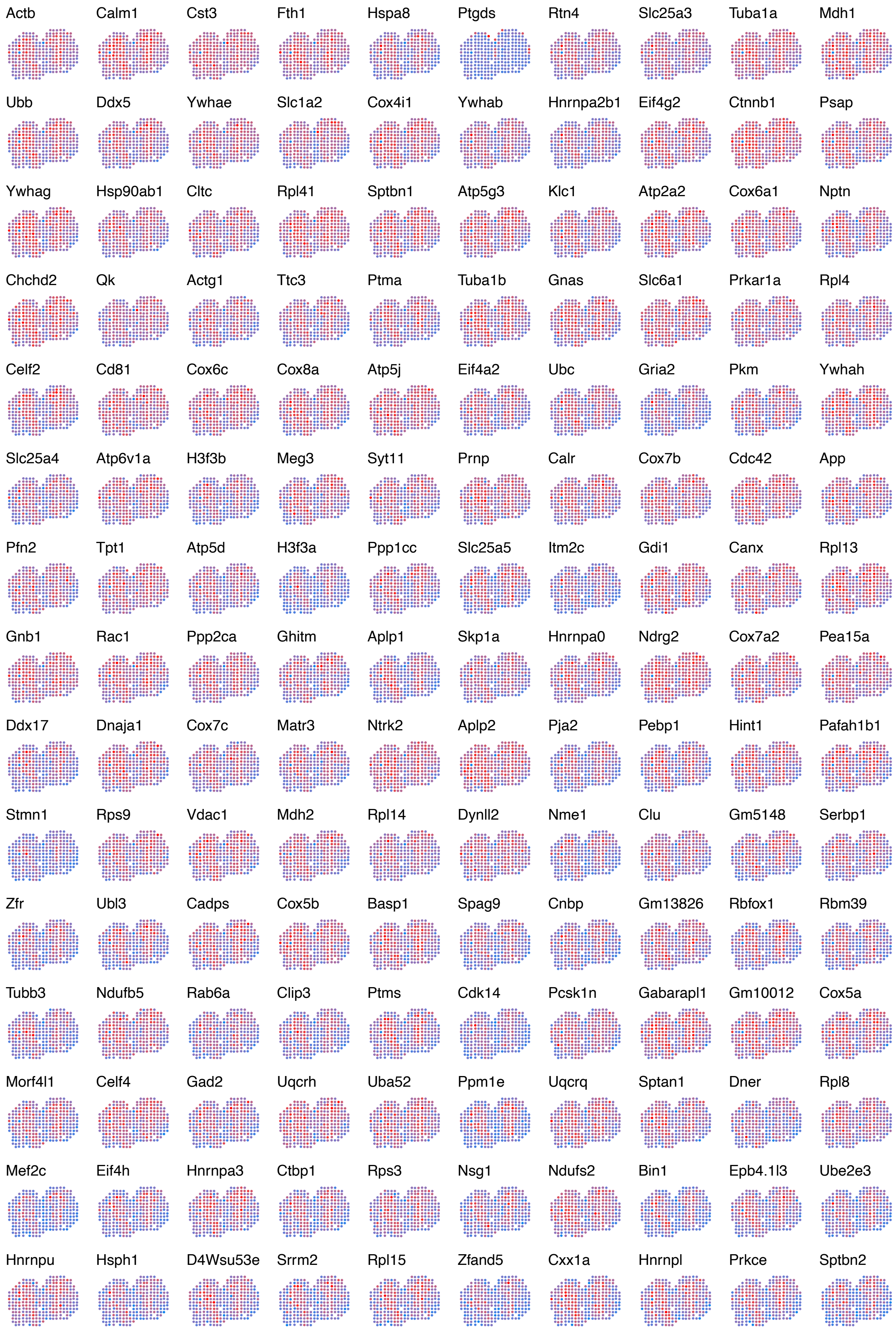


**Figure S28** Top 150 SVGs in the **HEARTSVG-only** SVG list of the MOB data.

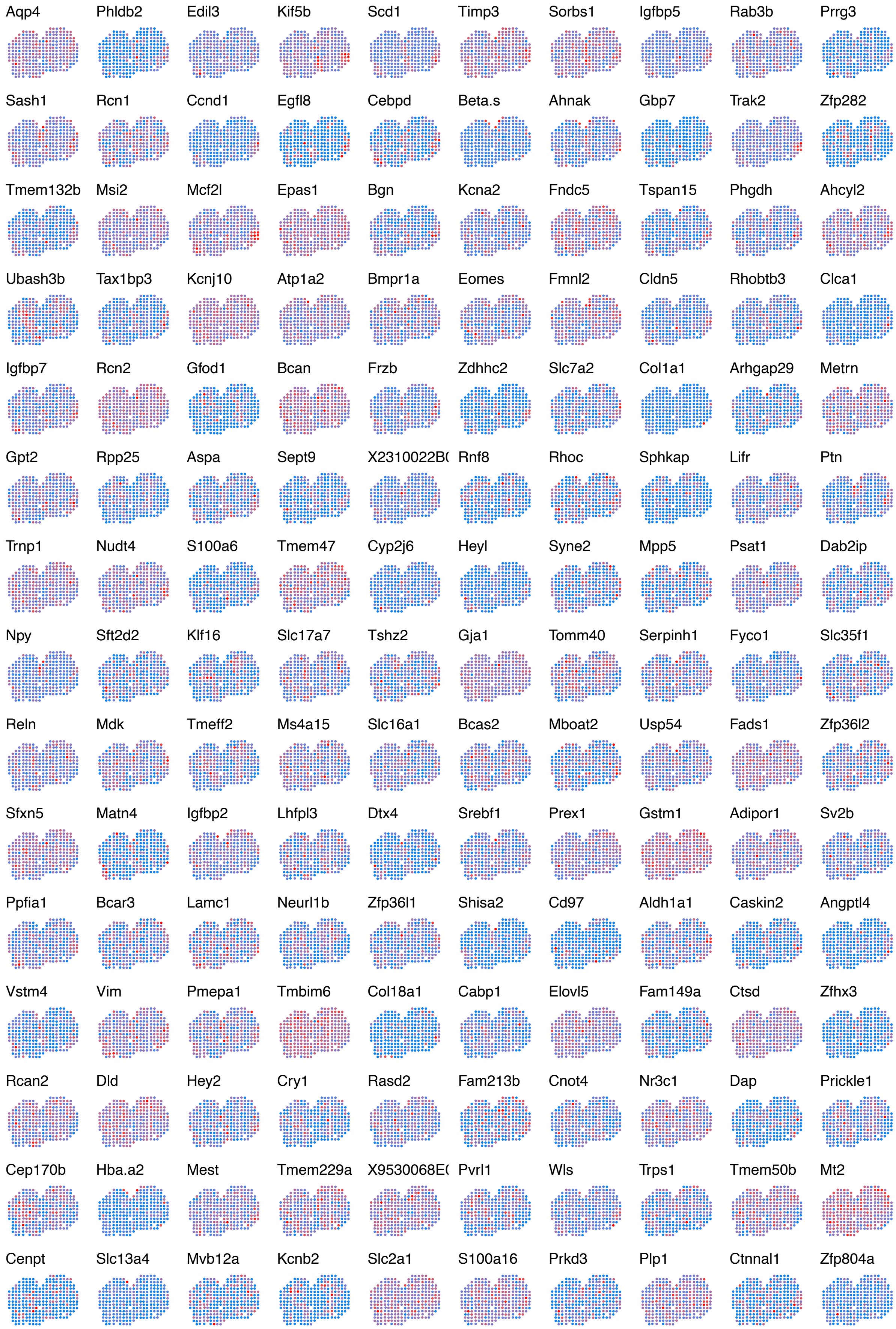


**Figure S29** Top 150 SVGs in the **scGCO-only** SVG list of the MOB data.

##### 9.2. Breast cancer data (BC data)

HEARTSVG identified 287 SVGs in the BC data. We reproduced scGCO's identification and detected 330 SVGs (in the original paper, it was reported as 309 SVGs).

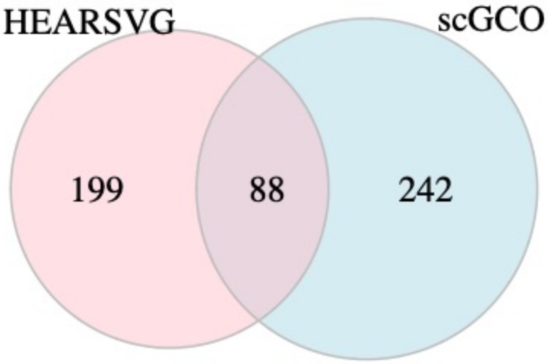


**Figure S30** Venn diagrams of SVGs in the BC data identified by HEARTSVG, and scGCO

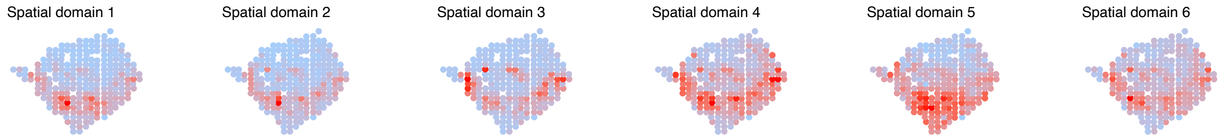


**Figure S31** HEARTSVG predicts spatial domains based on SVGs of the BC data and graphed the average expression of SVGs in each spatial domain.

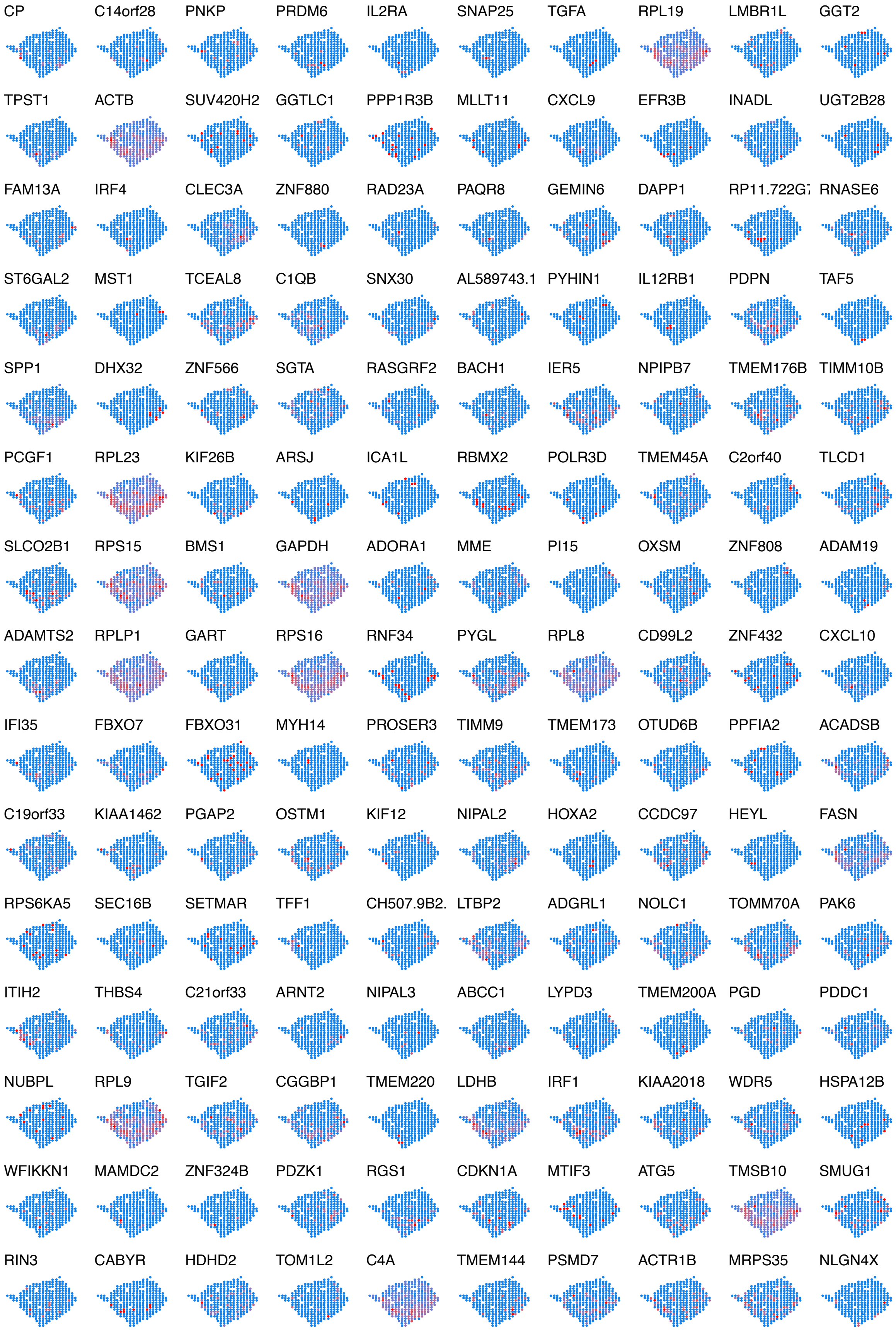


**Figure S32** Top 150 SVGs in the **HEARTSVG-only** SVG list of the BC data.

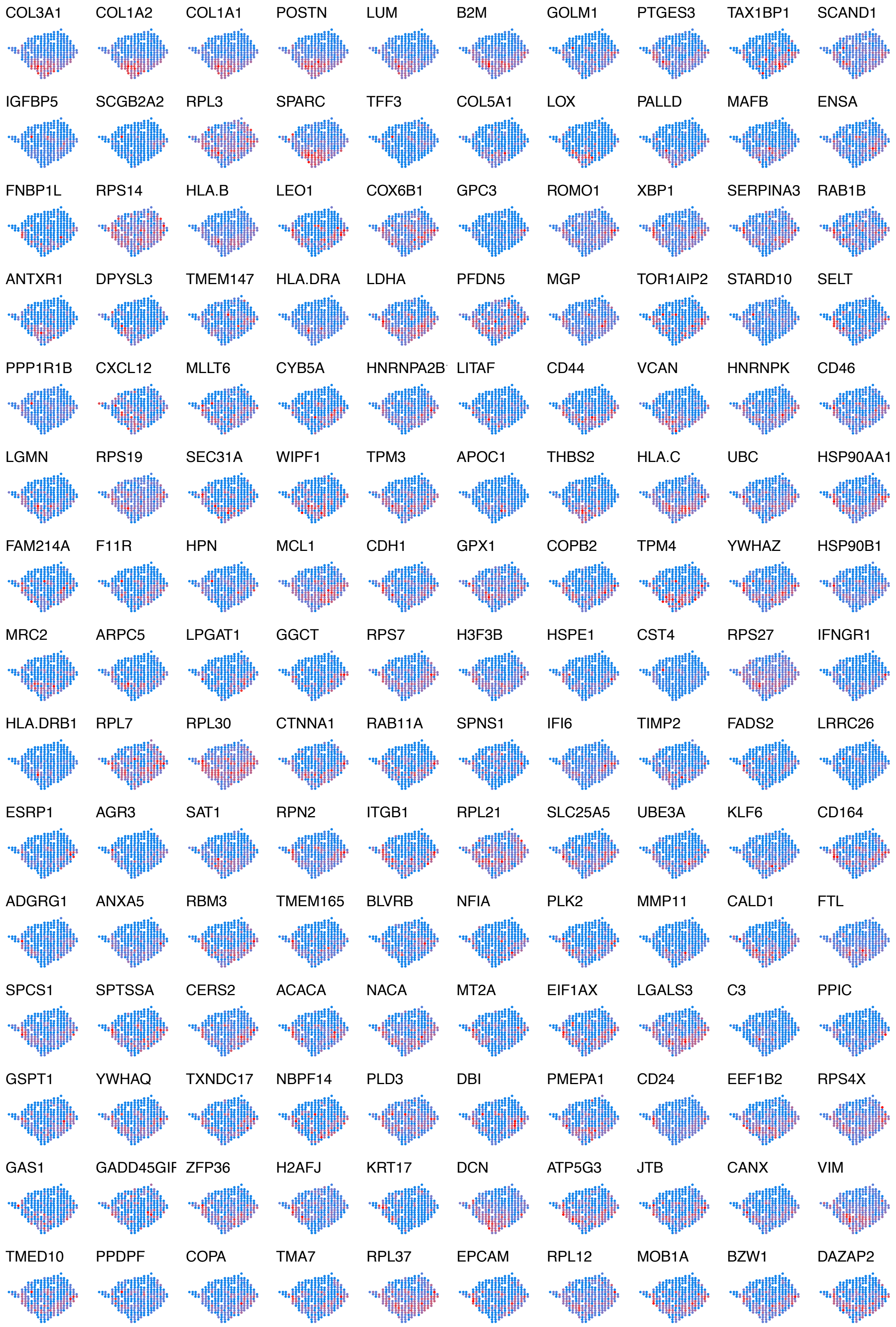


**Figure S33** Top 150 SVGs in the **scGCO-only** SVG list of the BC data.

##### 9.3. Mouse neuron tissue data with LCM technology (LCM data)

HEARTSVG identified 420 SVGs in the LCM data. We reproduced scGCO's identification and detected 754 SVGs (in the original paper, it was reported as 3867 SVGs). In our analysis, HEARTSVG and scGCO did not report spike genes as false positives.

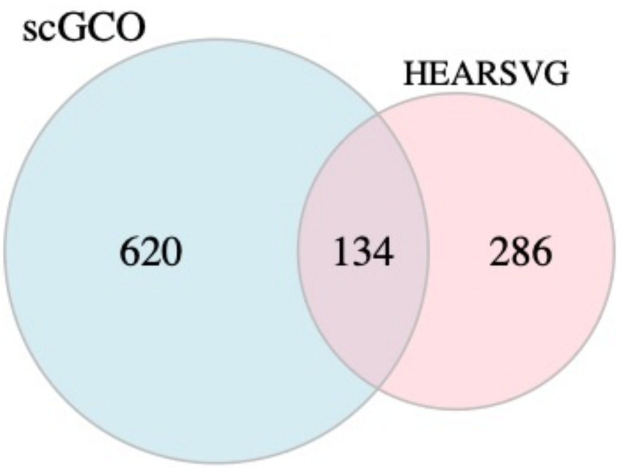


**Figure S34** Venn diagrams of SVGs in the LCM data identified by HEARTSVG, and scGCO

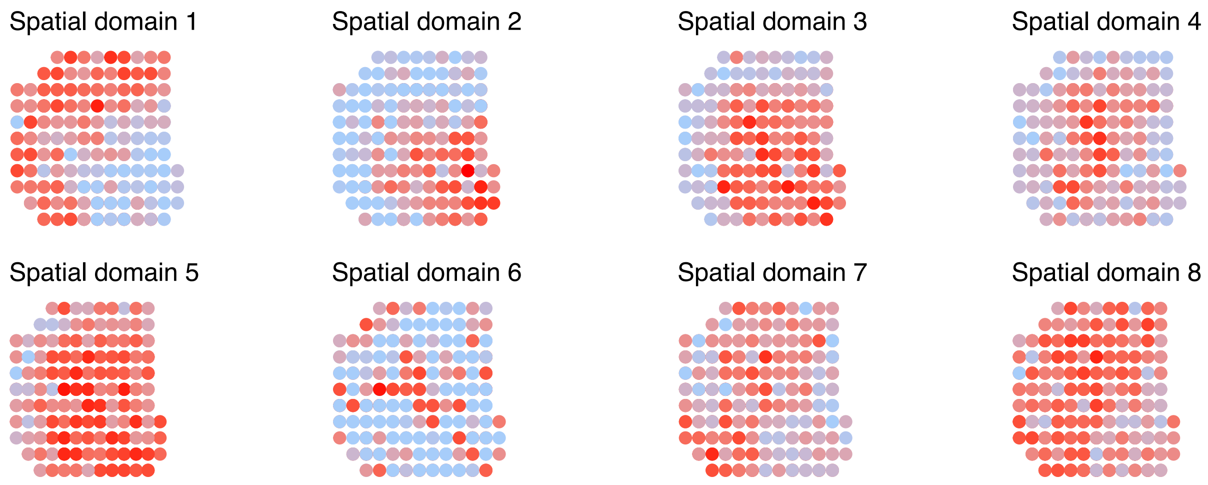


**Figure S35** HEARTSVG predicts spatial domains based on SVGs of the LCM data and graphed the average expression of SVGs in each spatial domain.

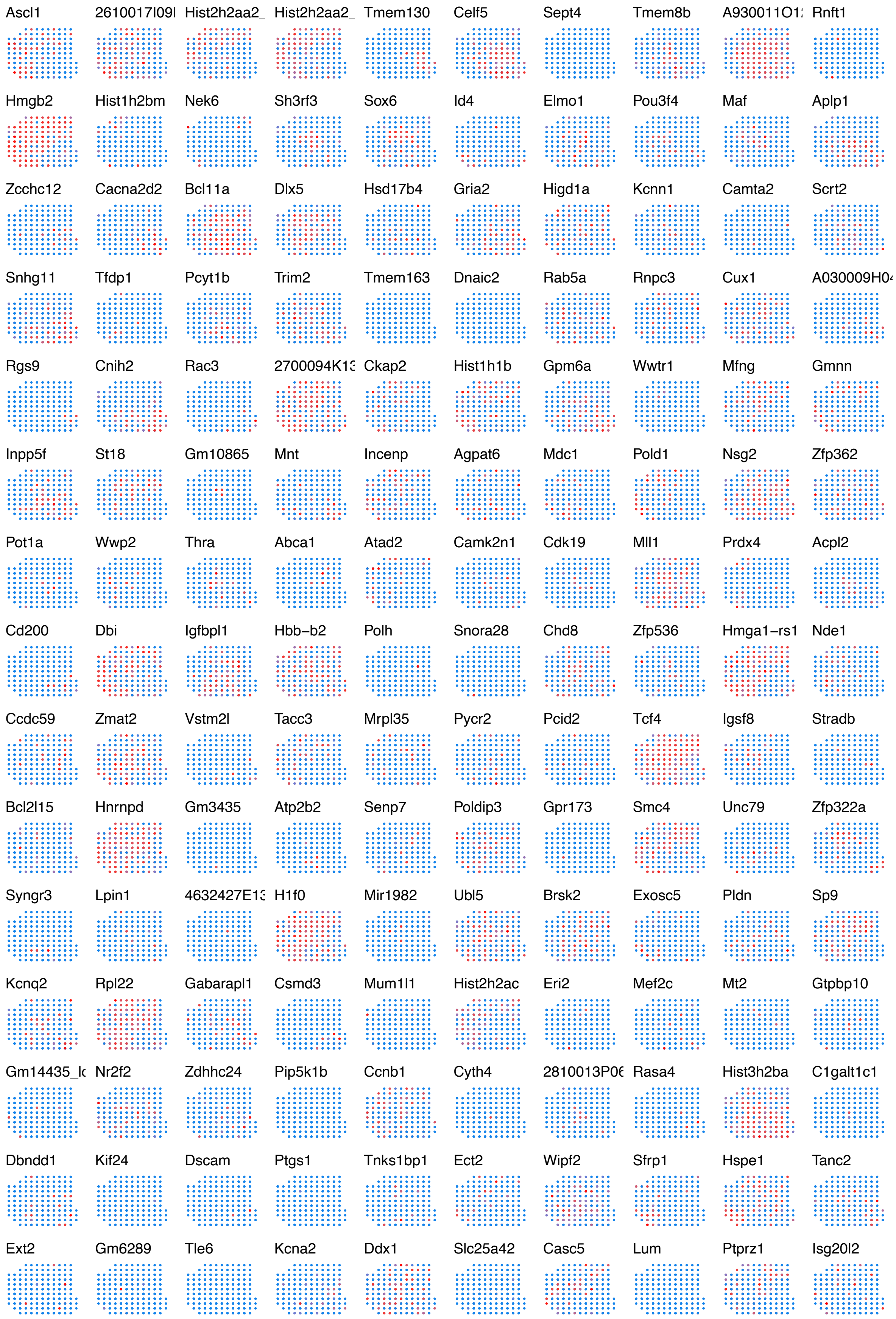


**Figure S36** Top 100 SVGs in the **HEARTSVG-only** SVG list of the LCM data.

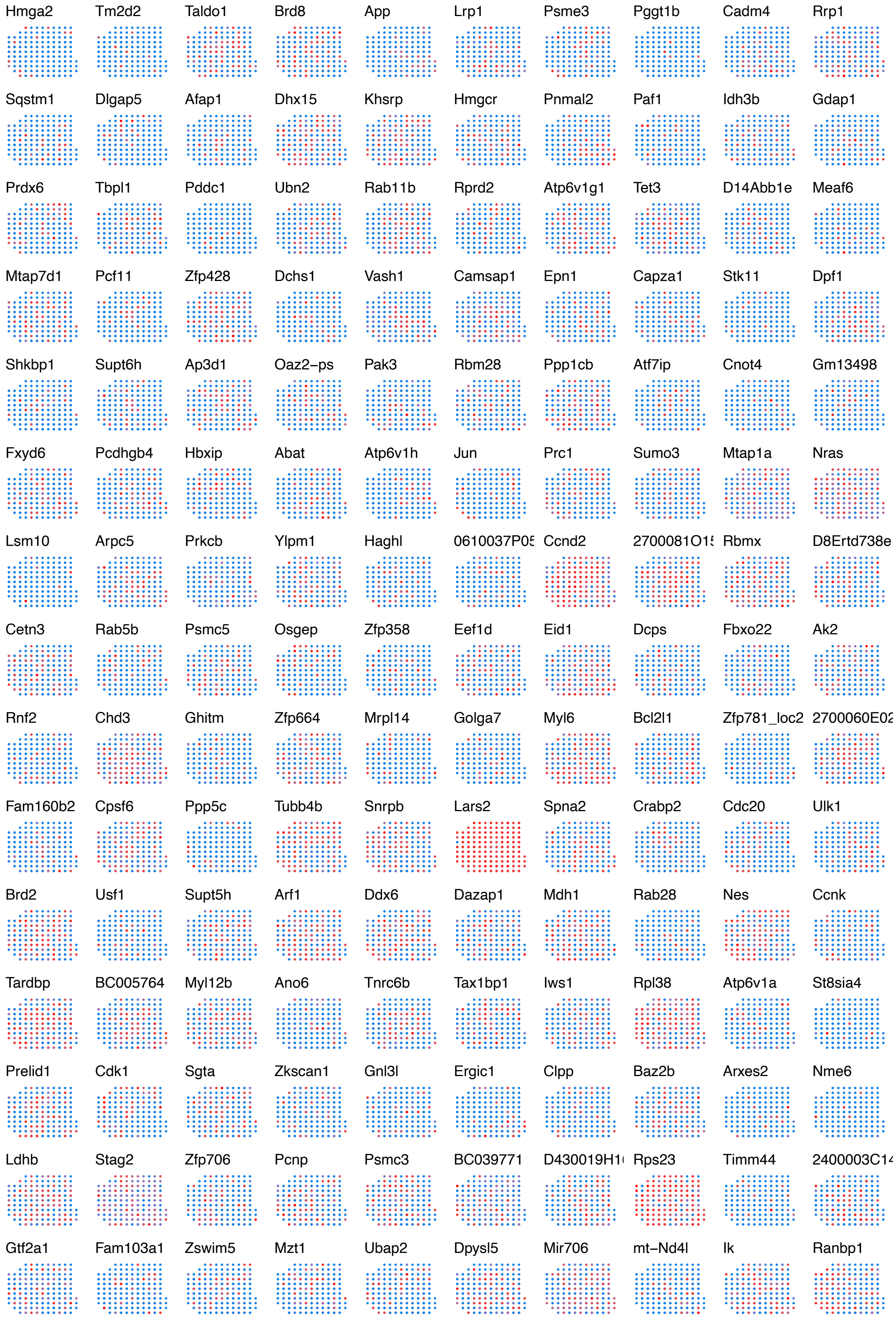


**Figure S37** Top 100 SVGs in the **scGCO-only** SVG list of the LCM data.
